# pI as a Potential Factor Influencing Evolutionary Residue Selection and Structural Stability Among Junctional Adhesion Molecules

**DOI:** 10.1101/2025.10.12.681872

**Authors:** Taner Karagöl, Alper Karagöl

**Affiliations:** Istanbul University Istanbul Medical Faculty, Istanbul, Turkey

**Keywords:** Junctional adhesion molecules, residue conservation, network centrality, pH-based molecular dynamics, evolutionary molecular dynamics

## Abstract

**Objective:** <u>J</u>unctional <u>a</u>dhesion <u>m</u>olecules (JAMs) are a family of conserved proteins involved in immune regulation and cell adhesion. In this study, we investigate the evolutionary and structural dynamics among three paralogs in *Homo sapiens*, which share similar tertiary structures but differ in isoelectric points (pI) (JAM-B: 9.23, JAM-A: 8.09, JAM-C: 7.53).

**Methods:** By integrating residue conservation, partial correlation, network centrality, pathogenicity analyses, and evolutionary molecular dynamics in various pH (6.5-10.5) conditions, we explore how these proteins have functionally and evolutionary diversified.

**Results:** Partial correlation-conservation analysis identified JAM-B functions as an evolutionary hub. Network analyses further highlighted Lys and Cys residues in JAM-B as central evolutionary residues. Negatively charged and hydrophobic residues (Tyr, Val, Asp) were conserved at lower-pI (JAM-C). AlphaMissense profiling revealed that acidic->basic mutations exhibit significantly lower pathogenicity scores, particularly in JAM-A and JAM-B. In dynamics simulations, root-<u>m</u>ean-<u>s</u>quare-<u>d</u>eviation (RMSD) profiles revealed a pI-stability relationship: JAM-B, the highest-pI paralog, remained stable across pH levels, while JAM-A and JAM-C displayed V-shaped pH-dependent deviations (JAM-A at pH 8.0, JAM-C at pH 8.5). Dynamics-aware evolutionary analyses identified key residues combining high evolutionary conservation with pH-sensitive fluctuations: JAM-A at Gln66, JAM-B at Gln36 and Val57, and JAM-C at several basic residues (Lys97, Arg108, Arg123, Arg191).

**Conclusion:** Together, these results demonstrate that pI is influencing evolutionary residue selection and pH-dependent structural dynamics. Our integrated evolutionary-dynamics framework provides mechanistic insight into paralog diversification and offering a foundation for targeted mutagenesis or therapeutic modulation of pH-sensitive adhesion processes.

## Introduction

<u>J</u>unctional <u>a</u>dhesion <u>m</u>olecules (JAMs) are members of an immunoglobulins subfamily localize to cell-cell contacts and are specifically found at tight junctions, expressed by platelets, leukocytes, endothelial and epithelial cells (1, 2). The recent understanding of JAMs’ functions revealed a wider role in diverse developmental processes, including the barrier-forming epithelia, the development of the peripheral nervous system and brain, and spermiogenesis (2, 3). These roles affect various diseases/disorders, including neurodevelopmental diseases, inflammatory and cardiovascular diseases (and atherosclerosis), reproductive diseases, and cancers (2, 4, 5).

JAMs engage in homophilic and heterophilic interactions with the leukocyte integrins, helping cell-cell and cell-<u>e</u>xtra<u>c</u>ellular <u>m</u>atrix (ECM) adhesion (1, 2). Regulation of cis-dimerization by heterophilic interactions, occurring either in cis or in trans (via both homophilic and heterophilic contacts), is emerging as a central mechanism in controlling JAM signaling (4). By integrating JAMs into novel protein design strategies, it can be possible to engineer cell adhesion mechanisms that including homotypic interactions, heterotypic interactions, or both (6). JAMs have also been shown to activate several intracellular signaling pathways, including NFκB, p44/42 MAPK and phosphatidylinositol 3-kinase (3). Although less studied, another aspect of JAMs is their higher pI. They are structurally, evolutionary similar and share close molecular weights (JAM-A: 32,583.14 Da; JAM-B: 33,206.89 Da; JAM-C: 35,019.95 Da), but differ in isoelectric points (pI) (JAM-B (9.23) > JAM-A (8.09) > JAM-C (7.53)).

A protein’s isoelectric point (pI) corresponds to the pH level at which the molecule carries no overall charge, below this level, proteins are positively (+) charged, above this level, they acquire a net negative (-) charge (7, 8). The protein pI varies greatly from about 4 to 12, this diversity of pI values brings the question on the cause of this variation. Protein isoelectric point greatly affects protein functions in different environmental conditions, and strongly linked to subcellular-localization, as local pH and membrane-charge influence localization-specific pI patterns (8). The pI values have also long been used to distinguish proteins in methods such as protein separation and protein crystallization (7). One of our recent studies also indicated that pI differences in same protein classes can be evolutionary significant, such as synaptogyrins in synaptic membrane (9). However, this pI difference is unlikely to be unidirectional in JAMs since all of them are higher pI proteins, unlike synaptogyrins. Additionally, pH-dependent stability is of particular interest given JAMs’ surface-exposed nature within diverse physiological environments (8). In this case, evolutionary analysis and variational profiling could have a strong potential to explain the functional implications of these chemically distinct forms (10).

Mutational clustering of specific amino acids can be inspected through a statistical analysis of homologous sequences and conservation. This methodology is previously reported for various proteins and produced significant functional insights (9, 11, 12) and our recent large-scale study (13) on alpha-helical proteins showed evolutionary effects on residues and had an impact on understand water soluble protein design approach - structurally similar but chemically distinct proteins, namely QTY-code (14, 15). Additionally, network centrality analysis can be further used to assess the structural communication roles of individual residues (16). To complement evolutionary and structural network approaches, we employed AlphaMissense (17) to evaluate the potential pathogenicity of amino acid substitutions. This deep-learning model predicts whether single-point mutations are likely to be benign or disease-associated, based on sequence context, structural features, and evolutionary constraints (17).

In addition, we conducted <u>m</u>olecular <u>d</u>ynamics (MD) simulations in 9 pH steps (6.5-10.5) on three JAM paralogs, providing results of their stability and structural congruence in various pH conditions. The fundamental concept of an MD simulation involves calculating the interatomic forces within a biomolecular system, such as a protein in aqueous solution, thereby enabling the study of its dynamic behavior (18). Since the three JAM human paralogs differ in isoelectric points, systematic sampling across a physiologically relevant pH range can provide insights into how electrostatic balance shapes their conformational dynamics (19). MD simulations are invaluable for elucidating protein and biomolecular mechanisms, structural disease bases, and guiding the design of peptides and proteins (18, 20).

Beyond capturing global stability, our integration of MD-derived metrics with evolutionary analyses provides a new framework for “dynamics-aware residue-wise evolutionary analysis”. Traditional conservation scoring identifies residues critical for structural or functional stability across homologs (21), yet does not account for their dynamic plasticity. By combining residue conservation with simulation-derived flexibility (root-<u>m</u>ean-<u>s</u>quare fluctuation, RMSF), <u>s</u>olvent <u>a</u>ccessibility <u>s</u>urface <u>a</u>rea (SASA), and their pH-dependent variation, it becomes possible to detect “key dynamic-conserved residues” (positions that are simultaneously evolutionary constrained and dynamically sensitive to environment).

Dynamic-conserved residues can be strong candidates for functional hotspots, structural pivots, and determinants of pH-dependent/adaptive behavior. Thus, this integrative framework enables a multi-scale understanding of proteins, linking evolutionary residue conservation with electrostatic modulation and dynamic stability. Our study further embraces the complexity of molecular evolution by including a modular approach. Understanding the structural principles in JAMs provides a framework for designing synthetic cell interactions that can adapt to different cellular compartments, as well as targeted-mutagenesis or therapeutic modulation in various environmental conditions, such as solubility and pH.

## Results and Conclusions

### Physicochemical Properties of JAM Members

The three JAM proteins (JAM-A, JAM-B, and JAM-C or JAM1, JAM2, and JAM3) display differences in their physicochemical characteristics that likely contribute to their distinct evolutionary and functional behaviors. JAM-A and JAM-B consist of 299 and 298 amino acids, respectively. JAM-C is slightly larger, comprising 310 amino acids. Analysis of charged residue distribution (22) revealed that JAM-A and JAM-B contain 28 and 29 negatively charged residues (Asp + Glu), whereas JAM-C contains 39. Conversely, JAM-B and JAM-C each contain 40 positively charged residues (Arg + Lys), in contrast to only 30 in JAM-A. These compositional differences in acidic and basic residues align with variations in the proteins’ isoelectric points (Table 1). Multiple sequence alignment also explained the residue divergence within the JAM family (Figure 1). The three JAM proteins share an average of 32.8% sequence identity, with JAM-A / JAM-B sharing 102 identical residues, JAM-B / JAM-C sharing 110, and JAM-A / JAM-C sharing 99. JAM-A exhibited a more positive GRAVY score compared to JAM-B and JAM-C, indicating lower hydrophilicity (23), though all three proteins still showed negative GRAVY scores overall.

**Table 1.**
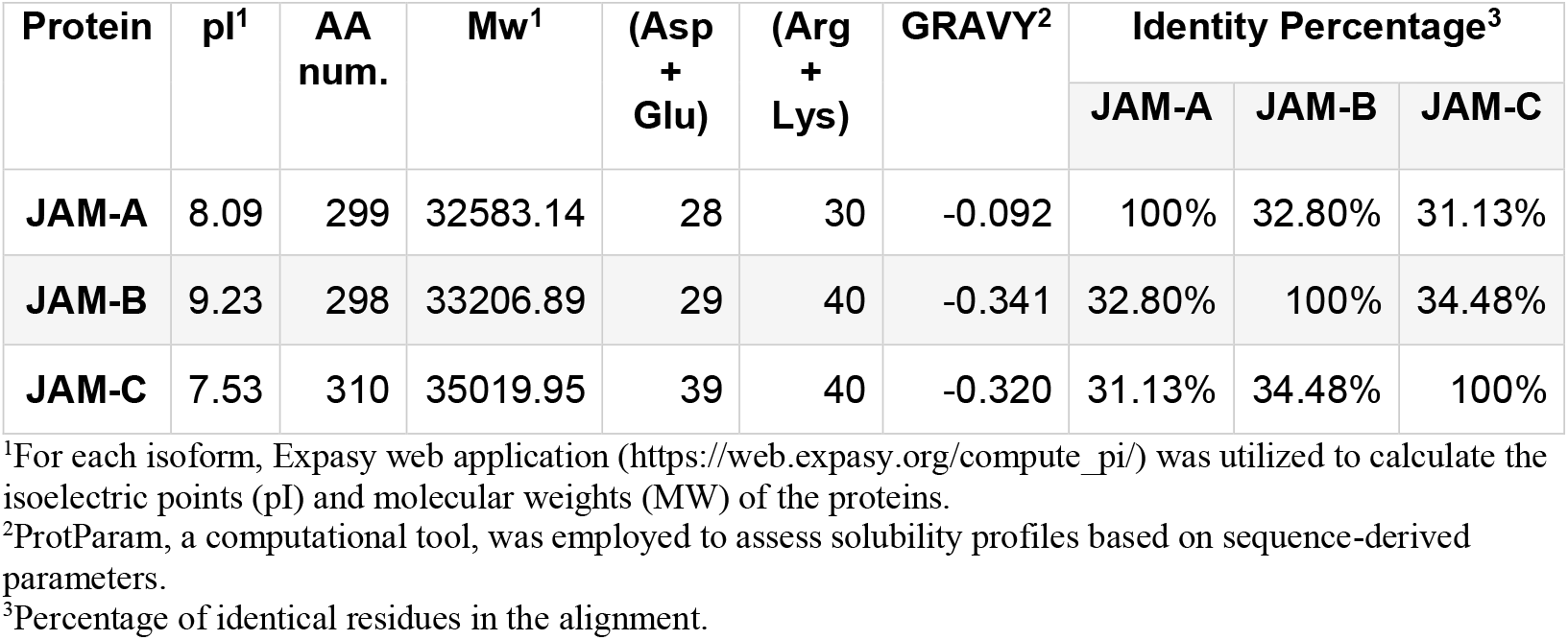
Protein characteristics among Junctional Adhesion Proteins.

| Protein | pI<sup>1</sup> | AA num. | Mw<sup>1</sup> | (Asp + Glu) | (Arg + Lys) | GRAVY<sup>2</sup> | Identity Percentage<sup>3</sup> |  |  |
| --- | --- | --- | --- | --- | --- | --- | --- | --- | --- |
|  |  |  |  |  |  |  | JAM-A | JAM-B | JAM-C |
| **JAM-A** | 8.09 | 299 | 32583.14 | 28 | 30 | -0.092 | 100% | 32.80% | 31.13% |
| **JAM-B** | 9.23 | 298 | 33206.89 | 29 | 40 | -0.341 | 32.80% | 100% | 34.48% |
| **JAM-C** | 7.53 | 310 | 35019.95 | 39 | 40 | -0.320 | 31.13% | 34.48% | 100% |

<sup>1</sup>For each isoform, Expasy web application (https://web.expasy.org/compute_pi/) was utilized to calculate the isoelectric points (pI) and molecular weights (MW) of the proteins.
<sup>2</sup>ProtParam, a computational tool, was employed to assess solubility profiles based on sequence-derived parameters.
<sup>3</sup>Percentage of identical residues in the alignment.

**Figure 1.**
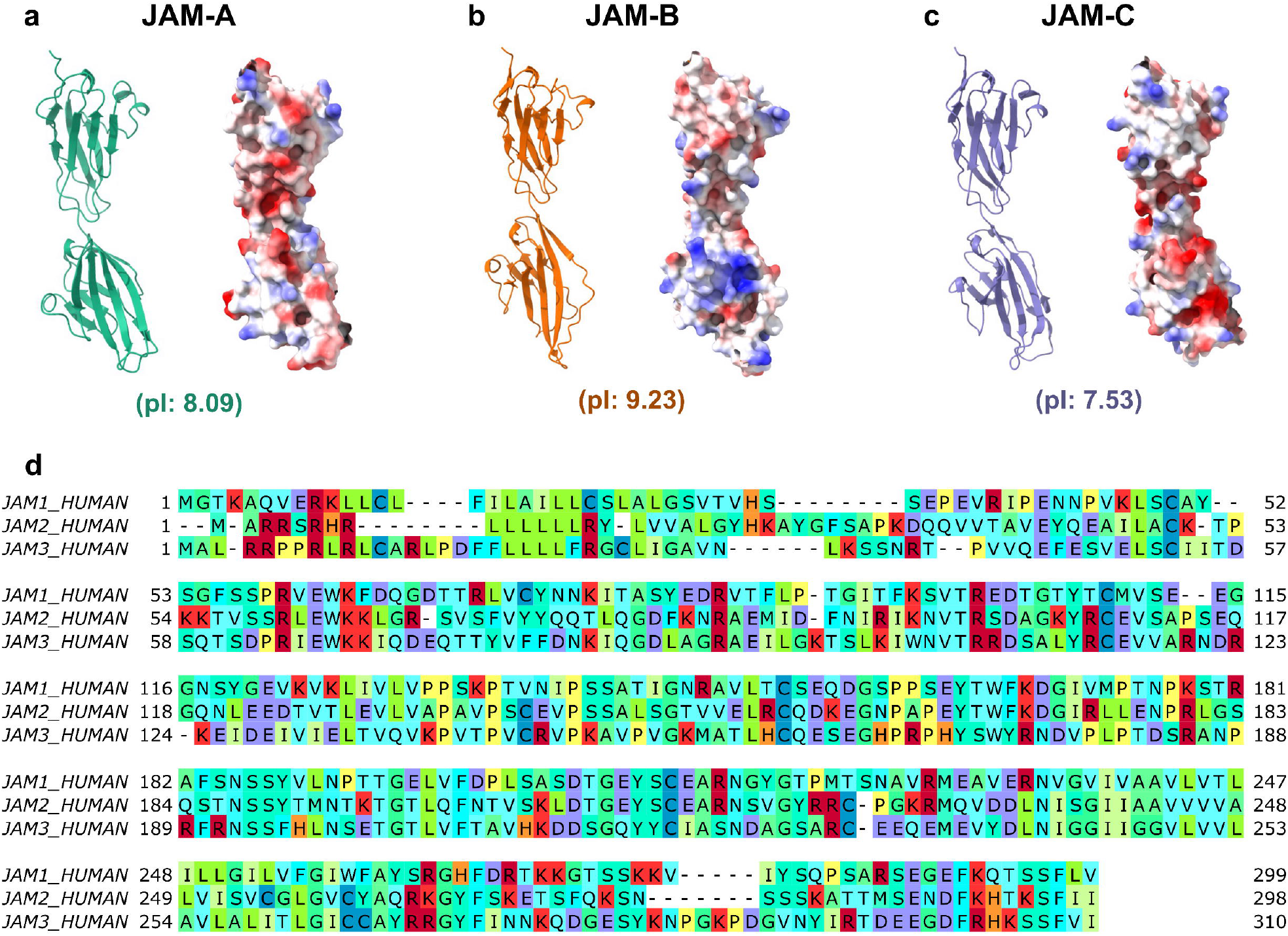
Junction Adhesion Molecules’ structures and multiple sequence alignment. AlphaFold3 predicted structures from UniProt FASTA sequences and electrostatic potential surface coloring of JAM-A (green) **(a)**, JAM-B (orange) **(b)**, and JAM-C (purple) **(c)** in *Homo sapiens*. Multiple sequence alignments **(d)** which were generated with Clustal Omega and visualized with custom color code by isoelectric points (red=highly basic, purple= highly acidic, yellow=neutral).

Comparative amino acid compositions (Supplementary Table 1) showed JAM-C displayed higher proportions of aspartic acid (D), arginine (R), and isoleucine (I), but lower amounts of serine (S). JAM-B, by contrast, was enriched in lysine (K) and glutamine (Q), but contained fewer prolines (P). JAM-A showed a relative enrichment in threonine (T), while being comparatively depleted in arginine (R) and glutamine (Q). The elevated lysine content in JAM-B appears to be the primary contributor to its higher pI, whereas in JAM-A, lower arginine content drives its reduced pI. Another substitution pattern is the increase in histidine (H) in JAM-C (6 residues). Histidine is unique among amino acids because its imidazole side chain has a pKa near physiological pH, allowing it to toggle between protonated and unprotonated states (24).

### Residue-Wise Substitutions

Detailed examination of residue substitutions revealed patterns that directly influence pI, hydrophobicity, and potential functional adaptability. In multiple sequence alignments residue substitutions (Supplementary Table 2), the most common pI-altering substitutions among JAM-A, JAM-B and JAM-C were K<=>T (13 events), T<=>E (12 events), K<=>S (11 events), R<=>S (10 events), R<=>T (8 events), G<=>D (8 events), S<=>E (7 events), and N<=>D (7 events). Within these, the most higher pI change mutations, K<=>E (5 events), R<=>E (4 events), and R<=>D (3 events) were frequently observed, but interestingly, K<=>D substitutions were absent. This suggests majority pI substations were indirect and that serine (S) and threonine (T) may serve as intermediate residues that facilitate mutational transitions between basic (K, R) and acidic residues (E), while glycine (G) and asparagine (N) may act as a mediator in transitions involving aspartic acid (D). K<=>D substitution requires multiple base changes, making it less probable. It has similarly with previously recognized T<=>V substitutions, which have functional significance but also high mutational barriers, can occur mediated with intermediate amino acid such as alanine (13). Extensive form games (25) can be used to represent the sequential nature of these evolutionary changes (13). In this case, the “mediator intermediate” nodes in pI-changing variations could represent a specific decision node in the evolutionary pathway. These specific mediator intermediate nodes found in hydropathy-changing QTY-code substitutions in our recent analysis (13).

**Table 2.** Key dynamic residues of pH change among Junctional Adhesion Molecules.

| Protein | Residue<sup>1</sup> | AA | Residue pI<sup>2</sup> | Residue Conserv.<sup>3</sup> | Avg. RMSF<sup>4</sup> | ΔRMSF<sup>4</sup> | Avg. SASA<sup>5</sup> | ΔSASA<sup>5</sup> |
| --- | --- | --- | --- | --- | --- | --- | --- | --- |
| JAM-A | 66 | Q | 5.65 | 3.01 | 0.17 | 0.07 | 0.82 | 0.34 |
| JAM-B | 36 | Q | 5.65 | 2.08 | 0.21 | 0.08 | 1.53 | 0.35 |
|  | 57 | V | 5.96 | 1.57 | 0.20 | 0.06 | 1.44 | 0.42 |
| JAM-C | 123 | R | 10.76 | 1.09 | 0.22 | 0.12 | 1.94 | 1.08 |
|  | 120 | R | 10.76 | 1.08 | 0.17 | 0.06 | 1.25 | 0.32 |
|  | 97 | K | 9.74 | 1.17 | 0.15 | 0.06 | 1.47 | 0.85 |
|  | 191 | R | 10.76 | 2.05 | 0.22 | 0.05 | 1.59 | 0.71 |
|  | 108 | R | 10.76 | 1.33 | 0.14 | 0.05 | 1.63 | 0.28 |

<sup>1</sup>Residues with high conservation (ConSurf score > 1.0) and elevated ΔRMSF (>0.05 nm) or ΔSASA (>0.3 nm<sup>2</sup>)
<sup>2</sup>pI of residue amino acid at 25 °C.
<sup>3</sup>Consurf evolutionary conservation score of the residue.
<sup>4</sup>The average root-mean-square fluctuation (RMSF) across all frames and steps, and the ΔRMSF (maximum minus minimum across pH).
<sup>5</sup>The average solvent accessibility surface area (SASA) across all frames and steps, and the ΔSASA (maximum minus minimum across pH).

The most frequent overall substitution was isoleucine<=>valine (I<=>V), observed 21 times. This substitution had little impact on either pI or hydrophobicity, as both residues are nonpolar, aliphatic amino acids with similar physicochemical properties (13, 8). The high frequency of such “physicochemically conservative” substitutions suggests that these sites are under stabilizing selection. Some substitutions occurred at residues that are often subject to post-translational modifications, such as serine and lysine. Examples are: K<=>S (11 events) and R<=>S (10 events), and F<=>Y (11 events). Serine is unique in that it is encoded by two completely separate sets of codons (AGY and TCN), which could lead to divergent substitution trends depending on evolutionary pressures. Lysine, on the other hand, is a hotspot for chemical modifications, such as acetylation, methylation, and ubiquitination (26).

In addition, our analysis reveals that the mutations are balanced to preserve protein features such as pI and hydropathy (Supplementary Table 2). For example, in JAM-A, the relative increase in serine (S) is counterbalanced by a decrease in threonine (T), both polar residues and hydroxyl groups, likely ensuring that hydrophilic interactions remain intact. Interestingly, we have previously recognized T<=>S substitutions as less pathogenic than others in alpha-helical membrane proteins (13).

### Evolutionary conservation analysis of residues

The analysis of specific amino acid frequencies in evolutionary conservation data produces insights into the evolutionary dynamics of residues in JAMs (Figure 2) (Supplementary Table 3, 4, and 5). In partial correlation analysis, JAM-B clearly functioned as the evolutionary bridge: removal of JAM-B caused a dramatic decrease in correlation between JAM-A and JAM-C (from mean 0.75 to 0.15) (Supplementary Table 3). Whereas, JAM-A maintained more independent correlation between other JAM members, as excluding JAM-A only made 0.12 correlation difference (0.88 to 0.76), suggesting that JAM-A’s evolutionary integrity is largely independent of the other two. By contrast, JAM-C, in turn, lowest conservation profile without JAM-B, underscoring its dependence on JAM-B for maintaining evolutionary constraints. At first glance, one might expect JAM-A to act as the bridge, given its intermediate pI. However, the pI ranking of the JAM proteins (JAM-B > JAM-A > JAM-C) may also provide a mechanistic explanation for the observed pattern. High-pI JAM-B contains residues that support correlations with lower-pI partners, enabling it to bridge evolutionary constraints across the family. Conversely, the lower-pI JAM-C depends more heavily on JAM-B to maintain correlations, while JAM-A’s intermediate pI allows it to sustain relatively independent evolutionary integrity, rather than serving as a connector.

**Figure 2.**
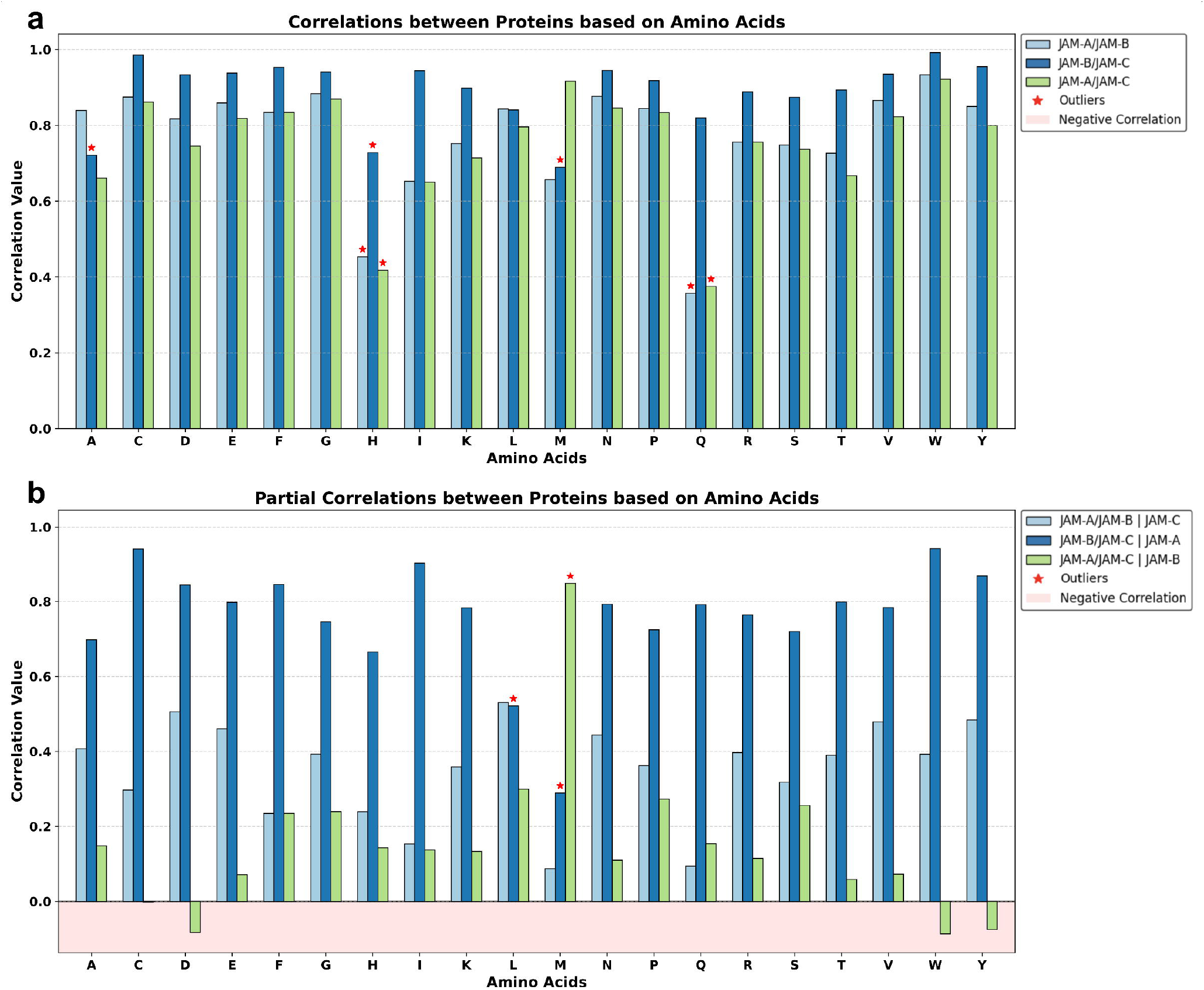
Protein correlation and partial correlation analysis for each amino acid residue-wise conservations in JAMs. The correlation analysis **(a)** between proteins JAM-A and JAM-B (light blue), JAM-B and JAM-C (dark blue), JAM-A and JAM-C (green), for amino acid conservations. In partial correlation analysis **(b)**, removal of JAM-B caused a dramatic decrease in correlation between JAM-A and JAM-C (green). The divergency (outliers) of residues may show specific evolutionary pressures for these residues (star signs on top of columns).

Correlation with pI change of proteins provided additional clarity (Supplementary Table 6). Lysine (K, r = 0.991), methionine (M, r = 0.897), and histidine (H, r = 0.619) correlated positively with pI, being preferentially conserved in higher-pI proteins. Conversely, tyrosine (Y, r = −0.983), valine (V, r = −0.955), phenylalanine (F, r = −0.858), and aspartic acid (D, r = −0.810) correlated negatively with pI, being conserved in lower-pI protein (JAM-C). These patterns suggest that charged residues play evolutionary stabilizing roles in high-pI proteins, while aromatic and acidic residues stabilize lower-pI proteins.

Interestingly, the divergency of residues from other residues may show evolutionary pressures for these residues (outliers in Figure 2). Histidine showed low in all protein correlations comparing other residues. Glutamine displayed a striking divergence in correlations involving JAM-A, showing significantly weaker associations in both JAM-A / JAM-B and JAM-A / JAM-C comparisons. This suggests that Q residues in JAM-A are subject to distinct evolutionary constraints not shared with JAM-B or JAM-C. Given glutamine’s role in stabilizing protein structures (27) through hydrogen bonding while also allowing conformational flexibility, these results imply that glutamine may contribute to JAM-A’s unique functional or structural requirements.

In the other hand, leucine (L) showed significant lower correlation in excluding JAM-A in partial correlation. Leucine’s strong hydrophobic character makes it critical for maintaining protein core packing and transmembrane stability. Leucine’s correlation declines more steeply may suggest that it plays a central role in stabilizing the evolutionary constraints that connect JAM-B and JAM-C through JAM-A.

Methionine exhibited particularly unusual behavior. Its correlations remained relatively stable when JAM-B was excluded, unlike most other residues. This may be explained by methionine’s unique role as the universal start codon in protein synthesis and the fact that the first two amino acids (including starting methionine) are absent in JAM-B in sequence alignments (Figure 1). As a result, JAM-A and JAM-C can maintain methionine-based correlations independently of JAM-B, whereas other residues show sharp declines in conservation once JAM-B is removed.

### Network Centrality Analysis

In consistent with correlation analysis, not surprisingly, centrality measures consistently identified JAM-B residues as the most critical evolutionary nodes. Lysine (high pI aminoacid) in JAM-B (Kb) displayed the highest betweenness centrality, serving as a bridging residue (Figure 3) (Supplementary Table 7). Further explaining our initial hypothesis on higher pI JAM-B being evolutionary bridge. Cysteine (Cb and Cc) also ranked highly across degree and betweenness metrics, likely due to its involvement in structural constraints such as disulfide bonds. Together, these findings reinforce JAM-B’s role as the primary evolutionary bridging protein in the JAM family.

**Figure 3.**
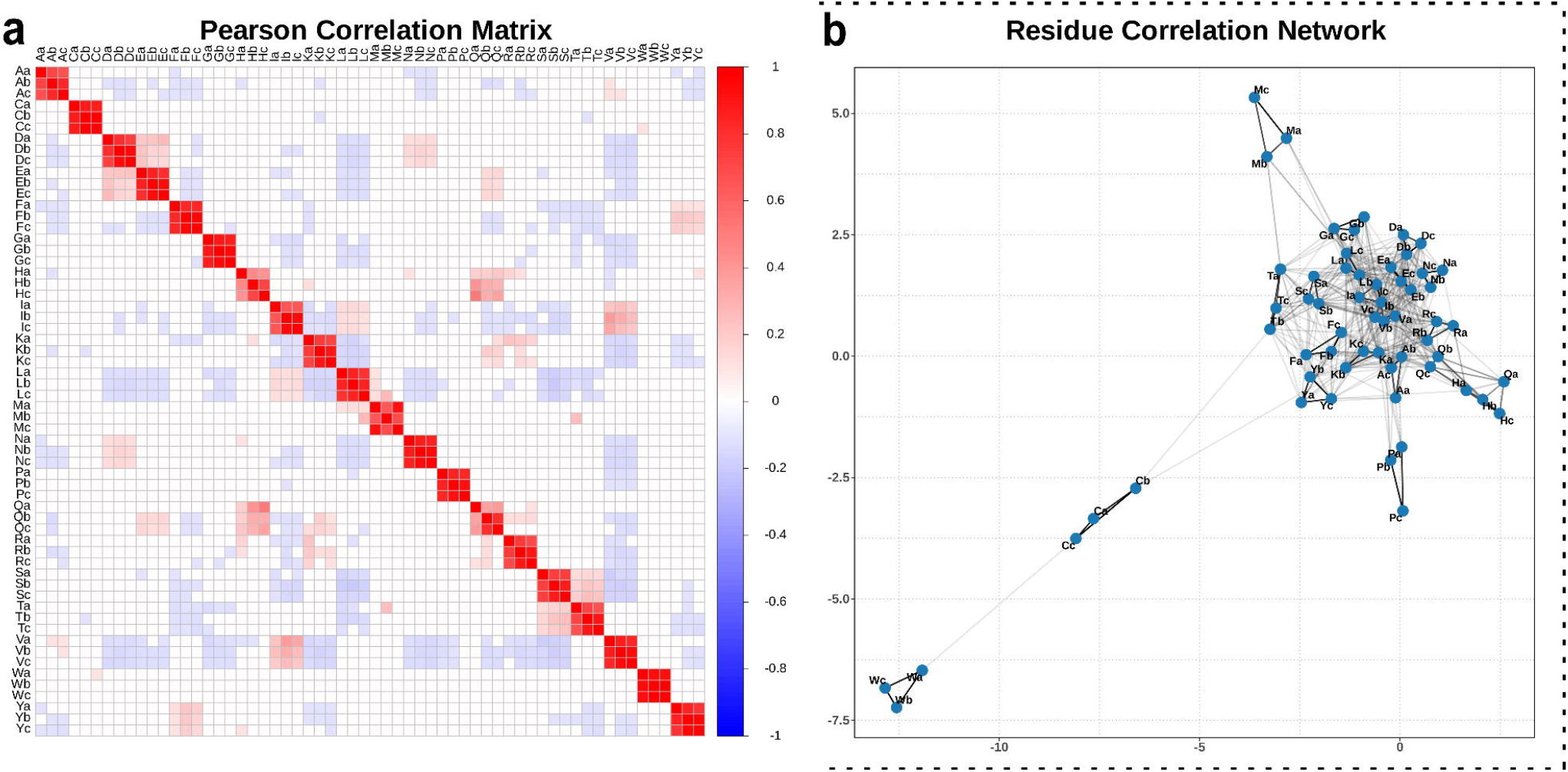
Pearson correlation matrix and residue correlation network for aminoacids in JAMs. Correlation scores for each amino acid in the JAM proteins (uppercase letters denote amino acids, while lowercase letters indicate proteins: *Xa* = JAM-A, *Xb* = JAM-B, *Xc* = JAM-C, where *X* represents the amino acid single letter code) were converted into a matrix representation **(a)**, which served as the basis for network construction (correlation coefficient threshold = 0.1) **(b)**. In the resulting networks, residues were represented as nodes (blue dots), and the correlations between them were represented as weighted edges.

In contrast, JAM-A residues demonstrated high degree centrality but low betweenness, reflecting strong connectivity without acting as evolutionary bridges. JAM-C residues overall showed lower centrality scores, supporting the conclusion that JAM-C is structurally flexible and evolutionarily variable (Supplementary Table 7).

### Mutational Pathogenicity Analyses

Analysis of predicted pathogenicity scores using AlphaMissense (17) further distinguished the three JAM proteins (Supplementary Table 8) (Figure 4). Across all mutations, JAM-C exhibited the highest mean pathogenicity score (0.5310), followed by JAM-A (0.5100) and JAM-B (0.4487). From an evolutionary perspective, this gradient suggests that JAM-B has accumulated greater mutational tolerance, consistent with its role as a hub paralog under stabilizing selection, whereas JAM-C appears to be more constrained.

**Figure 4.**
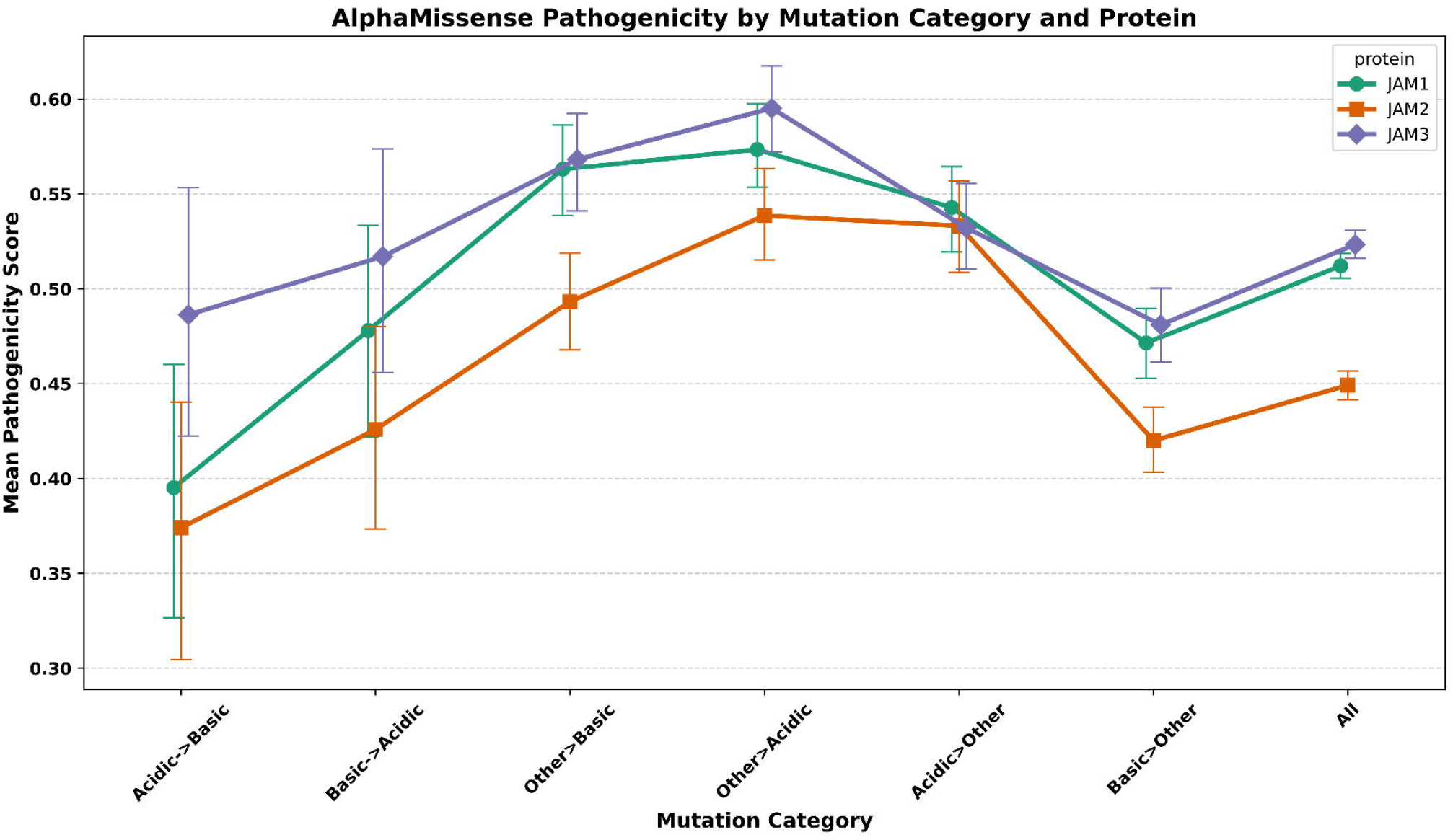
Combined AlphaMissense scores of Junctional Adhesion Proteins by category. Aminoacid pathology scores between mainly acidic (D and E), mainly basic (K and R) and other residue changes between each other, in JAM-A (green), JAM-B (orange), and JAM-C (purple).

More detailed statistical comparisons of mutation classes revealed systematic differences across the JAM proteins and mutation classes (Supplementary Table 9, 10, and 11). Acidic->basic substitutions were associated with significantly lower pathogenicity scores overall (Cohen’s d = −0.27, p < 0.001), with stronger effects in JAM-A (Cohen’s d = −0.36) and JAM-B (Cohen’s d = −0.30) compared to JAM-C (Cohen’s d = −0.20) (Supplementary Table 9). Basic->acidic substitutions showed a weaker and less consistent effect, with small decreases in mean pathogenicity scores that did not reach significance in individual proteins (Cohen’s d = −0.12) (Supplementary Table 10). These results indicate that acidic->basic substitutions are generally more benign across the JAM family, particularly in higher pI proteins JAM-A and JAM-B.

### Molecular Dynamics Simulations in pH change

We performed 3 x 180 ns of molecular dynamics simulations (20 ns x 9 incremental pH steps) to investigate the structural stability of the three JAM proteins under varying pH (6.5 to 10.5) conditions. <u>R</u>oot-<u>m</u>ean-<u>s</u>quare <u>d</u>eviation (RMSD) analysis revealed a clear relationship between pI and structural robustness (Figure 5) (Supplementary Figures 4, 5, and 6). The highest-pI protein, JAM-B, maintained a consistently stable structure across all pH levels, while JAM-A and JAM-C displayed pH-dependent fluctuations, forming a V-shaped RMSD profile depending on pH. Notably, at high pH (≥9), JAM-B retained the lowest RMSD among the three proteins (at pH 10.5 ≈ JAM-B 0.25, JAM-A 0.47, JAM-C 0.35 nm), indicating structural resilience under alkaline conditions. By contrast, JAM-A reached its minimal RMSD at pH 8.0 (≈ 0.19 nm), and JAM-C at pH 8.5 (≈ 0.24 nm), with RMSD increasing at pH levels above or below these optima (Figure 5a).

**Figure 5.**
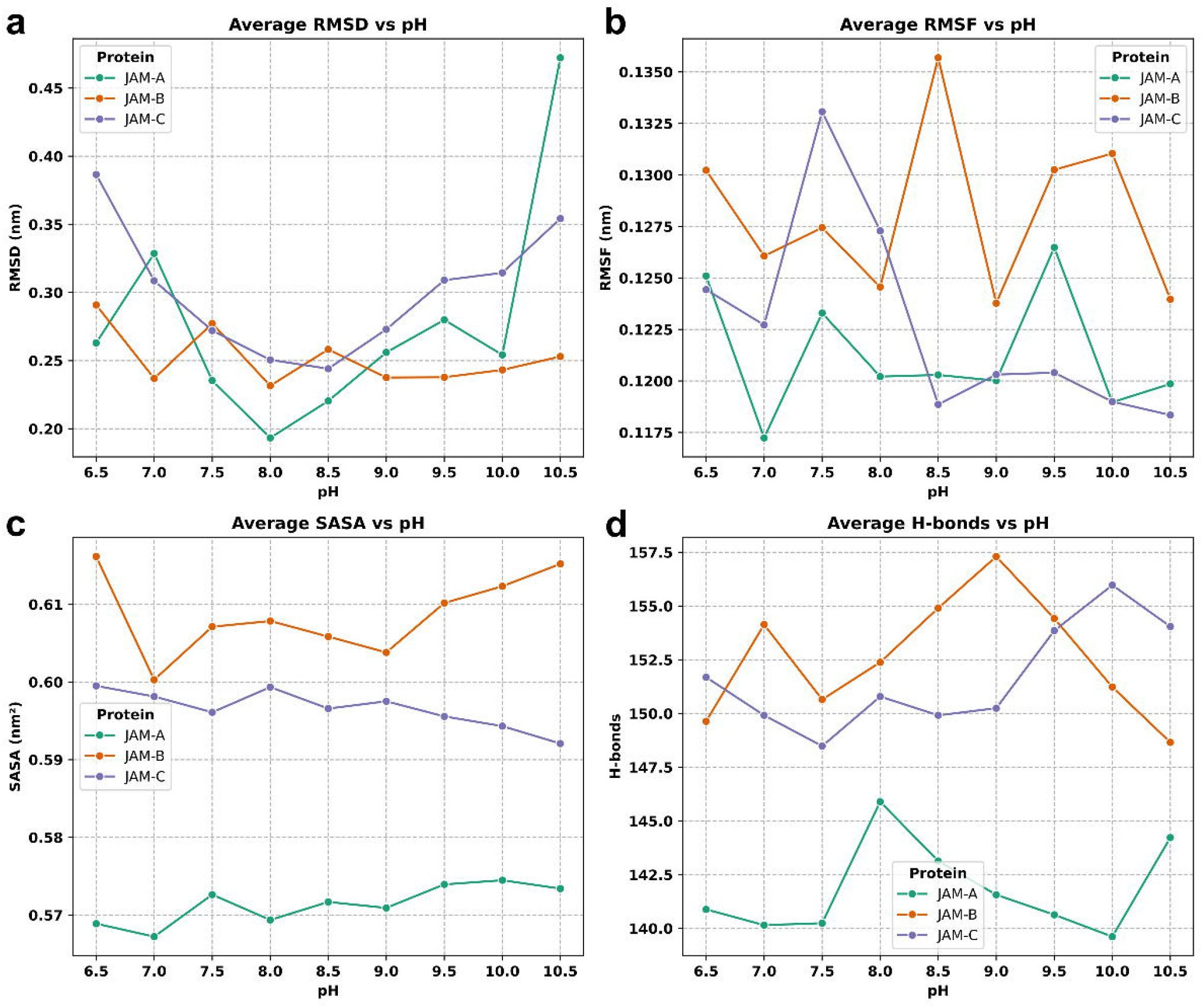
Structural analysis of molecular dynamics simulations of JAMs in pH change. <u>R</u>oot-<u>m</u>ean-<u>s</u>quare <u>d</u>eviation (RMSD) profiles **(a)** show JAM-A and JAM-C displayed pH-dependent fluctuations, forming a V-shaped as pH increased. JAM-A reached its minimal RMSD at pH 8.0 (0.19), and JAM-C at pH 8.5 (0.24). In root-mean-square fluctuation (RMSF) **(b)** analysis, JAM-C showed a steep decrease in RMSF at pH ≥ 8.5. <u>S</u>olvent-<u>a</u>ccessible <u>s</u>urface <u>a</u>rea (SASA) **(c)** and hydrogen-bonds analysis **(d)** showed lower values for JAM-A. Separated figures for each protein can be found in Supplementary Figures 4, 5, and 6.

While RMSD reflected pH-dependent stability graphs, average <u>s</u>olvent-<u>a</u>ccessible <u>s</u>urface <u>a</u>rea (SASA) showed relatively little change across pH levels (Figure 5c). Overall, JAM-B and JAM-C exhibited higher SASA values than JAM-A. <u>R</u>oot-<u>m</u>ean-<u>s</u>quare fluctuation (RMSF) analysis also provided insights: JAM-C (the lowest-pI protein) showed a steep decrease in RMSF at pH ≥ 8.5, indicating reduced local flexibility, whereas JAM-B and JAM-A displayed more oscillatory RMSF patterns across pH (Figure 5b).

High-pI JAM-B maintains both global stability (RMSD) and flexible dynamics across a broad pH range, this may support its role as the evolutionary bridge as more stable structure in various pH conditions. JAM-A and JAM-C show pH-sensitive structural adjustments, consistent with their pI values. Overall, the MD simulations could provide mechanistic support for the observed evolutionary patterns under varying pH conditions.

### Residue-wise Molecular Dynamics Analysis

To explore the interplay between protein dynamics and residue conservation, per-residue root-<u>m</u>ean-<u>s</u>quare fluctuations (RMSF) and <u>s</u>olvent-<u>a</u>ccessible <u>s</u>urface <u>a</u>rea (SASA) were calculated for all simulation steps, then their changes and averages calculated across the trajectories (Figure 6) (Supplementary Table 12, 13 and 14). Our analysis showed that residues in basic residues segments exhibited the higher RMSF and SASA changes (Δ) among JAMs, indicating elevated mobility and solvent exposure among basic residues (Supplementary Figure 7, 8 and 9). Titratable residues, including Glu, Asp, Lys, Arg, and His, showed the higher ΔRMSF and ΔSASA values, consistent with pH-dependent conformational adjustments. These findings were consistent across all three proteins.

**Figure 6.**
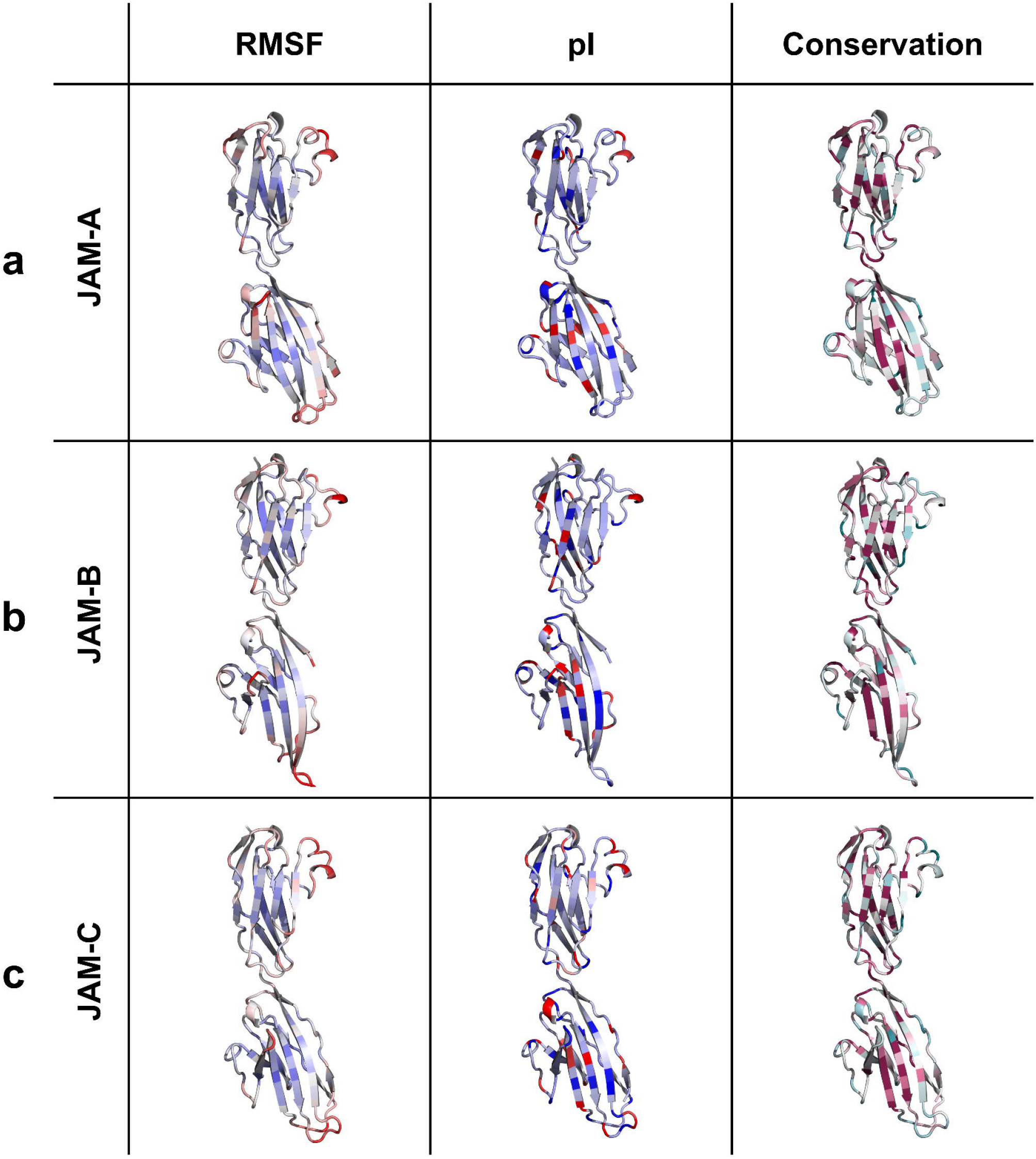
Per-residue molecular dynamics and evolutionary analysis of Junctional Adhesion Molecules. Per-residue average root-<u>m</u>ean-<u>s</u>quare fluctuations (RMSF) (blue-minimum=0, red-maximum=0.25) which were calculated for all simulation steps, then averaged across the trajectories, combined with pI scores (blue-minimum=2.77, red-maximum=10.76), and ConSurf evolutionary conservation scores, for JAM-A **(a)**, JAM-B **(b)**, and JAM-C **(c)**.

### Key Dynamic-Conserved Residues

Integrating evolutionary conservation residue profiles with pH-dependent dynamics revealed a focused set of residues that are both highly conserved (ConSurf score > 1.0) (19) and strongly responsive to pH (ΔRMSF > 0.05 nm or ΔSASA > 0.3 nm^2^). These sites could represent structural “hotspots” where evolutionary constraints preserve sequence identity while allowing conformational adaptability to changing electrostatic environments (Table 2).

In JAM-A, a single standout residue emerged: Gln66, which combined a very high conservation score (3.01) with notable dynamic sensitivity (ΔRMSF ≈ 0.07 nm, ΔSASA ≈ 0.34 nm^2^). Such coupling of strong evolutionary residue retention with measurable flexibility can suggest that Gln66 may play a critical role in mediating of local rearrangements under environmental transitions in JAM-A.

JAM-B exhibited two key positions, Gln36 and Val57, each showing moderate to strong conservation (ConSurf scores 2.08 and 1.57, respectively) alongside robust pH-dependent fluctuations. Gln36 displayed ΔRMSF of ∼0.08 nm and ΔSASA of ∼0.35 nm^2^, while Val57 showed slightly lower ΔRMSF (∼0.06 nm) but an even larger solvent-exposure shift (∼0.42 nm^2^). The presence of both polar (Gln36) and hydrophobic (Val57) residues among these sensitive sites indicates that pH-driven dynamics are not limited to ionizable side chains but can extend to neighboring structural elements. Notably, glutamine has also exhibited distinct correlation patterns in JAM-A (Figure 2). Considering its ability to stabilize protein structures (27) via hydrogen bonding, this finding may suggest that glutamine in JAM-A and JAM-B may underlie specific structural or functional features.

On the other hand, the JAM-C protein exhibited the richest network of dynamic-conserved “hotspots”, dominated by basic residues. Among these, Arg123 was particularly striking, with ΔRMSF reaching ∼0.12 nm and ΔSASA ∼1.08 nm^2^, values far above the protein-wide median. Other key positions included Arg191 (ConSurf=2.05, ΔSASA ∼0.71 nm^2^), Lys97 (ConSurf=1.17, ΔSASA ∼0.85 nm^2^), Arg120 (ConSurf=1.08), and Arg108 (ConSurf=1.33), all exhibiting both high conservation and measurable dynamic flexibility. The clustering of these positively charged residues suggests that local fluctuation changes may drive electrostatic repulsion or salt-bridge rearrangements, providing a structural mechanism for pH-responsive conformational shifts (8).

Collectively, these findings demonstrate that evolutionarily conserved residues are not necessarily static. Instead, a set of conserved-sites/residues, particularly glutamine, lysine, and arginine residues, retain the capacity for significant structural fluctuations when environmental pH approaches or crosses their side chain pKa. Such “dynamic-conserved” residues likely represent functional control points or important residues for pH-sensing, interaction modulation or conformational switching, making them promising targets for mutagenesis or biophysical validation in future studies.

### Future Scopes and the Potential Applications

This study reveals that JAM proteins may follow distinct evolutionary strategies to balance robustness, adaptability, and electrostatic regulation. JAM-B serves as an “evolutionary hub” connecting conservation across paralogs, JAM-C exhibits the highest electrostatic sensitivity, reflecting its lower pI and charge-dependent constraints. These differences not only illuminate mechanisms of paralog diversification but also highlight opportunities for translational applications.

The dynamics analyses (and dynamic-conserved residue approach) could provide a crucial foundation for future work, related or unrated to JAMs. The observed pI-stability relationship, with JAM-B maintaining structural stability across pH ranges, and JAM-A and JAM-C displaying V-shaped RMSD deviations, suggests that charge environments strongly dictate conformational flexibility. Moreover, the identification of pH-sensitive evolutionary residues (e.g. JAM-A at Gln66, JAM-B at Gln36/Val57, JAM-C at Lys97/Arg108/Arg123/Arg191) highlights specific evolutionary “key dynamic-conserved residues” where fluctuations intersect with conservation. These residues may represent potential targets for mutagenesis or therapeutic engineering, enabling the rational design of proteins. Also, future work could either mimic evolutionary robustness (as in JAM-A) or exploit central bridging residues (as in JAM-B) to tune synthetic cell adhesions or adaptations to local environments.

Future directions should examine how these dynamics-dependent structural transitions modulate functional outcomes, such as ligand recognition, allosteric regulation, or enzymatic efficiency under varying pH. Given the central role of Lys and Cys residues in JAM-B as evolutionary connectors, targeted manipulation of these amino acids could also help control network stability or adaptability. Additionally, the observed differences in residue conservation between low- and high-pI paralogs have potential in designing adhesion molecules or modulators of immune signaling in varying environmental conditions.

These insights are especially relevant to disease states characterized by pH dysregulation. For example, in cancer (28), heart failure (29) or stroke and depression (30), where acidic or alkaline shifts alter adhesion and signaling, engineered JAM variants or JAM-combined proteins (6) could be applied to stabilize adhesion complexes, enhance therapeutic targeting, or modulate immune responses. More broadly, the integration of evolutionary conservation, network analysis, and molecular dynamics perspectives offers a framework for rational mutagenesis and pH-aware protein design strategies that bridge fundamental biology with therapeutic developments.

## Methods

### Protein multiple sequence alignments and other characteristics

The protein sequence data were retrieved as FASTA formats from UniProt website (https://www.uniprot.org) (IDs: Q9Y624, P57087, Q9BX67) (31). To characterize the investigated proteins, their isoelectric points (pI), molecular weights (MW) scores were calculated through the ExPASy online platform (https://web.expasy.org/compute_pi/) (32, 33, 34). ProtParam tool was used to calculate hydropathy solubility profiles (34). 3D protein structures were generated via AlphaFold3 (35), large N- and C-terminal loops were removed. The proteins colored with ColorBrewer2 color scheme, colorblind safe and print-friendly color schemes for enhancing accessibility of figures (36). The protein surface colored by electrostatic potential with UCSF ChimeraX (version 1.8) (37).

Multiple sequence alignments were generated with Clustal Omega (38) and visualized with custom color code according to isoelectric points using Jalview 2.11.5 (39). Residue-wise substitutions analysis generated with our Evolutionary Statistics Toolkit interactive tool (version 1.2.2) Alignment Substitution Analyzer (AlignSubs version 0.3.1) from Clustal Omega aligned sequences, enabling reproductivity (https://www.tanerkaragol.com/alignment-substitution-analyzer) (40).

### Evolutionary conservation profiles

The Consurf server (https://consurf.tau.ac.il/) was used to generate residue-specific evolutionary conservation profiles (21, 41-44), consistent with our previous analyses (9, 11, 12, 13). Default parameters were applied for homolog search, thresholding, multiple sequence alignment, phylogenetic reconstruction, and conservation scoring. Homologous sequences were retrieved from UniRef90, which clusters UniRef100 entries at 90% or 50% sequence identity using the MMseqs2 algorithm (45). Sequence alignment was performed using MAFFT, while homolog identification utilized HMMER with an E-value cutoff of 0.0001. Only homologs with sequence identities between 35% and 95% were considered. Conservation scores were computed via the Bayesian method, with the amino acid substitution model automatically selected based on the best fit. The resulting residue-wise conservation grades were visualized in PyMOL version 3 (46).

### Residue conservation and correlation analysis

We examined the causal asymmetric relationships in the evolutionary occurrence of individual amino acids across aligned residues. We have conducted protein-based residue-wise correlation and partial correlation analysis for each amino acids conservation estimations. Because our data are not time-continuous, it is difficult to assume that the evolutionary dynamics follow a uniformly aggregated monotonic pattern (25). Furthermore, since the assumption of homoscedasticity may not hold, we employed the generalCorr package in R (47, 48, 49), which implements kernel regression of *x* on *y* (50). This method is particularly suitable for our analysis, as it does not depend on traditional parametric assumptions that our dataset may violate.

For the conservation estimations, ConSurf grades were used. ConSurf uses a phylogenetic approach, considering evolutionary history between sequences (41-44). Statistical calculations were performed using R (The R Foundation for Statistical Computing, Vienna, Austria), version 4.4.0 (https://www.r-project.org/) (51). Plots generated utilizing and matplotlib plotting library with Python (52).

### Network centrality analysis

Correlation and causality-derived matrices were converted into adjacency matrices, which served as the foundation for network construction (correlation coefficient threshold 0.1). In these networks, residues were represented as nodes, while correlations between them were represented as weighted edges. The igraph and corrplot package in R version 4.4.0 (51, 53, 54) was used to construct, analyze, and visualize the resulting residue networks. To characterize the structural and functional roles of residues within the network, several centrality measures were calculated. Degree centrality quantified the number of direct connections each residue had, thereby identifying connected “hub” residues. Weighted degree centrality, extended this measure by summing the correlation values of each residue’s connections. Betweenness centrality identified residues acting as “bridges” by lying on the shortest paths between other residues, marking them as critical nodes for information flow and network integrity. Final network representations were visualized using force-directed layouts in igraph (51, 53).

### AlphaMissense scores and variant pathogenicity analysis

JAM-A, JAM-B, JAM-C protein sequences were manually submitted to the AlphaMissense database via HageLab’s AlphaMissense webpage (https://alphamissense.hegelab.org) (17, 55). We have calculated aminoacid pathology scores between mainly acidic (D and E), mainly basic (K and R) and other residue changes between each other. For each category, variant scores were compiled into tab-delimited (TSV) files. Statistical comparisons between groups were performed using Student’s *t*-test and the Mann-Whitney *U* test (56), and effect sizes were estimated using Cohen’s *d* (57). Python scripts was utilized to handle the data using statsmodels library (58) and generate plots utilizing matplotlib plotting library (52).

### All-atom MD simulations in pH conditions

All <u>m</u>olecular <u>d</u>ynamics (MD) simulations were run using GROMACS 2025.2 (59) on Google Colab, running an Ubuntu environment equipped with L4 GPUs, 106 GB RAM, and 43 GB VRAM. To maximize computational efficiency, simulations were parallelized across multiple cores. The GROMACS source code was recompiled with full core usage, enabling CUDA GPU acceleration (*DGMX_GPU=CUDA*) and OpenMP multithreading (*GMX_OPENMP_MAX_THREADS=128*) optimized for L4 GPUs, following our recent benchmarking protocol with publicly accessible configuration files and scripts, accompanied by a repository, detailed step-by-step instructions (60).

The protein systems were assembled in solution using the CHARMM-GUI web server (59, 61, 62) with predicted AlphaFold3 (35) structures. Each system was solvated in a rectangular box filled with TIP3P water molecules, a minimum buffer distance of 10 Å between the solute and the edges of the periodic simulation box. For each protein, MD simulations were performed at nine pH condition steps - 6.5, 7.0, 7.5, 8.0, 8.5, 9.0, 9.5, 10.0, and 10.5. Each pH-specific simulation was divided into 20 sequential MD steps to facilitate sampling across the simulation range. The all-atom CHARMM36m force field (63) was employed for all simulations. Energy minimization was performed using the steepest descent algorithm until the maximum force converged below 1000 kJ/mol/nm. Subsequently, each system underwent 125ps of equilibration, following the standard CHARMM-GUI equilibration protocol (61, 62, 63), in-line with our previous studies (9, 20, 64). After the NVT (constant volume) and NPT (constant pressure) equilibration, a 3 x (for three proteins) 180 (9 x 20) ns production MD simulation was started with timestamps for every 0.5ns, in pH steps starting from 6.5 to 10.5. The temperature was maintained at 303.15 K, and the pressure control at 1 bar that was achieved using the stochastic cell-rescaling barostat and isotropic coupling. The <u>P</u>article <u>M</u>esh <u>E</u>wald (PME) method used for calculations of long-range electrostatic interactions (with cutoffs of 1.2 nm applied to both Coulomb and van der Waals interactions).

### Dynamics-aware evolutionary analysis

For each trajectory, root-<u>m</u>ean-<u>s</u>quare fluctuation (RMSF) per residue was calculated using gmx rmsf, using the protein backbone as the reference. The average RMSF across all frames and steps was determined for each residue, and the ΔRMSF (maximum - minimum across pH) was computed to quantify pH-dependent flexibility changes. Per-residue <u>s</u>olvent-<u>a</u>ccessible <u>s</u>urface <u>a</u>rea (SASA) was computed using gmx sasa, with total and residue wise contributions recorded. Average SASA values and ΔSASA across pH were similarly calculated to assess solvent exposure changes. Hydrogen bonds (H-bonds) were calculated using gmx hbond to determine the number of intramolecular H-bonds per frame.

Average RMSF and SASA per residue were combined with ConSurf scores (41-44) to identify residues that are both conserved and dynamically responsive. Residues with high conservation (ConSurf score > 1.0) and elevated ΔRMSF (>0.05 nm) or ΔSASA (>0.3 nm^2^) were considered key dynamic-conserved residues. All data processing and calculations were performed using Python (v3.12). Visualization of residue-specific metrics and dynamics trends was performed using xmgrace (https://plasma-gate.weizmann.ac.il/Grace/) and custom Python scripts to generate graphs of RMSF and SASA compared with conservation and pI, using matplotlib (52). Structures were visualized using PyMOL and custom Python scripts, with residue-wise pI, RMSF, and conservation represented as B-factor coloring (46).

## Supplementary information (SI)

Supplementary Information.pdf

Supplementary data file featuring graphical results from the molecular dynamics simulations.

## Ethics approval

Ethics approval was not required for this computational study as it did not involve animal subjects, human participants, and identifiable data.

## Consent to participate

Not applicable. This computational study did not involve human participants.

## Consent for publication

Not applicable. This computational study did not involve human participants.

## Availability of data and materials

Conservation scores for these proteins are available at the ConSurf Database (https://consurf.tau.ac.il/), while individual AlphaMissense scores can be found at HageLab’s AlphaMissense webpage (https://alphamissense.hegelab.org). For more detailed information on the statistical analyses, input files and detailed outputs, and codes to regenerate analyses and graphs, please visit the website: https://github.com/karagol-taner/pI_evolution_JAMs. Requests for additional materials should be addressed to the corresponding authors Taner Karagöl or Alper Karagöl, and will be fulfilled upon reasonable request.

## Author contributions

Both authors (T.K. and A.K.) contribute equally to this study. All authors have read and agreed to the published version of the manuscript.

## References

1) Weber C, Fraemohs L, Dejana E. The role of junctional adhesion molecules in vascular inflammation. Nat Rev Immunol. 2007 Jun;7(6):467–77. PMID: 17525755. 10.1038/nri2096

2) Mandell KJ, Parkos CA. The JAM family of proteins. Advanced drug delivery reviews. 2005 Apr 25;57(6):857–67. 10.1016/j.addr.2005.01.005

3) Ebnet K. Junctional Adhesion Molecules (JAMs): Cell Adhesion Receptors With Pleiotropic Functions in Cell Physiology and Development. Physiol Rev. 2017 Oct 1;97(4):1529–1554. PMID: 28931565. 10.1152/physrev.00004.2017

4) Wang J, Liu H. The Roles of Junctional Adhesion Molecules (JAMs) in Cell Migration. Front Cell Dev Biol. 2022 Mar 9;10:843671. PMID: 35356274; PMCID: PMC8959349. 10.3389/fcell.2022.843671

5) Solimando AG, Brandl A, Mattenheimer K, Graf C, Ritz M, Ruckdeschel A, Stühmer T, Mokhtari Z, Rudelius M, Dotterweich J, Bittrich M. JAM-A as a prognostic factor and new therapeutic target in multiple myeloma. Leukemia. 2018 Mar;32(3):736–43. 10.1038/leu.2017.287

6) Mendoza C, Mizrachi D. Using the Power of Junctional Adhesion Molecules Combined with the Target of CAR-T to Inhibit Cancer Proliferation, Metastasis and Eradicate Tumors. Biomedicines. 2022 Feb 4;10(2):381. PMID: 35203590; PMCID: PMC8962422. 10.3390/biomedicines10020381

7) Tokmakov AA, Kurotani A, Sato KI. Protein pI and Intracellular Localization. Front Mol Biosci. 2021 Nov 29;8:775736. PMID: 34912847; PMCID: PMC8667598. 10.3389/fmolb.2021.775736

8) Qing R, Hao S, Smorodina E, Jin D, Zalevsky A, Zhang S. Protein design: From the aspect of water solubility and stability. Chemical Reviews. 2022 Aug 3;122(18):14085–179. 10.1021/acs.chemrev.1c00757

9) Karagöl T, Karagöl A. pH-Dependent Membrane Binding Specificity of Synaptogyrins 1-3 Provides Mechanistic Insights into Synaptic Vesicle Regulation and Neurological Disease. bioRxiv. 2025 Mar 10:2025–03. 10.1101/2025.03.03.641025

10) Koch JS, Romero-Romero S, Höcker B. Stepwise introduction of stabilizing mutations reveals nonlinear additive effects in de novo TIM barrels. Protein Sci. 2024 Mar;33(3):e4926. PMID: 38380781; PMCID: PMC10880431. 10.1002/pro.4926

11) Karagöl T, Karagöl A, Zhang S. Structural bioinformatics studies of serotonin, dopamine and norepinephrine transporters and their AlphaFold2 predicted water-soluble QTY variants and uncovering the natural mutations of L-> Q, I-> T, F-> Y and Q-> L, T-> I and Y-> F. PloS one. 2024 Mar 22;19(3):e0300340. 10.1371/journal.pone.0300340

12) Karagöl A, Karagöl T. Adaptation to Solvent Environment in Toll-like Receptor 5: A Comparative Evolutionary Analysis of Membrane-bound and Soluble Forms in Epinephelus coioides. bioRxiv. 2025 Mar 11:2025–02. 10.1101/2025.02.28.640895

13) Karagöl T, Karagöl A, Zhang S. Co-evolution of alpha-helical transmembrane protein residues: large-scale variant profiling and complete mutational landscape of 2277 known PDB entries representing 504 unique human protein sequences. Journal of Molecular Evolution. 2025 Sep 24:1–9. 10.1007/s00239-025-10262-8

14) Zhang S et al (2018) QTY code enables design of detergent-free chemokine receptors that retain ligand-binding activities. Proc Natl Acad Sci USA 115(37):E8652–E8659. 10.1073/pnas.1811031115

15) Zhang S, Egli M. Hiding in plain sight: three chemically distinct α-helix types. Quarterly Reviews of Biophysics. 2022 Jan;55:e7. 10.1017/S0033583522000063

16) Beleva Guthrie V, Masica DL, Fraser A, Federico J, Fan Y, Camps M, Karchin R. Network Analysis of Protein Adaptation: Modeling the Functional Impact of Multiple Mutations. Mol Biol Evol. 2018 Jun 1;35(6):1507–1519. PMID: 29522102; PMCID: PMC5967520. 10.1093/molbev/msy036

17) Cheng J, Novati G, Pan J, Bycroft C, Žemgulytė A, Applebaum T, Pritzel A, Wong LH, Zielinski M, Sargeant T, Schneider RG, Senior AW, Jumper J, Hassabis D, Kohli P, Avsec Ž. Accurate proteome-wide missense variant effect prediction with AlphaMissense. Science. 2023 Sep 22;381(6664):eadg7492. Epub 2023 Sep 22. PMID: 37733863. 10.1126/science.adg7492

18) Hollingsworth SA, Dror RO. Molecular dynamics simulation for all. Neuron. 2018 Sep 19;99(6):1129–43. 10.1016/j.neuron.2018.08.011

19) Radak BK, Chipot C, Suh D, Jo S, Jiang W, Phillips JC, Schulten K, Roux B. Constant-pH molecular dynamics simulations for large biomolecular systems. Journal of chemical theory and computation. 2017 Dec 12;13(12):5933–44. 10.1021/acs.jctc.7b00875

20) Karagöl, A., Karagöl, T. & Zhang, S. Molecular Dynamic Simulations Reveal that Water-Soluble QTY-Variants of Glutamate Transporters EAA1, EAA2 and EAA3 Retain the Conformational Characteristics of Native Transporters. Pharm Res 41, 1965–1977 (2024). 10.1007/s11095-024-03769-0

21) Ben Chorin A, Masrati G, Kessel A, Narunsky A, Sprinzak J, Lahav S, Ashkenazy H, Ben-Tal N. ConSurf-DB: an accessible repository for the evolutionary conservation patterns of the majority of PDB proteins. Protein Science. 2020 Jan;29(1):258–67. 10.1002/pro.3779

22) Monné M, Nilsson I, Johansson M, Elmhed N, von Heijne G. Positively and negatively charged residues have different effects on the position in the membrane of a model transmembrane helix. J Mol Biol. 1998 Dec 11;284(4):1177–83. PMID: 9837735. 10.1006/jmbi.1998.2218

23) Kyte J, Doolittle RF. A simple method for displaying the hydropathic character of a protein. J Mol Biol. 1982 May 5;157(1):105–32. PMID: 7108955. 10.1016/0022-2836(82)90515-0

24) Sundberg RJ, Martin RB. Interactions of histidine and other imidazole derivatives with transition metal ions in chemical and biological systems. Chemical reviews. 1974 Aug 1;74(4):471–517. 10.1021/cr60290a003

25) Cressman R. Evolutionary dynamics and extensive form games. MIT Press; 2003.

26) Wang ZA, Cole PA. The chemical biology of reversible lysine post-translational modifications. Cell chemical biology. 2020 Aug 20;27(8):953–69. 10.1016/j.chembiol.2020.07.002

27) D’Auria S, Scire A, Varriale A, Scognamiglio V, Staiano M, Ausili A, Marabotti A, Rossi M, Tanfani F. Binding of glutamine to glutamine-binding protein from Escherichia coli induces changes in protein structure and increases protein stability. Proteins: Structure, Function, and Bioinformatics. 2005 Jan 1;58(1):80–7. 10.1002/prot.20289

28) White, K.A., Grillo-Hill, B.K. and Barber, D.L., 2017. Cancer cell behaviors mediated by dysregulated pH dynamics at a glance. Journal of cell science, 130(4), pp.663–669. 10.1242/jcs.195297

29) Lyu, Y., Thai, P., Trinh, P., Timofeyev, V., Ginsburg, K.S., Bossuyt, J., Bers, D.M., Yamoah, E.N., Chiamvimonvat, N. and Zhang, X.D., 2023. Dysregulation of intracellular pH in the failing heart. Biophysical Journal, 122(3), p.383a. 10.1016/j.bpj.2022.11.2101

30) Mutch WA, Hansen AJ. Extracellular pH changes during spreading depression and cerebral ischemia: mechanisms of brain pH regulation. Journal of Cerebral Blood Flow & Metabolism. 1984 Mar;4(1):17–27. 10.1038/jcbfm.1984.3

31) UniProt Consortium T. UniProt: the universal protein knowledgebase. Nucleic acids research. 2018 Mar 16;46(5):2699-. 10.1093/nar/gkh131

32) Bjellqvist B, Hughes GJ, Pasquali C, Paquet N, Ravier F, Sanchez JC, Frutiger S, Hochstrasser D. The focusing positions of polypeptides in immobilized pH gradients can be predicted from their amino acid sequences. Electrophoresis. 1993;14(1):1023–31. 10.1002/elps.11501401163

33) Bjellqvist B, Basse B, Olsen E, Celis JE. Reference points for comparisons of two-dimensional maps of proteins from different human cell types defined in a pH scale where isoelectric points correlate with polypeptide compositions. Electrophoresis. 1994;15(1):529–39. 10.1002/elps.1150150171

34) Gasteiger E, Hoogland C, Gattiker A, Duvaud SE, Wilkins MR, Appel RD, Bairoch A. Protein identification and analysis tools on the ExPASy server. In The proteomics protocols handbook 2005 Mar 22 (pp. 571–607). Totowa, NJ: Humana press. 10.1385/1-59259-890-0:571

35) Abramson, J., Adler, J., Dunger, J. et al. Accurate structure prediction of biomolecular interactions with AlphaFold 3. Nature 630, 493–500 (2024). 10.1038/s41586-024-07487-w

36) Harrower M, Brewer CA. ColorBrewer. org: an online tool for selecting colour schemes for maps. The Cartographic Journal. 2003 Jun 1;40(1):27–37. 10.1179/000870403235002042

37) Meng EC, Goddard TD, Pettersen EF, Couch GS, Pearson ZJ, Morris JH, Ferrin TE. UCSF ChimeraX: Tools for structure building and analysis. Protein Science. 2023 Nov;32(11):e4792. 10.1002/pro.4792

38) Sievers F, Higgins DG. Clustal Omega for making accurate alignments of many protein sequences. Protein Science. 2018 Jan;27(1):135–45. 10.1002/pro.3290

39) Waterhouse AM, Procter JB, Martin DM, Clamp M, Barton GJ. Jalview Version 2—a multiple sequence alignment editor and analysis workbench. Bioinformatics. 2009 May 1;25(9):1189–91. 10.1093/bioinformatics/btp033

40) Karagöl A, Karagöl T. An Evolutionary Statistics Toolkit for Simplified Sequence Analysis on Web with Client-Side Processing. bioRxiv. 2024 Aug 4:2024–08. 10.1101/2024.08.01.606148

41) Yariv B, Yariv E, Kessel A, Masrati G, Chorin AB, Martz E, Mayrose I, Pupko T, Ben-Tal N. Using evolutionary data to make sense of macromolecules with a “face-lifted” ConSurf. Protein Science. 2023 Mar;32(3):e4582. 10.1002/pro.4582

42) Ashkenazy H, Abadi S, Martz E, Chay O, Mayrose I, Pupko T, Ben-Tal N. ConSurf 2016: an improved methodology to estimate and visualize evolutionary conservation in macromolecules. Nucleic acids research. 2016 Jul 8;44(W1):W344–50. 10.1093/nar/gkw408

43) Celniker G, Nimrod G, Ashkenazy H, Glaser F, Martz E, Mayrose I, Pupko T, Ben-Tal N. ConSurf: using evolutionary data to raise testable hypotheses about protein function. Israel Journal of Chemistry. 2013 Apr;53(3-4):199–206. 10.1002/ijch.201200096

44) Landau M, Mayrose I, Rosenberg Y, Glaser F, Martz E, Pupko T, Ben-Tal N. ConSurf 2005: the projection of evolutionary conservation scores of residues on protein structures. Nucleic acids research. 2005 Jul 1;33(suppl_2):W299–302. 10.1093/nar/gki370

45) Suzek BE, Huang H, McGarvey P, Mazumder R, Wu CH. UniRef: comprehensive and non-redundant UniProt reference clusters. Bioinformatics. 2007 May 15;23(10):1282–8. 10.1093/bioinformatics/btm098

46) Schrödinger LLC. The PyMOL Molecular Graphics System, Version 3, https://www.pymol.org. Accessed 17 September 2024

47) Vinod HD. Generalized correlation and kernel causality with applications in development economics. Communications in Statistics-Simulation and Computation. 2017 Jul 3;46(6):4513–34. 10.1080/03610918.2015.1122048

48) Vinod HD. Generalized correlations and kernel causality using R package generalCorr. Available at SSRN 2782223. 2016 May 19. 10.2139/ssrn.2782223

49) Kim S. ppcor: an R package for a fast calculation to semi-partial correlation coefficients. Communications for statistical applications and methods. 2015 Nov 30;22(6):665. 10.5351/CSAM.2015.22.6.665

50) Zheng S, Shi NZ, Zhang Z. Generalized measures of correlation for asymmetry, nonlinearity, and beyond. Journal of the American Statistical Association. 2012 Sep 1;107(499):1239–52. 10.1080/01621459.2012.710509

51) R Core Team. R: A language and environment for statistical computing. Foundation for Statistical Computing, Vienna, Austria. 2013.

52) Hunter JD. Matplotlib: A 2D graphics environment. Computing in science & engineering. 2007 May 1;9(03):90–5. 10.1109/MCSE.2007.55

53) Csardi G, Nepusz T. The igraph software. Complex syst. 2006;1695:1–9.

54) Wei T, Simko V, Levy M, Xie Y, Jin Y, Zemla J. Package ‘corrplot’. Statistician. 2017 Oct 16;56(316):e24.

55) Tordai, H., Torres, O., Csepi, M. et al. Analysis of AlphaMissense data in different protein groups and structural context. Sci Data 11, 495 (2024). 10.1038/s41597-024-03327-8

56) Zimmerman DW. Comparative power of Student t test and Mann-Whitney U test for unequal sample sizes and variances. The Journal of Experimental Education. 1987 Apr 1;55(3):171–4. 10.1080/00220973.1987.10806451

57) Goulet-Pelletier JC, Cousineau D. A review of effect sizes and their confidence intervals, Part I: The Cohen’sd family. The Quantitative Methods for Psychology. 2018 Dec 1;14(4):242–65. 10.20982/tqmp.14.4.p242

58) Seabold S, Perktold J. Statsmodels: econometric and statistical modeling with python. SciPy. 2010 Jun 28;7(1):92–6. 10.25080/Majora-92bf1922-011

59) Abraham MJ, Murtola T, Schulz R, Páll S, Smith JC, Hess B, Lindahl E. GROMACS: High performance molecular simulations through multi-level parallelism from laptops to supercomputers. SoftwareX. 2015 Sep 1;1:19–25. 10.1016/j.softx.2015.06.001

60) Karagöl T, Karagöl A. Benchmarking GROMACS on Optimized Colab Processors and the Flexibility of Cloud Computing for Molecular Dynamics. bioRxiv. 2024 Nov 15:2024–11. 10.1101/2024.11.14.623563

61) Jo S, Kim T, Iyer VG, Im W. CHARMM-GUI: a web-based graphical user interface for CHARMM. J Comput Chem. 2008;29: 1859–1865. 10.1002/jcc.20945

62) Lee J, Cheng X, Jo S, MacKerell AD, Klauda JB, Im W. CHARMM-GUI input generator for NAMD, GROMACS, AMBER, OpenMM, and CHARMM/OpenMM simulations using the CHARMM36 additive force field. Biophysical journal. 2016 Feb 16;110(3):641a. 10.1021/acs.jctc.5b00935

63) Huang J, Rauscher S, Nawrocki G, Ran T, Feig M, de Groot BL, et al. CHARMM36m: an improved force field for folded and intrinsically disordered proteins. Nat Methods. 2017;14: 71–73. 10.1038/nmeth.4067

64) Sajeev-Sheeja A, Karagöl A, Karagöl T, Zhang S. Molecular dynamics simulations and structural bioinformatics of bacterial integral alpha-helical membrane enzymes and their AlphaFold2-predicted water-soluble QTY analogues. Molecular Simulation. 2025 Sep 30:1–5. 10.1080/08927022.2025.2562932

