## Supplementary Information for "pI as a Potential Factor Influencing Evolutionary Residue Selection and Structural Stability Among Junctional Adhesion Molecules"

### Table of Contents

1. Table S1. Amino acid counts in protein sequences of JAMs in *Homo sapiens*. Pg. 2
2. Table S2. Substitutions in multiple aligned sequences, observed more than 2 times. Pg. 3-4
3. Table S3. Correlations of amino acid conservation between proteins. Pg. 5
4. Table S4. Partial correlations of amino acid conservation between proteins. Pg. 6
5. Table S5. Amino acid score means of residue conservation among JAMs. Pg. 7
6. Table S6. Correlations of amino acid conservation with JAM protein pI. Pg. 8
7. Figure S1. JAM-A residue-wise evolutionary conservation profiles Pg. 9
8. Figure S2. JAM-B residue-wise evolutionary conservation profiles. Pg. 10
9. Figure S3. JAM-C residue-wise evolutionary conservation profiles. Pg. 11
10. Table S7. Centrality scores of JAM protein amino acids (ranked by centrality score). Pg. 12-13
11. Table S8. Comparison of AlphaMissense pathogenicity scores among proteins. Pg. 14-15
12. Table S9. Comparison of AlphaMissense pathogenicity scores of Acidic->Basic vs. others. Pg. 16
13. Table S10. Comparison of AlphaMissense pathogenicity scores of Basic->Acidic vs. others. Pg. 17
14. Table S11. Multiple comparison of Alphamissense means (Tukey HSD, FWER=0.05). Pg. 18
15. Figure S4. Structural analysis of molecular dynamics simulations of JAM-A. Pg. 19
16. Figure S5. Structural analysis of molecular dynamics simulations of JAM-B. Pg. 20
17. Figure S6. Structural analysis of molecular dynamics simulations of JAM-C. Pg. 21
18. Table S12. pH-dependent dynamics and evolutionary conservation of JAM-A residues. Pg. 22-26
19. Table S13. pH-dependent dynamics and evolutionary conservation of JAM-B residues. Pg. 27-31
20. Table S14. pH-dependent dynamics and evolutionary conservation of JAM-C residues. Pg. 32-36
21. Figure S7. Interplay between protein dynamics with residue conservation and pI of JAM-A. Pg. 37
22. Figure S8. Interplay between protein dynamics with residue conservation and pI of JAM-B. Pg. 38
23. Figure S9. Interplay between protein dynamics with residue conservation and pI of JAM-C. Pg. 39

**Table S1. Amino acid counts in protein sequences of JAMs in *Homo sapiens*.**

| AA | JAM-A[1] | JAM-B[1] | JAM-C[1] |
|---|---|---|---|
| **A** | 16 | 20 | 17 |
| **C** | 7 | 8 | 10 |
| **D** | 9 | 11 | 19 |
| **E** | 19 | 18 | 20 |
| **F** | 13 | 10 | 12 |
| **G** | 22 | 18 | 20 |
| **H** | 2 | 3 | 6 |
| **I** | 14 | 11 | 20 |
| **K** | 16 | 21 | 15 |
| **L** | 23 | 25 | 26 |
| **M** | 5 | 5 | 3 |
| **N** | 13 | 13 | 15 |
| **P** | 17 | 10 | 16 |
| **Q** | 5 | 14 | 10 |
| **R** | 14 | 19 | 25 |
| **S** | 34 | 30 | 20 |
| **T** | 27 | 20 | 16 |
| **V** | 29 | 27 | 26 |
| **W** | 3 | 2 | 3 |
| **Y** | 11 | 13 | 11 |

[1]The protein sequence data were retrieved from UniProt website (https://www.uniprot.org) (IDs: Q9Y624, P57087, Q9BX67)

**Table S2. Substitutions in multiple aligned sequences, observed more than 2 times.**
Full list can be found at:
https://github.com/karagol-taner/pI_evolution_JAMs/blob/main/MSA_analysis/SUM.xlsx

| Protein 1 | Protein 2 | AA change | Change in pI (25 C) | Change in Hydropathy | Time Observed | Total pI Change |
|---|---|---|---|---|---|---|
| JAM-A | JAM-B | I>L | -0.04 | -0.7 | 4 | -0.16 |
| | | L>V | -0.02 | 0.4 | 4 | -0.08 |
| | | S>K | 4.06 | -3.1 | 4 | 16.24 |
| | | N>Q | 0.24 | 0 | 4 | 0.96 |
| | | K>T | -4.14 | 3.2 | 4 | -16.56 |
| | | S>A | 0.32 | 2.6 | 3 | 0.96 |
| | | T>E | -2.38 | -2.8 | 3 | -7.14 |
| | | L>I | 0.04 | 0.7 | 3 | 0.12 |
| | | T>R | 5.16 | -3.8 | 3 | 15.48 |
| | | S>N | -0.27 | -2.7 | 3 | -0.81 |
| | | P>A | -0.3 | 3.4 | 3 | -0.9 |
| | | I>V | -0.06 | -0.3 | 3 | -0.18 |
| | | E>D | -0.45 | 0 | 3 | -1.35 |
| | | G>S | -0.29 | -0.4 | 3 | -0.87 |
| | | V>I | 0.06 | 0.3 | 3 | 0.18 |
| JAM-A | JAM-C | V>I | 0.06 | 0.3 | 6 | 0.36 |
| | | G>D | -3.2 | -3.1 | 5 | -16 |
| | | S>R | 5.08 | -3.7 | 4 | 20.32 |
| | | I>V | -0.06 | -0.3 | 4 | -0.24 |
| | | T>S | 0.08 | -0.1 | 4 | 0.32 |
| | | G>A | 0.03 | 2.2 | 3 | 0.09 |
| | | V>P | 0.34 | -5.8 | 3 | 1.02 |
| | | I>L | -0.04 | -0.7 | 3 | -0.12 |
| | | S>A | 0.32 | 2.6 | 3 | 0.96 |
| | | T>N | -0.19 | -2.8 | 3 | -0.57 |
| | | P>S | -0.62 | 0.8 | 3 | -1.86 |
| | | S>D | -2.91 | -2.7 | 3 | -8.73 |
| | | A>G | -0.03 | -2.2 | 3 | -0.09 |
| | | E>D | -0.45 | 0 | 3 | -1.35 |
| | | S>E | -2.46 | -2.7 | 3 | -7.38 |
| | | K>R | 1.02 | -0.6 | 3 | 3.06 |
| | | L>V | -0.02 | 0.4 | 3 | -0.06 |
| JAM-B | JAM-A | L>I | 0.04 | 0.7 | 4 | 0.16 |
| | | V>L | 0.02 | -0.4 | 4 | 0.08 |
| | | K>S | -4.06 | 3.1 | 4 | -16.24 |
| | | Q>N | -0.24 | 0 | 4 | -0.96 |
| | | T>K | 4.14 | -3.2 | 4 | 16.56 |
| | | A>S | -0.32 | -2.6 | 3 | -0.96 |
| | | E>T | 2.38 | 2.8 | 3 | 7.14 |
| | | I>L | -0.04 | -0.7 | 3 | -0.12 |
| | | R>T | -5.16 | 3.8 | 3 | -15.48 |
| | | N>S | 0.27 | 2.7 | 3 | 0.81 |
| | | A>P | 0.3 | -3.4 | 3 | 0.9 |

| | | | | | | |
|---|---|---|---|---|---|---|
| | | V>I | 0.06 | 0.3 | 3 | 0.18 |
| | | D>E | 0.45 | 0 | 3 | 1.35 |
| | | S>G | 0.29 | 0.4 | 3 | 0.87 |
| | | I>V | -0.06 | -0.3 | 3 | -0.18 |
| | | | | | | -5.89 |
| JAM-B | JAM-C | A>V | -0.04 | 2.4 | 4 | -0.16 |
| | | Y>F | -0.18 | 4.1 | 4 | -0.72 |
| | | L>I | 0.04 | 0.7 | 4 | 0.16 |
| | | T>K | 4.14 | -3.2 | 4 | 16.56 |
| | | S>V | 0.28 | 5 | 4 | 1.12 |
| | | S>P | 0.62 | -0.8 | 3 | 1.86 |
| | | V>I | 0.06 | 0.3 | 3 | 0.18 |
| | | A>S | -0.32 | -2.6 | 3 | -0.96 |
| | | K>S | -4.06 | 3.1 | 3 | -12.18 |
| | | I>L | -0.04 | -0.7 | 3 | -0.12 |
| | | E>T | 2.38 | 2.8 | 3 | 7.14 |
| | | T>S | 0.08 | -0.1 | 3 | 0.24 |
| | | K>R | 1.02 | -0.6 | 3 | 3.06 |
| JAM-C | JAM-A | I>V | -0.06 | -0.3 | 6 | -0.36 |
| | | D>G | 3.2 | 3.1 | 5 | 16 |
| | | R>S | -5.08 | 3.7 | 4 | -20.32 |
| | | V>I | 0.06 | 0.3 | 4 | 0.24 |
| | | S>T | -0.08 | 0.1 | 4 | -0.32 |
| | | A>G | -0.03 | -2.2 | 3 | -0.09 |
| | | P>V | -0.34 | 5.8 | 3 | -1.02 |
| | | L>I | 0.04 | 0.7 | 3 | 0.12 |
| | | A>S | -0.32 | -2.6 | 3 | -0.96 |
| | | N>T | 0.19 | 2.8 | 3 | 0.57 |
| | | S>P | 0.62 | -0.8 | 3 | 1.86 |
| | | D>S | 2.91 | 2.7 | 3 | 8.73 |
| | | G>A | 0.03 | 2.2 | 3 | 0.09 |
| | | D>E | 0.45 | 0 | 3 | 1.35 |
| | | E>S | 2.46 | 2.7 | 3 | 7.38 |
| | | R>K | -1.02 | 0.6 | 3 | -3.06 |
| | | V>L | 0.02 | -0.4 | 3 | 0.06 |
| JAM-C | JAM-B | V>A | 0.04 | -2.4 | 4 | 0.16 |
| | | F>Y | 0.18 | -4.1 | 4 | 0.72 |
| | | I>L | -0.04 | -0.7 | 4 | -0.16 |
| | | K>T | -4.14 | 3.2 | 4 | -16.56 |
| | | V>S | -0.28 | -5 | 4 | -1.12 |
| | | P>S | -0.62 | 0.8 | 3 | -1.86 |
| | | I>V | -0.06 | -0.3 | 3 | -0.18 |
| | | S>A | 0.32 | 2.6 | 3 | 0.96 |
| | | S>K | 4.06 | -3.1 | 3 | 12.18 |
| | | L>I | 0.04 | 0.7 | 3 | 0.12 |
| | | T>E | -2.38 | -2.8 | 3 | -7.14 |
| | | S>T | -0.08 | 0.1 | 3 | -0.24 |
| | | R>K | -1.02 | 0.6 | 3 | -3.06 |

**Table S3. Correlations of amino acid conservation between proteins.**

| | JAM-A/JAM-B | JAM-B/JAM-C | JAM-A/JAM-C |
|---|---|---|---|
| A | 0.8394629 | 0.7211387 | 0.6611426 |
| C | 0.874546 | 0.9852315 | 0.8614692 |
| D | 0.8166244 | 0.9333274 | 0.7450934 |
| E | 0.8590077 | 0.9375555 | 0.818111 |
| F | 0.8340313 | 0.9529493 | 0.834056 |
| G | 0.8833609 | 0.9405178 | 0.8689652 |
| H | 0.4530124 | 0.7280754 | 0.4174055 |
| I | 0.6523397 | 0.9444954 | 0.6502374 |
| K | 0.7518182 | 0.8984889 | 0.7140847 |
| L | 0.8429708 | 0.8406604 | 0.7959727 |
| M | 0.6568363 | 0.68905 | 0.9163494 |
| N | 0.8765369 | 0.9449151 | 0.8455381 |
| P | 0.8444415 | 0.9182613 | 0.8333685 |
| Q | 0.3569415 | 0.8193384 | 0.3748823 |
| R | 0.7559558 | 0.8880582 | 0.7559558 |
| S | 0.7480981 | 0.8740332 | 0.7364964 |
| T | 0.7263794 | 0.8933799 | 0.6669663 |
| V | 0.865871 | 0.9352707 | 0.8226961 |
| W | 0.933395 | 0.9914435 | 0.9213462 |
| Y | 0.8498277 | 0.9547976 | 0.7996234 |
| **Mean** | 0.771072875 | 0.88954941 | 0.75198801 |
| **Q1** | 0.742668425 | 0.86569 | 0.7023051 |
| **Q3** | 0.860723525 | 0.944600325 | 0.836926525 |
| **IQR** | 0.1180551 | 0.078910325 | 0.134621425 |
| **Q3 + 1.5IQR** | 1.037806175 | 1.062965813 | 1.038858663 |
| **Q1 - 1.5IQR** | 0.565585775 | 0.747324513 | 0.500372963 |
| **SDEV (σ)** | 0.146523856 | 0.087637341 | 0.144873947 |
| **Mean + 2σ** | 1.064120587 | 1.064824091 | 1.041735905 |
| **Mean - 2σ** | 0.478025163 | 0.714274729 | 0.462240115 |

**Table S4. Partial correlations of amino acid conservation between proteins.**

| | pcor.test(Xa, Xb, Xc) | pcor.test(Xb, Xc, Xa) | pcor.test(Xa, Xc, Xb) |
|---|---|---|---|
| A | 0.4074866 | 0.6977799 | 0.1481468 |
| C | 0.2967109 | 0.9414373 | -0.001938557 |
| D | 0.5061814 | 0.8439195 | -0.08244703 |
| E | 0.4598549 | 0.7975012 | 0.07156181 |
| F | 0.2345158 | 0.8454167 | 0.2347886 |
| G | 0.39305 | 0.7454731 | 0.2395728 |
| H | 0.2393702 | 0.6653091 | 0.1433071 |
| I | 0.1530246 | 0.9036145 | 0.1369713 |
| K | 0.3586483 | 0.7834235 | 0.1332973 |
| L | 0.5302457 | 0.5210554 | 0.2997216 |
| M | 0.08762657 | 0.2887 | 0.8486584 |
| N | 0.443899 | 0.7929035 | 0.1097133 |
| P | 0.3618325 | 0.7246127 | 0.2732185 |
| Q | 0.09367089 | 0.7915991 | 0.1539114 |
| R | 0.3964115 | 0.7643952 | 0.1148177 |
| S | 0.3175799 | 0.7197083 | 0.2562964 |
| T | 0.3898953 | 0.7985038 | 0.05839834 |
| V | 0.4792511 | 0.783853 | 0.07269942 |
| W | 0.3927921 | 0.9423867 | -0.08672027 |
| Y | 0.4837407 | 0.8696834 | -0.07525352 |
| **Mean** | 0.351289398 | 0.761063795 | 0.15243607 |
| **Q1** | 0.282375725 | 0.7233866 | 0.068270943 |
| **Q3** | 0.447887975 | 0.8442938 | 0.23598465 |
| **IQR** | 0.16551225 | 0.1209072 | 0.167713708 |
| **Q3 + 1.5IQR** | 0.69615635 | 1.0256546 | 0.487555211 |
| **Q1 - 1.5IQR** | 0.03410735 | 0.5420258 | -0.183299619 |
| **SDEV (σ)** | 0.131005254 | 0.147684842 | 0.19998684 |
| **Mean + 2σ** | 0.613299906 | 1.056433479 | 0.55240975 |
| **Mean - 2σ** | 0.08927889 | 0.465694111 | -0.247537611 |

**Table S5. Amino acid score means of residue conservation among JAMs.**

| | JAM-A | JAM-B | JAM-C |
|---|---|---|---|
| mean(A) | 6.70111 | 6.32636 | 6.10198 |
| mean(C) | 1.98228 | 2.96597 | 2.80035 |
| mean(D) | 4.61153 | 4.58065 | 4.8851 |
| mean(E) | 5.08388 | 5.74389 | 6.29269 |
| mean(F) | 4.49373 | 4.03835 | 4.39375 |
| mean(G) | 7.80973 | 6.47073 | 6.73638 |
| mean(H) | 0.80296 | 1.7013 | 1.29172 |
| mean(I) | 3.52881 | 4.61364 | 5.08718 |
| mean(K) | 6.54316 | 6.83717 | 6.30563 |
| mean(L) | 7.57567 | 8.1758 | 7.98578 |
| mean(M) | 1.46355 | 1.5684 | 1.17496 |
| mean(N) | 4.12375 | 4.34793 | 4.56065 |
| mean(P) | 5.86126 | 4.76077 | 4.78703 |
| mean(Q) | 3.05098 | 3.08572 | 3.15749 |
| mean(R) | 4.61861 | 6.03542 | 6.05811 |
| mean(S) | 9.87842 | 8.66351 | 8.00484 |
| mean(T) | 7.37002 | 6.09425 | 5.92379 |
| mean(V) | 9.2352 | 9.16294 | 9.23752 |
| mean(W) | 1.11813 | 1.00816 | 1.04346 |
| mean(Y) | 4.11395 | 3.81674 | 4.16921 |

**Table S6. Correlations of amino acid conservation with JAM protein pI.**

| Rank | Aa. | Correlation with JAM protein pI |
|---|---|---|
| 1 | K | 0.991 |
| 2 | Y | -0.983 |
| 3 | V | -0.955 |
| 4 | M | 0.897 |
| 5 | F | -0.858 |
| 6 | D | -0.81 |
| 7 | H | 0.619 |
| 8 | Q | -0.503 |
| 9 | W | -0.492 |
| 10 | L | 0.488 |
| 11 | G | -0.374 |
| 12 | C | 0.345 |
| 13 | N | -0.309 |
| 14 | E | -0.273 |
| 15 | P | -0.214 |
| 16 | A | 0.184 |
| 17 | R | 0.18 |
| 18 | S | 0.159 |
| 19 | I | -0.106 |
| 20 | T | -0.0863 |

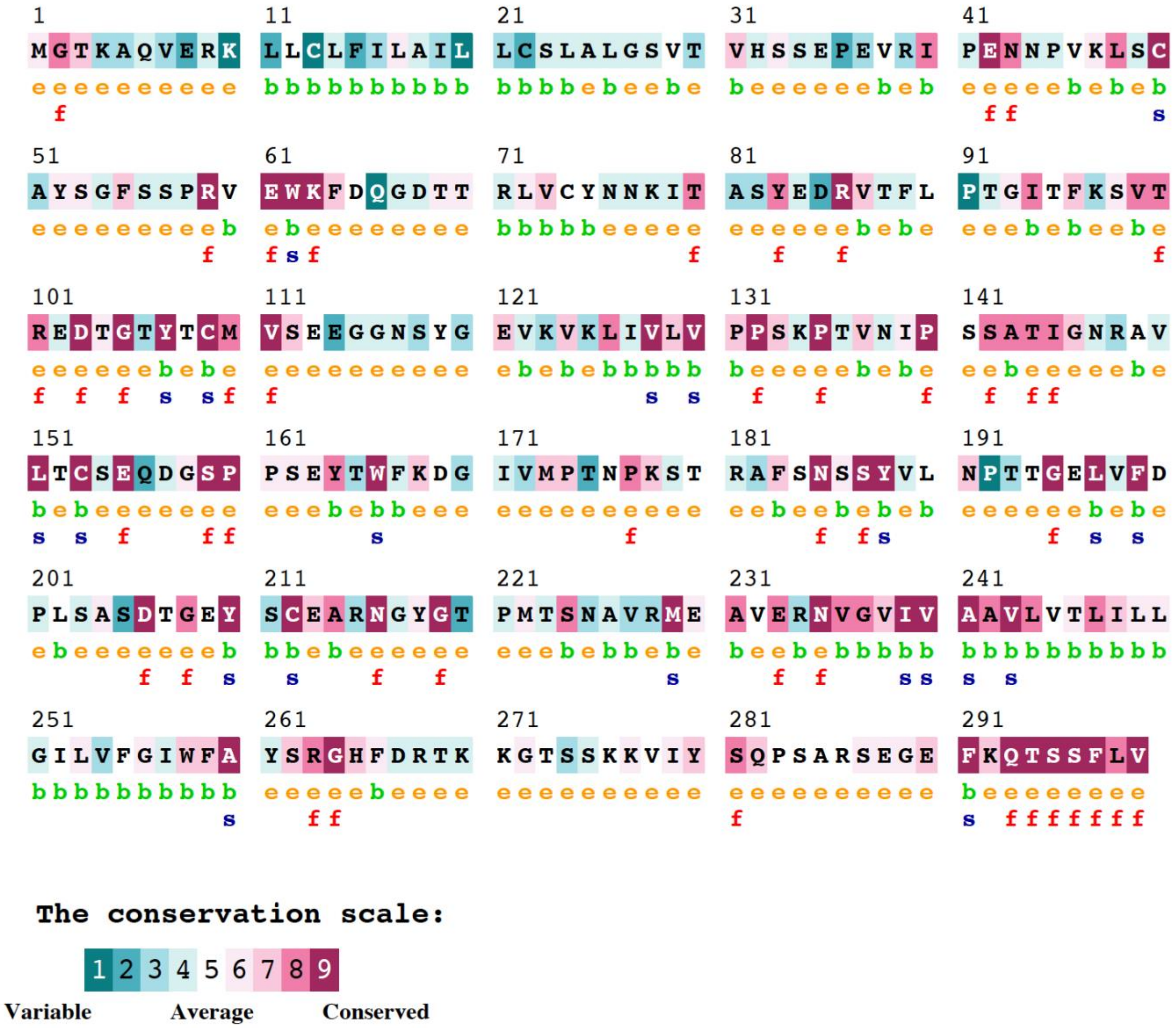

**Figure S1. JAM-A residue-wise evolutionary conservation profiles.** Evolutionary conservation grades of each amino acid residue predicted by ConSurf server; visualized by the color-coding scheme of nine colors, ranging from turquoise (variable) through white (average) through burgundy (conserved) represents conservation grades 1 to 9, in order of increasing conservation (1= Variable, 5= Average, 9= Conserved).

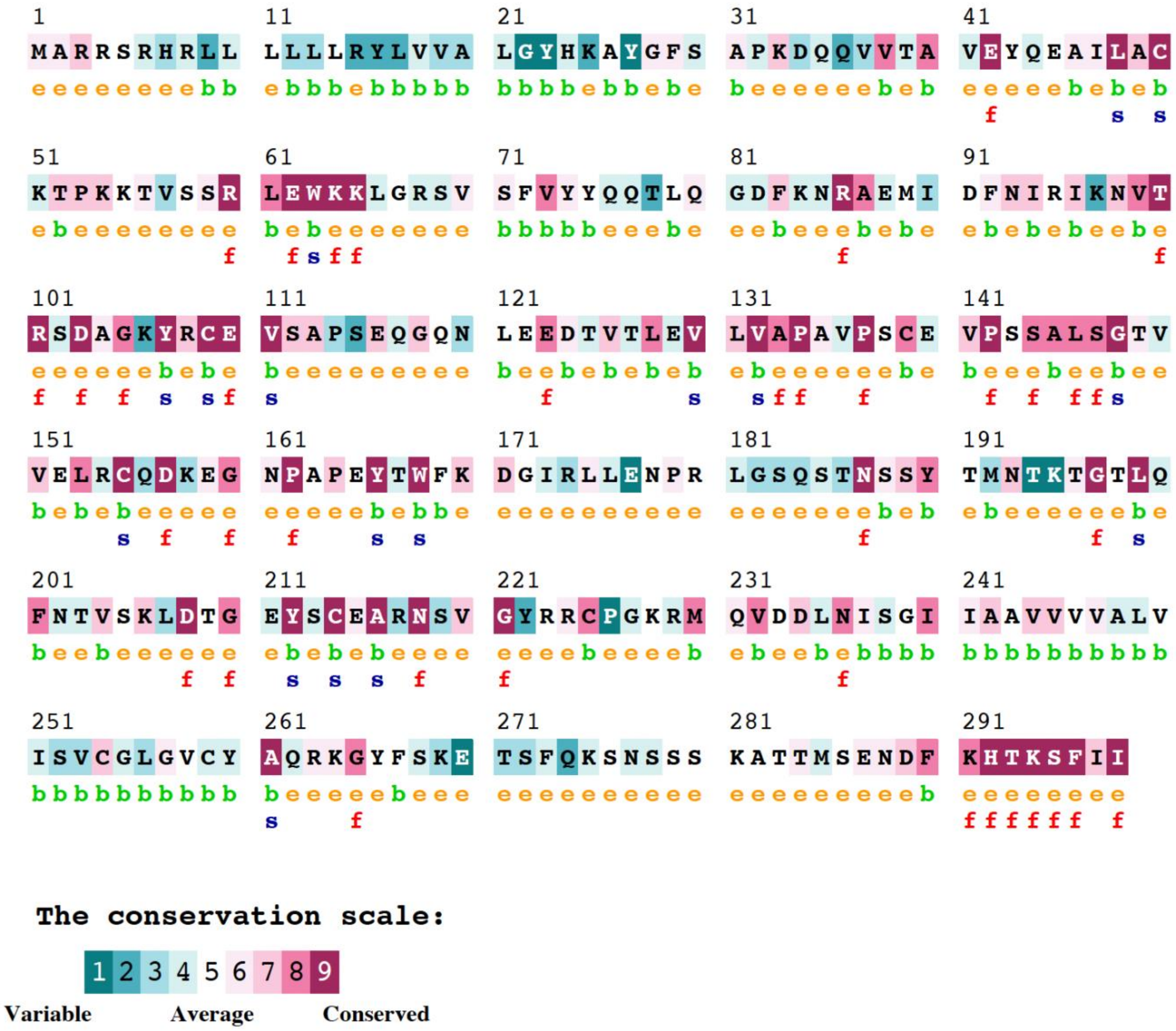

**Figure S2. JAM-B residue-wise evolutionary conservation profiles.** Evolutionary conservation grades of each amino acid residue predicted by ConSurf server; visualized by the color-coding scheme of nine colors, ranging from turquoise (variable) through white (average) through burgundy (conserved) represents conservation grades 1 to 9, in order of increasing conservation (1= Variable, 5= Average, 9= Conserved).

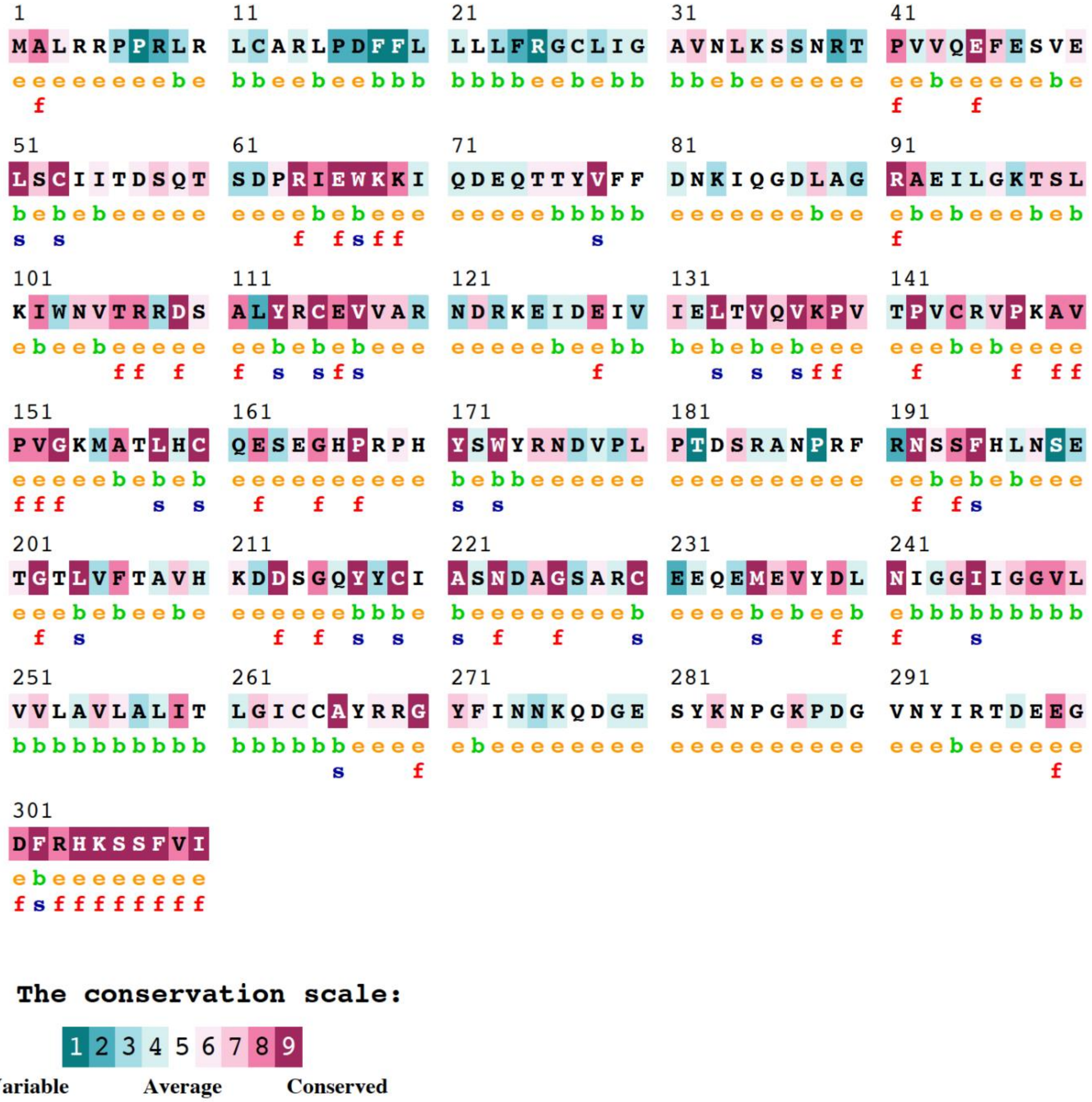

**Figure S3. JAM-C residue-wise evolutionary conservation profiles.** Evolutionary conservation grades of each amino acid residue predicted by ConSurf server; visualized by the color-coding scheme of nine colors, ranging from turquoise (variable) through white (average) through burgundy (conserved) represents conservation grades 1 to 9, in order of increasing conservation (1= Variable, 5= Average, 9= Conserved).

**Table S7. Centrality scores of JAM protein amino acids (ranked by centrality score).**

| Residue[1] | Betweenness Centrality | Degree Centrality | Weighted Degree Centrality | Centrality Score |
|---|---|---|---|---|
| Cb | 0.15838691 | 0.06779661 | 2.06625949 | 100 |
| Kb | 0.12799532 | 0.28813559 | 3.71014343 | 80.8118081 |
| Cc | 0.09818819 | 0.05084746 | 1.94813685 | 61.9926199 |
| Ic | 0.0742256 | 0.42372881 | 4.73339678 | 46.8634686 |
| Wa | 0.0666277 | 0.05084746 | 1.95617735 | 42.0664207 |
| Lc | 0.06487434 | 0.47457627 | 5.13997964 | 40.9594096 |
| Tb | 0.06078317 | 0.25423729 | 3.46861048 | 38.3763838 |
| Ab | 0.05084746 | 0.3220339 | 3.42829775 | 32.103321 |
| Va | 0.04792519 | 0.54237288 | 6.11311812 | 30.2583026 |
| Fc | 0.04617183 | 0.3559322 | 4.01079598 | 29.1512915 |
| Yb | 0.04266511 | 0.28813559 | 3.57660603 | 26.9372694 |
| Lb | 0.0414962 | 0.45762712 | 5.10996933 | 26.199262 |
| La | 0.03740503 | 0.45762712 | 4.93585812 | 23.6162362 |
| Ta | 0.03565167 | 0.25423729 | 3.15342508 | 22.5092251 |
| Qb | 0.03448276 | 0.37288136 | 4.18386942 | 21.7712177 |
| Ib | 0.03331385 | 0.45762712 | 5.20345212 | 21.0332103 |
| Rc | 0.03214494 | 0.23728814 | 3.05023361 | 20.295203 |
| Pa | 0.03214494 | 0.10169492 | 2.14060485 | 20.295203 |
| Mb | 0.03039158 | 0.06779661 | 1.85555627 | 19.1881919 |
| Fa | 0.02922268 | 0.25423729 | 3.09881797 | 18.4501845 |
| Ac | 0.02805377 | 0.27118644 | 2.98737796 | 17.7121771 |
| Ia | 0.01870251 | 0.30508475 | 3.4915564 | 11.8081181 |
| Vc | 0.01811806 | 0.52542373 | 5.88823004 | 11.4391144 |
| Ka | 0.01811806 | 0.3559322 | 4.09667616 | 11.4391144 |
| Fb | 0.01753361 | 0.30508475 | 3.80971047 | 11.0701107 |
| Ya | 0.01344243 | 0.18644068 | 2.822429 | 8.48708487 |
| Vb | 0.01285798 | 0.55932203 | 6.37948398 | 8.11808118 |
| Gc | 0.01227352 | 0.22033898 | 3.06978629 | 7.74907749 |
| Yc | 0.01052016 | 0.23728814 | 3.18377349 | 6.64206642 |
| Ec | 0.00993571 | 0.30508475 | 3.85634032 | 6.27306273 |
| Nc | 0.00993571 | 0.28813559 | 3.71649098 | 6.27306273 |
| Kc | 0.0087668 | 0.30508475 | 3.73516806 | 5.53505535 |
| Gb | 0.0087668 | 0.20338983 | 3.01366469 | 5.53505535 |
| Qc | 0.00818235 | 0.3220339 | 3.81062827 | 5.16605166 |

| | | | | |
|---|---|---|---|---|
| **Tc** | 0.00818235 | 0.23728814 | 3.1425057 | 5.16605166 |
| **Ha** | 0.00818235 | 0.18644068 | 2.14517381 | 5.16605166 |
| **Na** | 0.0075979 | 0.22033898 | 3.07184647 | 4.79704797 |
| **Aa** | 0.00701344 | 0.13559322 | 2.01068492 | 4.42804428 |
| **Ea** | 0.00642899 | 0.30508475 | 3.88973604 | 4.05904059 |
| **Rb** | 0.00642899 | 0.23728814 | 3.1973414 | 4.05904059 |
| **Ra** | 0.00409117 | 0.18644068 | 2.63929264 | 2.58302583 |
| **Sa** | 0.00350672 | 0.25423729 | 3.27923613 | 2.21402214 |
| **Ga** | 0.00350672 | 0.20338983 | 3.00837943 | 2.21402214 |
| **Pb** | 0.00350672 | 0.10169492 | 2.18856127 | 2.21402214 |
| **Ma** | 0.00292227 | 0.08474576 | 1.93084278 | 1.84501845 |
| **Sc** | 0.00175336 | 0.28813559 | 3.91471943 | 1.10701107 |
| **Nb** | 0.00175336 | 0.27118644 | 3.63652374 | 1.10701107 |
| **Eb** | 0.00175336 | 0.25423729 | 3.54939634 | 1.10701107 |
| **Db** | 0.00116891 | 0.3220339 | 4.13239963 | 0.73800738 |
| **Da** | 0.00058445 | 0.27118644 | 3.63681227 | 0.36900369 |
| **Sb** | 0 | 0.30508475 | 4.14926801 | 0 |
| **Dc** | 0 | 0.27118644 | 3.6381247 | 0 |
| **Hb** | 0 | 0.11864407 | 2.43137756 | 0 |
| **Hc** | 0 | 0.08474576 | 2.3301053 | 0 |
| **Qa** | 0 | 0.10169492 | 1.95952858 | 0 |
| **Wb** | 0 | 0.03389831 | 1.92483856 | 0 |
| **Wc** | 0 | 0.03389831 | 1.91278976 | 0 |
| **Pc** | 0 | 0.03389831 | 1.75162971 | 0 |
| **Ca** | 0 | 0.03389831 | 1.73601521 | 0 |
| **Mc** | 0 | 0.03389831 | 1.60539941 | 0 |

[1]Uppercase letters denote amino acids, while lowercase letters indicate proteins: *Xa* = JAM-A, *Xb* = JAM-B, *Xc* = JAM-C, where *X* represents the amino acid single letter code.

**Table S8. Comparison of AlphaMissense pathogenicity scores among proteins.**

| Mutation Type | Metric | JAM-A | JAM-B | JAM-C |
|---|---|---|---|---|
| **All mutations** | Count | 5662 | 5643 | 5871 |
| | Mean Pathogenicity | 0.51 | 0.4487 | 0.531 |
| | Median | 0.4436 | 0.3473 | 0.4863 |
| | Std Deviation | 0.3012 | 0.3164 | 0.3405 |
| | Variance | 0.0907 | 0.1001 | 0.116 |
| **Acidic → Basic** | Count | 56 | 58 | 78 |
| | Max Score | 0.9982 | 0.9924 | 0.9982 |
| | Mean Pathogenicity | 0.441 | 0.4191 | 0.4574 |
| | Median | 0.3292 | 0.3351 | 0.3881 |
| | Min Score | 0.1428 | 0.0707 | 0.0654 |
| | Std Deviation | 0.2703 | 0.2944 | 0.3318 |
| | Variance | 0.0731 | 0.0867 | 0.1101 |
| **Basic → Acidic** | Count | 60 | 80 | 80 |
| | Max Score | 0.9974 | 0.9943 | 0.9984 |
| | Mean Pathogenicity | 0.4636 | 0.4235 | 0.4814 |
| | Median | 0.4166 | 0.3588 | 0.3627 |
| | Min Score | 0.1104 | 0.0705 | 0.0963 |
| | Std Deviation | 0.2339 | 0.2746 | 0.3038 |
| | Variance | 0.0547 | 0.0754 | 0.0923 |
| **Other → Basic** | Count | 536 | 514 | 538 |
| | Max Score | 0.9999 | 0.9997 | 1 |
| | Mean Pathogenicity | 0.5533 | 0.4908 | 0.5812 |
| | Median | 0.5115 | 0.4032 | 0.6332 |
| | Min Score | 0.0721 | 0.0575 | 0.0341 |
| | Std Deviation | 0.3053 | 0.3357 | 0.3585 |
| | Variance | 0.0932 | 0.1127 | 0.1285 |
| **Other → Acidic** | Count | 540 | 536 | 540 |
| | Max Score | 1 | 0.9998 | 1 |
| | Mean Pathogenicity | 0.5821 | 0.5133 | 0.605 |
| | Median | 0.5495 | 0.4273 | 0.6424 |
| | Min Score | 0.0752 | 0.0566 | 0.0613 |
| | Std Deviation | 0.2955 | 0.3304 | 0.3442 |
| | Variance | 0.0873 | 0.1092 | 0.1185 |
| **Acidic → Other** | Count | 504 | 522 | 702 |
| | Max Score | 0.9999 | 0.9994 | 0.9998 |
| | Mean Pathogenicity | 0.5456 | 0.5156 | 0.5245 |

| | | | | |
|---|---|---|---|---|
| | Median | 0.5018 | 0.4604 | 0.4813 |
| | Min Score | 0.1019 | 0.0531 | 0.0559 |
| | Std Deviation | 0.2966 | 0.3095 | 0.3315 |
| | Variance | 0.088 | 0.0958 | 0.1099 |
| **Basic → Other** | Count | 540 | 720 | 720 |
| | Max Score | 0.9993 | 0.9978 | 0.9994 |
| | Mean Pathogenicity | 0.4754 | 0.417 | 0.4688 |
| | Median | 0.4189 | 0.3421 | 0.3699 |
| | Min Score | 0.107 | 0.0669 | 0.0605 |
| | Std Deviation | 0.2403 | 0.2596 | 0.2999 |
| | Variance | 0.0577 | 0.0674 | 0.0899 |

**Table S9. Comparison of AlphaMissense pathogenicity scores of Acidic->Basic vs. others.**

| Dataset | Metric | Acidic->Basic | Others |
|---|---|---|---|
| All Proteins (JAM-A/JAM-B/ JAM-C) | **N** | 192 | 24308 |
| | **Mean** | 0.4258 | 0.5028 |
| | **Median** | 0.4169 | 0.5025 |
| | **T-test** | t = -3.923, p = 1.215e-04 | |
| | **Mann–Whitney U** | U = 1975604.0, p = 2.447e-04 | |
| | **Cohen's d** | -0.269 | |
| JAM-B | **N** | 58 | 8015 |
| | **Mean** | 0.3741 | 0.4606 |
| | **Median** | 0.3502 | 0.4543 |
| | **T-test** | t = -2.593, p = 1.201e-02 | |
| | **Mann–Whitney U** | U = 191786.0, p = 2.151e-02 | |
| | **Cohen's d** | -0.305 | |
| JAM-A | **N** | 56 | 7842 |
| | **Mean** | 0.3952 | 0.5187 |
| | **Median** | 0.4054 | 0.5191 |
| | **T-test** | t = -3.586, p = 7.062e-04 | |
| | **Mann–Whitney U** | U = 164505.5, p = 1.198e-03 | |
| | **Cohen's d** | -0.455 | |
| JAM-C | **N** | 78 | 8451 |
| | **Mean** | 0.4863 | 0.5279 |
| | **Median** | 0.4762 | 0.5331 |
| | **T-test** | t = -1.286, p = 2.022e-01 | |
| | **Mann–Whitney U** | U = 303086.0, p = 2.207e-01 | |
| | **Cohen's d** | -0.14 | |

**Table S10. Comparison of AlphaMissense pathogenicity scores of Basic->Acidic vs. others.**

| Dataset | Metric | Basic->Acidic | Others |
|---|---|---|---|
| All Proteins (JAM-A/JAM-B/ JAM-C) | **N** | 220 | 24280 |
| | **Mean** | 0.4732 | 0.5024 |
| | **Median** | 0.4592 | 0.502 |
| | **T-test** | t = -1.655, p = 9.926e-02 | |
| | **Mann–Whitney U** | U = 2510465.0, p = 1.246e-01 | |
| | **Cohen's d** | -0.102 | |
| JAM-B | **N** | 80 | 7993 |
| | **Mean** | 0.4259 | 0.4604 |
| | **Median** | 0.4151 | 0.4543 |
| | **T-test** | t = -1.240, p = 2.185e-01 | |
| | **Mann–Whitney U** | U = 297314.0, p = 2.799e-01 | |
| | **Cohen's d** | -0.122 | |
| JAM-A | **N** | 60 | 7838 |
| | **Mean** | 0.478 | 0.5182 |
| | **Median** | 0.4863 | 0.5187 |
| | **T-test** | t = -1.378, p = 1.733e-01 | |
| | **Mann–Whitney U** | U = 214602.5, p = 2.431e-01 | |
| | **Cohen's d** | -0.148 | |
| JAM-C | **N** | 80 | 8449 |
| | **Mean** | 0.517 | 0.5276 |
| | **Median** | 0.5087 | 0.533 |
| | **T-test** | t = -0.323, p = 7.472e-01 | |
| | **Mann–Whitney U** | U = 331218.0, p = 7.583e-01 | |
| | **Cohen's d** | -0.036 | |

**Table S11. Multiple comparison of Alphamissense means (Tukey HSD, FWER=0.05).**

| Group1 | Group2 | meandiff | p-adj | lower | upper | reject |
|---|---|---|---|---|---|---|
| JAM-A | JAM-B | -0.0578 | 0 | -0.0684 | -0.0473 | True |
| JAM-A | JAM-C | 0.0097 | 0.0755 | -0.0007 | 0.0201 | False |
| JAM-B | JAM-C | 0.0675 | 0 | 0.0572 | 0.0778 | TRUE |
| Acidic>Basic | Acidic>Other | 0.1097 | 0 | 0.0459 | 0.1735 | TRUE |
| Acidic>Basic | All | 0.0694 | 0.0135 | 0.0086 | 0.1303 | TRUE |
| Acidic>Basic | Basic>Acidic | 0.0474 | 0.6249 | -0.0354 | 0.1302 | FALSE |
| Acidic>Basic | Basic>Other | 0.0304 | 0.7943 | -0.033 | 0.0938 | FALSE |
| Acidic>Basic | Other>Acidic | 0.1433 | 0 | 0.0793 | 0.2073 | TRUE |
| Acidic>Basic | Other>Basic | 0.1164 | 0 | 0.0523 | 0.1804 | TRUE |
| Acidic>Other | All | -0.0403 | 0 | -0.0615 | -0.0191 | TRUE |
| Acidic>Other | Basic->Acidic | -0.0624 | 0.0357 | -0.1224 | -0.0023 | TRUE |
| Acidic>Other | Basic>Other | -0.0793 | 0 | -0.1069 | -0.0517 | TRUE |
| Acidic>Other | Other>Acidic | 0.0336 | 0.0116 | 0.0045 | 0.0626 | TRUE |
| Acidic>Other | Other>Basic | 0.0066 | 0.9942 | -0.0225 | 0.0358 | FALSE |
| All | Basic->Acidic | -0.0221 | 0.9147 | -0.079 | 0.0348 | FALSE |
| All | Basic>Other | -0.039 | 0 | -0.0589 | -0.0191 | TRUE |
| All | Other>Acidic | 0.0739 | 0 | 0.052 | 0.0957 | TRUE |
| All | Other>Basic | 0.0469 | 0 | 0.0249 | 0.0689 | TRUE |
| Basic->Acidic | Basic>Other | -0.017 | 0.9808 | -0.0766 | 0.0426 | FALSE |
| Basic->Acidic | Other>Acidic | 0.0959 | 0.0001 | 0.0357 | 0.1562 | TRUE |
| Basic->Acidic | Other>Basic | 0.069 | 0.0132 | 0.0086 | 0.1293 | TRUE |
| Basic>Other | Other>Acidic | 0.1129 | 0 | 0.0848 | 0.141 | TRUE |
| Basic>Other | Other>Basic | 0.086 | 0 | 0.0577 | 0.1142 | TRUE |
| Other>Acidic | Other>Basic | -0.0269 | 0.1032 | -0.0566 | 0.0027 | FALSE |

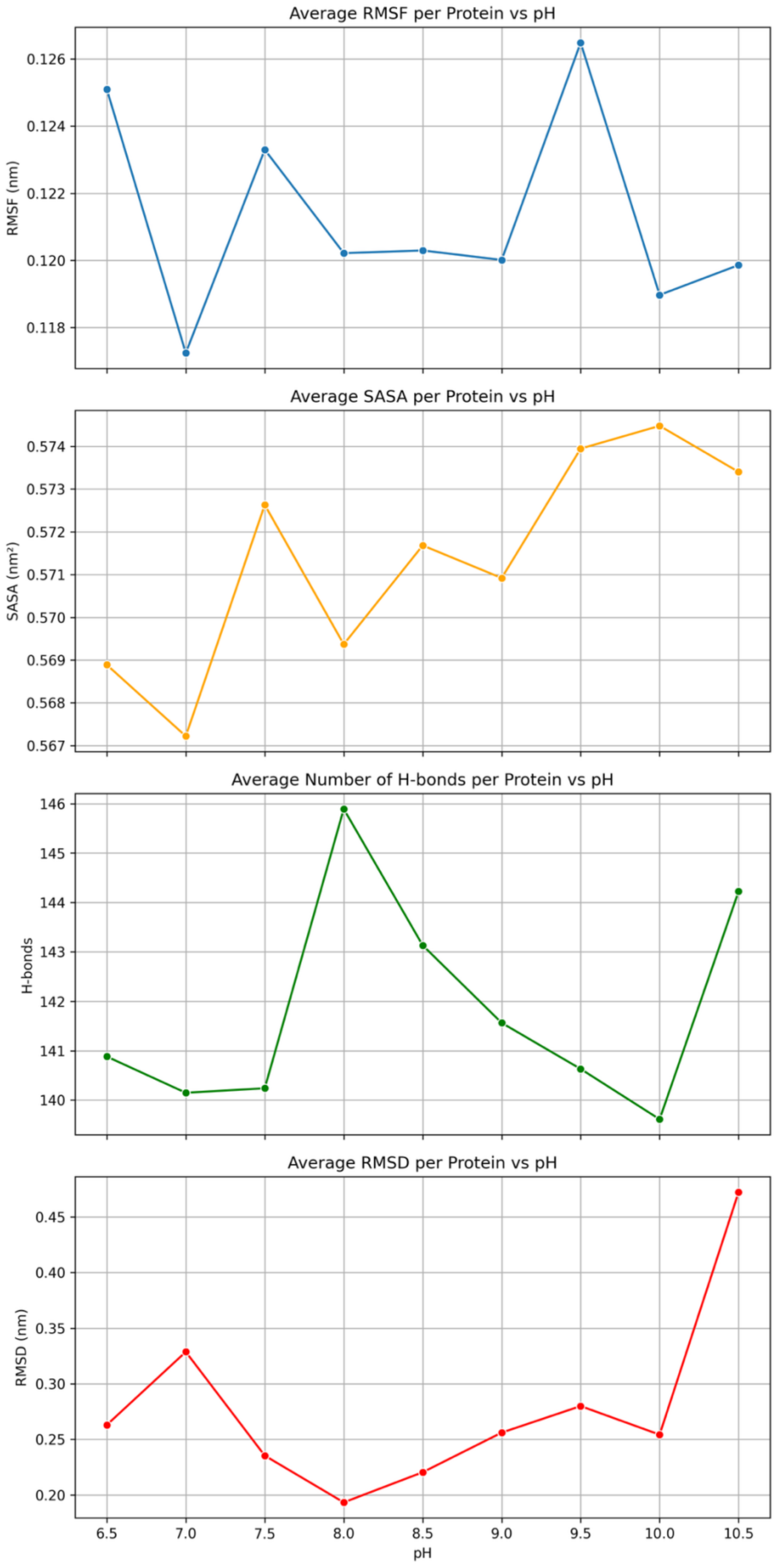

**Figure S4. Structural analysis of molecular dynamics simulations of JAM-A.**

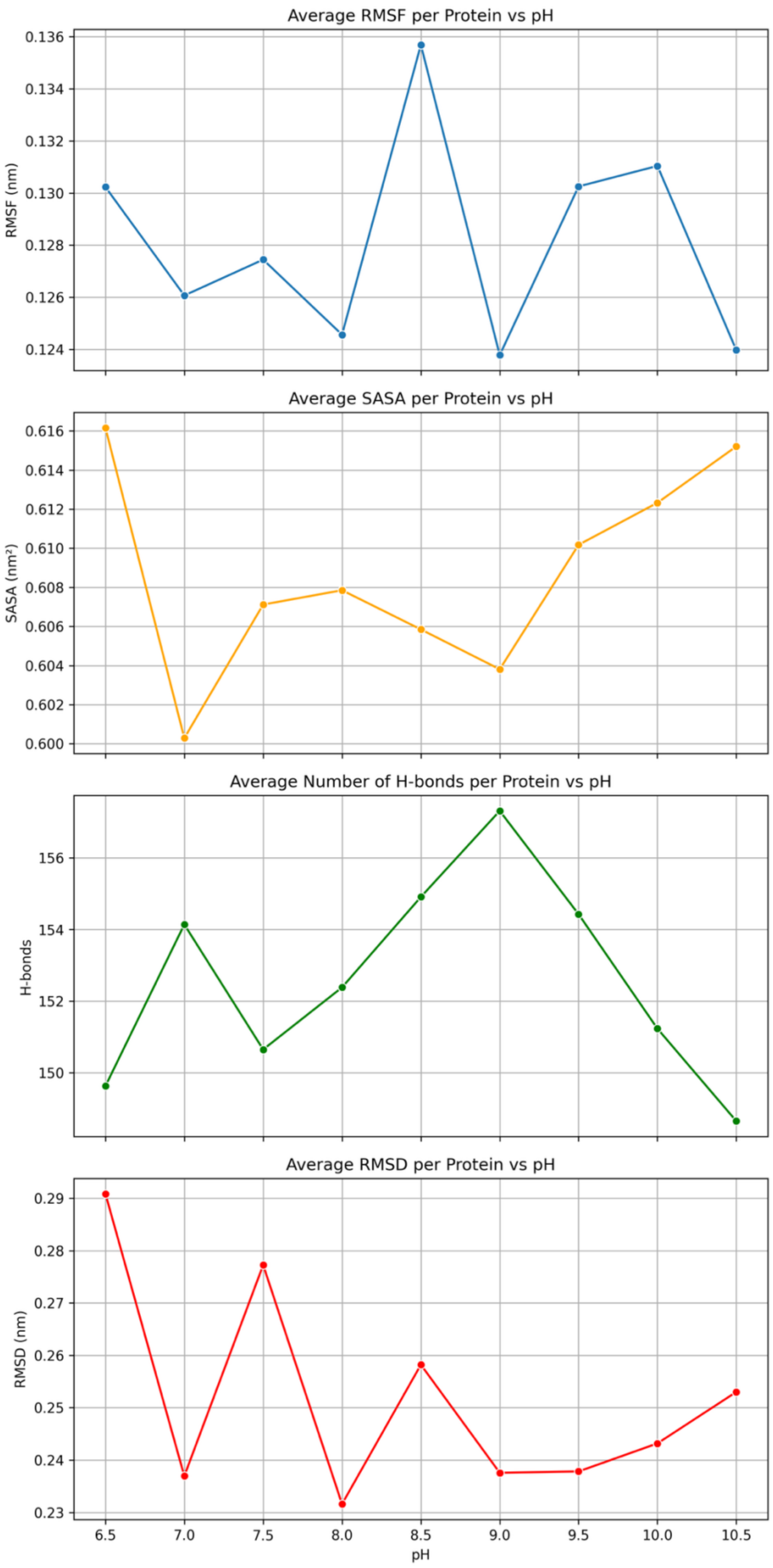

**Figure S5. Structural analysis of molecular dynamics simulations of JAM-B.**

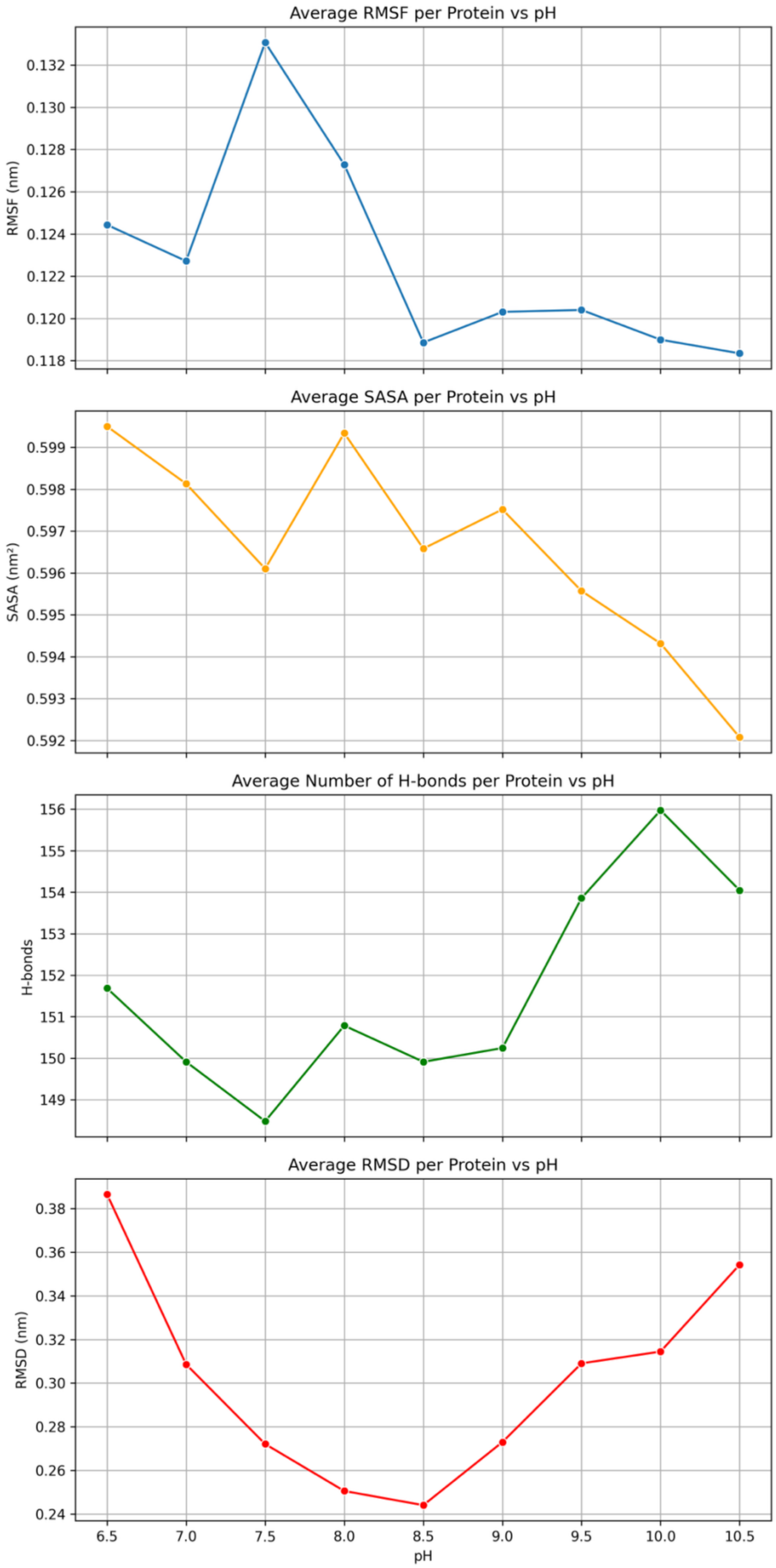

**Figure S6. Structural analysis of molecular dynamics simulations of JAM-C.**

**Table S12. pH-dependent dynamics and evolutionary conservation of JAM-A residues.**

| Residue | AA | Residue pI[1] | Residue Conserv.[2] | Avg. RMSF[3] | ΔRMSF[3] | Avg. SASA[4] | ΔSASA[4] |
|---|---|---|---|---|---|---|---|
| 30 | T | 5.6 | 1.19 | 0.18873833 | 0.049055 | 1.30162222 | 0.4045 |
| 31 | V | 5.96 | -0.753 | 0.12387278 | 0.0259 | 0.17083889 | 0.09905 |
| 32 | H | 7.95 | 0.417 | 0.15853333 | 0.03559 | 1.07177222 | 0.0951 |
| 33 | S | 5.68 | -0.352 | 0.11159278 | 0.0128 | 0.17498889 | 0.1221 |
| 34 | S | 5.68 | 0.388 | 0.14087111 | 0.024415 | 0.98342222 | 0.0852 |
| 35 | E | 3.22 | 0.879 | 0.15716444 | 0.035045 | 1.01872222 | 0.0793 |
| 36 | P | 6.3 | 2.332 | 0.12465667 | 0.01906 | 0.86212222 | 0.131 |
| 37 | E | 3.22 | 1.651 | 0.13338833 | 0.02664 | 0.77260556 | 0.17215 |
| 38 | V | 5.96 | -0.06 | 0.09333389 | 0.01394 | 0.1767 | 0.0835 |
| 39 | R | 10.76 | 1.044 | 0.12545667 | 0.024815 | 1.36355 | 0.07775 |
| 40 | I | 6.02 | -1.014 | 0.09727556 | 0.0224 | 0.33206111 | 0.0664 |
| 41 | P | 6.3 | 0.672 | 0.10451778 | 0.02857 | 0.41767778 | 0.1198 |
| 42 | E | 3.22 | -1.231 | 0.12230333 | 0.035595 | 0.57952778 | 0.2573 |
| 43 | N | 5.41 | -0.899 | 0.12846667 | 0.015915 | 1.11488889 | 0.0917 |
| 44 | N | 5.41 | 0.829 | 0.10841611 | 0.01741 | 0.5712 | 0.0441 |
| 45 | P | 6.3 | 0.852 | 0.09585667 | 0.01697 | 0.92086667 | 0.02625 |
| 46 | V | 5.96 | 0.109 | 0.08079556 | 0.01392 | 0.1384 | 0.02745 |
| 47 | K | 9.74 | -0.38 | 0.12139167 | 0.01914 | 1.22937222 | 0.1331 |
| 48 | L | 5.98 | -0.866 | 0.07717389 | 0.0064 | 0.00348889 | 0.0048 |
| 49 | S | 5.68 | 0.706 | 0.09261556 | 0.01665 | 0.26371111 | 0.13305 |
| 50 | C | 5.07 | -1.357 | 0.09465611 | 0.01548 | 0.02895 | 0.00935 |
| 51 | A | 6 | 1.237 | 0.11238389 | 0.02305 | 0.44391111 | 0.1454 |
| 52 | Y | 5.66 | -0.443 | 0.11622 | 0.018755 | 0.53938333 | 0.0631 |
| 53 | S | 5.68 | -0.36 | 0.15823333 | 0.03493 | 0.87228889 | 0.19145 |
| 54 | G | 5.97 | 0.377 | 0.16607444 | 0.03102 | 0.75855556 | 0.04465 |
| 55 | F | 5.48 | -0.606 | 0.14420611 | 0.016125 | 0.32569444 | 0.07825 |
| 56 | S | 5.68 | 0.639 | 0.176615 | 0.0227 | 0.98843333 | 0.2992 |
| 57 | S | 5.68 | 0.804 | 0.16154 | 0.01677 | 0.86735556 | 0.26185 |
| 58 | P | 6.3 | 0.756 | 0.12908889 | 0.01376 | 0.40867778 | 0.0743 |
| 59 | R | 10.76 | -1.271 | 0.11922222 | 0.012645 | 0.61639444 | 0.28745 |
| 60 | V | 5.96 | 0.061 | 0.08917611 | 0.01003 | 0.01802778 | 0.0152 |
| 61 | E | 3.22 | -1.266 | 0.09927167 | 0.009995 | 0.19041667 | 0.04985 |
| 62 | W | 5.89 | -1.266 | 0.062035 | 0.012665 | 0.00193333 | 0.0031 |
| 63 | K | 9.74 | -1.169 | 0.11082333 | 0.022105 | 0.61966667 | 0.07395 |
| 64 | F | 5.48 | -0.651 | 0.09620111 | 0.019465 | 0.08905556 | 0.08065 |
| 65 | D | 2.77 | 0.246 | 0.13611722 | 0.02363 | 0.51568333 | 0.24935 |
| 66 | Q | 5.65 | 3.006 | 0.1652 | 0.07057 | 0.81787222 | 0.34095 |
| 67 | G | 5.97 | 0.819 | 0.19369667 | 0.05655 | 0.70286667 | 0.2851 |
| 68 | D | 2.77 | 0.667 | 0.24268722 | 0.05535 | 1.53906667 | 0.38655 |
| 69 | T | 5.6 | -0.164 | 0.20126222 | 0.049855 | 0.85715556 | 0.23715 |
| 70 | T | 5.6 | -0.193 | 0.15011889 | 0.03674 | 0.76203333 | 0.1951 |
| 71 | R | 10.76 | 0.869 | 0.15421222 | 0.03294 | 1.19798889 | 0.26815 |

| | | | | | | | |
|---|---|---|---|---|---|---|---|
| 72 | L | 5.98 | -0.264 | 0.11700611 | 0.020845 | 0.8015 | 0.10385 |
| 73 | V | 5.96 | -0.673 | 0.07867889 | 0.017985 | 0.00075556 | 0.002 |
| 74 | C | 5.07 | 0.174 | 0.07748556 | 0.0139 | 0.00037778 | 0.00065 |
| 75 | Y | 5.66 | 0.047 | 0.13242389 | 0.01572 | 1.27531111 | 0.11415 |
| 76 | N | 5.41 | 0.48 | 0.12516111 | 0.01373 | 0.7193 | 0.08405 |
| 77 | N | 5.41 | 0.933 | 0.11102333 | 0.01075 | 0.56170556 | 0.05475 |
| 78 | K | 9.74 | 0.398 | 0.15701944 | 0.024005 | 1.57058333 | 0.17685 |
| 79 | I | 6.02 | 0.467 | 0.09235333 | 0.018145 | 0.20987222 | 0.05335 |
| 80 | T | 5.6 | -1.039 | 0.10695722 | 0.02408 | 0.29746667 | 0.0737 |
| 81 | A | 6 | 1.503 | 0.13471167 | 0.037755 | 0.93664444 | 0.1284 |
| 82 | S | 5.68 | 1.5 | 0.13379722 | 0.02985 | 1.00855556 | 0.06165 |
| 83 | Y | 5.66 | -0.955 | 0.09592722 | 0.013275 | 0.18976667 | 0.1104 |
| 84 | E | 3.22 | 0.966 | 0.14530556 | 0.01579 | 1.09263333 | 0.07 |
| 85 | D | 2.77 | 1.917 | 0.15221944 | 0.01737 | 1.35435556 | 0.08395 |
| 86 | R | 10.76 | -1.274 | 0.09616222 | 0.01716 | 0.69805 | 0.0774 |
| 87 | V | 5.96 | -0.618 | 0.08418722 | 0.00971 | 0.03516667 | 0.01425 |
| 88 | T | 5.6 | 0.497 | 0.10677778 | 0.009835 | 0.68335 | 0.05455 |
| 89 | F | 5.48 | 0.744 | 0.09652444 | 0.012525 | 0.53763333 | 0.0675 |
| 90 | L | 5.98 | 0.291 | 0.12489889 | 0.02377 | 0.85091667 | 0.2799 |
| 91 | P | 6.3 | 2.635 | 0.12109167 | 0.0227 | 0.85868889 | 0.21055 |
| 92 | T | 5.6 | 0.756 | 0.11664778 | 0.027385 | 0.74565556 | 0.11405 |
| 93 | G | 5.97 | -0.302 | 0.08128778 | 0.011205 | 0.01642222 | 0.01955 |
| 94 | I | 6.02 | -0.975 | 0.06670278 | 0.005635 | 0.00082778 | 0.0024 |
| 95 | T | 5.6 | -0.048 | 0.08228056 | 0.009475 | 0.32650556 | 0.04535 |
| 96 | F | 5.48 | -0.618 | 0.08638722 | 0.02024 | 0.00941111 | 0.01555 |
| 97 | K | 9.74 | 1.175 | 0.13844556 | 0.015045 | 1.3367 | 0.24765 |
| 98 | S | 5.68 | -0.26 | 0.10239111 | 0.015985 | 0.50482778 | 0.1301 |
| 99 | V | 5.96 | -0.856 | 0.09158167 | 0.01192 | 0.00173333 | 0.00295 |
| 100 | T | 5.6 | -1.049 | 0.10564167 | 0.01246 | 0.53083889 | 0.0506 |
| 101 | R | 10.76 | -0.978 | 0.11773667 | 0.017285 | 0.99915556 | 0.0411 |
| 102 | E | 3.22 | 0.623 | 0.150365 | 0.015395 | 1.45802222 | 0.0841 |
| 103 | D | 2.77 | -1.371 | 0.08724556 | 0.010565 | 0.07922778 | 0.0376 |
| 104 | T | 5.6 | -0.389 | 0.10610556 | 0.013845 | 0.3934 | 0.2082 |
| 105 | G | 5.97 | -1.286 | 0.09750222 | 0.0119 | 0.10727778 | 0.08565 |
| 106 | T | 5.6 | 1.326 | 0.10810389 | 0.02201 | 0.36278333 | 0.1067 |
| 107 | Y | 5.66 | -1.272 | 0.07412167 | 0.01343 | 0.00470556 | 0.00725 |
| 108 | T | 5.6 | 0.055 | 0.08989556 | 0.017925 | 0.03423333 | 0.0254 |
| 109 | C | 5.07 | -1.358 | 0.07466056 | 0.011405 | 0.00073333 | 0.00185 |
| 110 | M | 5.74 | -0.872 | 0.09428333 | 0.01869 | 0.20673889 | 0.0405 |
| 111 | V | 5.96 | -1.249 | 0.09952556 | 0.01245 | 0.00022778 | 0.0004 |
| 112 | S | 5.68 | -0.731 | 0.11388833 | 0.00997 | 0.09992778 | 0.0259 |
| 113 | E | 3.22 | 0.155 | 0.16041222 | 0.020455 | 0.35511111 | 0.15615 |
| 114 | E | 3.22 | 1.697 | 0.20878667 | 0.036455 | 1.23980556 | 0.2694 |
| 115 | G | 5.97 | 0.863 | 0.19184167 | 0.026795 | 0.80206111 | 0.17315 |
| 116 | G | 5.97 | 0.914 | 0.18078444 | 0.02001 | 0.59220556 | 0.1942 |

| | | | | | | | |
|---|---|---|---|---|---|---|---|
| 117 | N | 5.41 | 0.874 | 0.19316222 | 0.039525 | 1.03694444 | 0.4338 |
| 118 | S | 5.68 | 1.257 | 0.15735389 | 0.010625 | 0.67008889 | 0.29255 |
| 119 | Y | 5.66 | 0.161 | 0.13642389 | 0.016845 | 1.33506111 | 0.1022 |
| 120 | G | 5.97 | 1.377 | 0.11459 | 0.01595 | 0.11871667 | 0.042 |
| 121 | E | 3.22 | -0.635 | 0.13523667 | 0.019115 | 0.77317222 | 0.11685 |
| 122 | V | 5.96 | 0.629 | 0.10520556 | 0.019215 | 0.23677778 | 0.03355 |
| 123 | K | 9.74 | 1.064 | 0.13290611 | 0.01628 | 1.17985 | 0.10505 |
| 124 | V | 5.96 | -0.699 | 0.08949111 | 0.01163 | 0.00326667 | 0.00945 |
| 125 | K | 9.74 | 1.276 | 0.14050944 | 0.028525 | 1.00577778 | 0.0923 |
| 126 | L | 5.98 | -1.089 | 0.08677056 | 0.012765 | 0.00241111 | 0.00245 |
| 127 | I | 6.02 | 0.368 | 0.11115056 | 0.00997 | 0.22668333 | 0.1999 |
| 128 | V | 5.96 | -1.381 | 0.09881056 | 0.015555 | 0.00656111 | 0.0429 |
| 129 | L | 5.98 | -0.743 | 0.10983056 | 0.028095 | 0.19851667 | 0.1358 |
| 130 | V | 5.96 | -1.411 | 0.10806667 | 0.029085 | 0.09613333 | 0.0389 |
| 131 | P | 6.3 | -0.681 | 0.10102611 | 0.021585 | 0.57585 | 0.18195 |
| 132 | P | 6.3 | -1.165 | 0.091205 | 0.016405 | 0.062 | 0.01615 |
| 133 | S | 5.68 | -0.77 | 0.09234 | 0.014355 | 0.3205 | 0.0505 |
| 134 | K | 9.74 | 0.374 | 0.13237778 | 0.01996 | 1.3037 | 0.1179 |
| 135 | P | 6.3 | -1.377 | 0.08553167 | 0.012915 | 0.03306667 | 0.01515 |
| 136 | T | 5.6 | 0.721 | 0.11985167 | 0.018655 | 0.856 | 0.08645 |
| 137 | V | 5.96 | -0.718 | 0.09728 | 0.01955 | 0.1564 | 0.1065 |
| 138 | N | 5.41 | 0.516 | 0.13221278 | 0.03515 | 0.86711667 | 0.06035 |
| 139 | I | 6.02 | -0.465 | 0.13067667 | 0.056565 | 0.64422222 | 0.9684 |
| 140 | P | 6.3 | -1.377 | 0.13468556 | 0.032655 | 0.36203889 | 0.23015 |
| 141 | S | 5.68 | -0.097 | 0.194485 | 0.02155 | 1.20665 | 0.36005 |
| 142 | S | 5.68 | -1.075 | 0.19647611 | 0.036275 | 1.01963333 | 0.52495 |
| 143 | A | 6 | -0.792 | 0.15523444 | 0.028345 | 0.38082222 | 0.17205 |
| 144 | T | 5.6 | -0.888 | 0.17051444 | 0.02669 | 0.87216667 | 0.0927 |
| 145 | I | 6.02 | -0.987 | 0.15232 | 0.024205 | 1.01834444 | 0.0912 |
| 146 | G | 5.97 | -0.615 | 0.14856833 | 0.016995 | 0.50906111 | 0.0183 |
| 147 | N | 5.41 | 0.478 | 0.17180833 | 0.0139 | 0.89667222 | 0.0627 |
| 148 | R | 10.76 | 1.567 | 0.17014722 | 0.01768 | 1.61910556 | 0.12525 |
| 149 | A | 6 | 0.038 | 0.11067389 | 0.010725 | 0.04242222 | 0.0401 |
| 150 | V | 5.96 | 0.935 | 0.11622444 | 0.010025 | 0.74340556 | 0.25335 |
| 151 | L | 5.98 | -1.209 | 0.09513833 | 0.01879 | 0.00161111 | 0.00435 |
| 152 | T | 5.6 | 0.16 | 0.10883389 | 0.01254 | 0.49974444 | 0.0833 |
| 153 | C | 5.07 | -1.357 | 0.08678333 | 0.013205 | 0.0275 | 0.04 |
| 154 | S | 5.68 | 0.751 | 0.10334167 | 0.02095 | 0.47650556 | 0.16705 |
| 155 | E | 3.22 | -1.299 | 0.09823944 | 0.0147 | 0.14255 | 0.0391 |
| 156 | Q | 5.65 | 2.036 | 0.17996056 | 0.02631 | 1.6057 | 0.18095 |
| 157 | D | 2.77 | 0.454 | 0.13051556 | 0.01643 | 0.64745 | 0.11735 |
| 158 | G | 5.97 | -0.345 | 0.10974222 | 0.01813 | 0.11561667 | 0.02265 |
| 159 | S | 5.68 | -1.118 | 0.12046333 | 0.02546 | 0.22585556 | 0.33885 |
| 160 | P | 6.3 | -1.377 | 0.12129611 | 0.020035 | 0.31396667 | 0.218 |
| 161 | P | 6.3 | -0.204 | 0.12126444 | 0.02223 | 1.22685 | 0.0613 |

| | | | | | | | |
|---|---|---|---|---|---|---|---|
| 162 | S | 5.68 | -0.297 | 0.09361722 | 0.01687 | 0.04810556 | 0.02205 |
| 163 | E | 3.22 | -0.276 | 0.12737833 | 0.01807 | 0.88825556 | 0.26625 |
| 164 | Y | 5.66 | -0.894 | 0.07989556 | 0.011455 | 0.16982222 | 0.05195 |
| 165 | T | 5.6 | 1.155 | 0.07471889 | 0.01419 | 0.37880556 | 0.1452 |
| 166 | W | 5.89 | -1.354 | 0.06390944 | 0.00878 | 0.00553333 | 0.0039 |
| 167 | F | 5.48 | 0.391 | 0.08419833 | 0.01389 | 0.47438333 | 0.0917 |
| 168 | K | 9.74 | -0.529 | 0.09191278 | 0.013275 | 0.28823889 | 0.12925 |
| 169 | D | 2.77 | -0.134 | 0.11552111 | 0.031055 | 0.82952222 | 0.20245 |
| 170 | G | 5.97 | 1.091 | 0.11495389 | 0.023865 | 0.67315556 | 0.09205 |
| 171 | I | 6.02 | 0.819 | 0.12308944 | 0.020715 | 1.02293889 | 0.11715 |
| 172 | V | 5.96 | 1.513 | 0.09921833 | 0.013415 | 0.79805556 | 0.0474 |
| 173 | M | 5.74 | -0.768 | 0.0796 | 0.01301 | 0.06783333 | 0.02405 |
| 174 | P | 6.3 | -0.64 | 0.10855056 | 0.020235 | 0.40773889 | 0.0811 |
| 175 | T | 5.6 | 1.671 | 0.13427611 | 0.016125 | 0.78505556 | 0.12455 |
| 176 | N | 5.41 | 0.13 | 0.15993222 | 0.0318 | 0.91406111 | 0.12115 |
| 177 | P | 6.3 | -1.032 | 0.14018833 | 0.035315 | 0.17193333 | 0.1572 |
| 178 | K | 9.74 | -0.493 | 0.21296111 | 0.06564 | 1.59285556 | 0.3118 |
| 179 | S | 5.68 | 0.725 | 0.199095 | 0.05335 | 0.98442222 | 0.42395 |
| 180 | T | 5.6 | 0.06 | 0.1728 | 0.00621 | 0.49045556 | 0.17505 |
| 181 | R | 10.76 | 0.399 | 0.25744222 | 0.04243 | 2.35066111 | 0.18185 |
| 182 | A | 6 | 1.134 | 0.16017611 | 0.020415 | 0.75597778 | 0.23775 |
| 183 | F | 5.48 | -0.644 | 0.12557722 | 0.023925 | 0.20197778 | 0.188 |
| 184 | S | 5.68 | 0.078 | 0.16567778 | 0.032345 | 0.53281111 | 0.3455 |
| 185 | N | 5.41 | -1.372 | 0.17374944 | 0.033565 | 1.38673889 | 0.1035 |
| 186 | S | 5.68 | -0.459 | 0.11523778 | 0.020435 | 0.11723333 | 0.0543 |
| 187 | S | 5.68 | -1.248 | 0.12448778 | 0.024465 | 0.5066 | 0.1455 |
| 188 | Y | 5.66 | -1.274 | 0.09844056 | 0.02454 | 0.11533333 | 0.0464 |
| 189 | V | 5.96 | 0.51 | 0.12349556 | 0.02152 | 0.94258889 | 0.1008 |
| 190 | L | 5.98 | 0.261 | 0.10683833 | 0.01778 | 0.13183333 | 0.03545 |
| 191 | N | 5.41 | -0.723 | 0.12517222 | 0.028215 | 0.64320556 | 0.01445 |
| 192 | P | 6.3 | 2.844 | 0.10687889 | 0.020545 | 0.44456111 | 0.0997 |
| 193 | T | 5.6 | 1.477 | 0.13722778 | 0.016855 | 0.95346667 | 0.1248 |
| 194 | T | 5.6 | 0.055 | 0.12341389 | 0.01725 | 0.78160556 | 0.13595 |
| 195 | G | 5.97 | -1.28 | 0.09388889 | 0.01409 | 0.00045 | 0.0015 |
| 196 | E | 3.22 | -0.038 | 0.13007889 | 0.014525 | 0.73883333 | 0.06255 |
| 197 | L | 5.98 | -1.337 | 0.08322944 | 0.011555 | 0.00021667 | 0.0016 |
| 198 | V | 5.96 | 0.718 | 0.10943222 | 0.0128 | 0.31115556 | 0.05465 |
| 199 | F | 5.48 | -1.208 | 0.09587611 | 0.013 | 0.00757222 | 0.00635 |
| 200 | D | 2.77 | 0.273 | 0.14320389 | 0.01476 | 0.55235556 | 0.1059 |
| 201 | P | 6.3 | 0.642 | 0.13210278 | 0.014745 | 0.71490556 | 0.0444 |
| 202 | L | 5.98 | 0.101 | 0.11449722 | 0.00924 | 0.03315 | 0.0492 |
| 203 | S | 5.68 | 0.953 | 0.11807333 | 0.020145 | 0.35929444 | 0.0427 |
| 204 | A | 6 | -0.43 | 0.137795 | 0.025935 | 0.7585 | 0.0545 |
| 205 | S | 5.68 | 2.02 | 0.12998 | 0.021825 | 0.84419444 | 0.14255 |
| 206 | D | 2.77 | -1.374 | 0.09873944 | 0.01209 | 0.04065556 | 0.0275 |

| | | | | | | | |
|---|---|---|---|---|---|---|---|
| 207 | T | 5.6 | 0.122 | 0.11937389 | 0.01997 | 0.59522222 | 0.1616 |
| 208 | G | 5.97 | -0.86 | 0.10097333 | 0.01569 | 0.10806111 | 0.1059 |
| 209 | E | 3.22 | 0.245 | 0.12700333 | 0.025485 | 0.81212222 | 0.35685 |
| 210 | Y | 5.66 | -1.275 | 0.07490278 | 0.012015 | 0.00644444 | 0.01915 |
| 211 | S | 5.68 | 1.045 | 0.06753333 | 0.007665 | 0.17923333 | 0.06675 |
| 212 | C | 5.07 | -1.358 | 0.06484444 | 0.00947 | 0.00045 | 0.00105 |
| 213 | E | 3.22 | -0.656 | 0.08223111 | 0.00919 | 0.34405556 | 0.11165 |
| 214 | A | 6 | -0.967 | 0.07012778 | 0.014075 | 2.78E-05 | 0.00015 |
| 215 | R | 10.76 | 1.383 | 0.12016833 | 0.039115 | 1.14217778 | 0.2208 |
| 216 | N | 5.41 | -1.421 | 0.09845778 | 0.01722 | 0.23626667 | 0.0477 |
| 217 | G | 5.97 | 0.906 | 0.11083611 | 0.01945 | 0.72207778 | 0.17065 |
| 218 | Y | 5.66 | -0.272 | 0.10998722 | 0.039675 | 0.87389444 | 0.0454 |
| 219 | G | 5.97 | -1.191 | 0.10422333 | 0.02922 | 0.49466667 | 0.09395 |
| 220 | T | 5.6 | 1.816 | 0.12017611 | 0.025105 | 1.05187778 | 0.0231 |
| 221 | P | 6.3 | 0.826 | 0.09147278 | 0.02381 | 0.56476667 | 0.1318 |
| 222 | M | 5.74 | -0.463 | 0.09941444 | 0.012105 | 0.63864444 | 0.0387 |
| 223 | T | 5.6 | 0.551 | 0.09707444 | 0.006905 | 0.68978333 | 0.05355 |
| 224 | S | 5.68 | -0.945 | 0.07928944 | 0.013315 | 0.10394444 | 0.0521 |
| 225 | N | 5.41 | 1.035 | 0.13035111 | 0.017955 | 1.18357778 | 0.2 |
| 226 | A | 6 | 0.672 | 0.10683056 | 0.010425 | 0.60744444 | 0.14915 |
| 227 | V | 5.96 | 1.066 | 0.10786333 | 0.0137 | 0.39117778 | 0.3497 |
| 228 | R | 10.76 | 1.129 | 0.15828389 | 0.0287 | 1.73051667 | 0.2203 |
| 229 | M | 5.74 | -1.254 | 0.12621 | 0.022405 | 0.40261667 | 0.2188 |

[1]pI of residue amino acid at 25 °C.
[2]Consurf evolutionary conservation score of the residue.
[3]The average root-mean-square fluctuation (RMSF) across all frames and steps, and the ΔRMSF (maximum minus minimum across pH).
[4]The average solvent accessibility surface area (SASA) across all frames and steps, and the ΔSASA (maximum minus minimum across pH).

**Table S13. pH-dependent dynamics and evolutionary conservation of JAM-B residues.**

| Residue | AA | Residue pI[1] | Residue Conserv.[2] | Avg. RMSF[3] | ΔRMSF[3] | Avg. SASA[4] | ΔSASA[4] |
|---|---|---|---|---|---|---|---|
| 36 | Q | 5.65 | 2.077 | 0.20769389 | 0.08033 | 1.53115 | 0.3528 |
| 37 | V | 5.96 | 1.568 | 0.14406667 | 0.021335 | 0.89574444 | 0.0755 |
| 38 | V | 5.96 | -0.943 | 0.11325222 | 0.01696 | 0.33897222 | 0.06865 |
| 39 | T | 5.6 | 0.662 | 0.13576889 | 0.023935 | 0.87035556 | 0.02225 |
| 40 | A | 6 | -0.892 | 0.12289722 | 0.02114 | 0.08730556 | 0.01245 |
| 41 | V | 5.96 | 0.82 | 0.13252444 | 0.02577 | 0.5242 | 0.0439 |
| 42 | E | 3.22 | -1.242 | 0.13201722 | 0.02666 | 0.26266111 | 0.04415 |
| 43 | Y | 5.66 | 0.271 | 0.15630444 | 0.020735 | 1.55289444 | 0.06975 |
| 44 | Q | 5.65 | 0.675 | 0.147145 | 0.0202 | 1.07194444 | 0.0439 |
| 45 | E | 3.22 | -0.151 | 0.12953111 | 0.024065 | 0.78243333 | 0.28485 |
| 46 | A | 6 | -0.333 | 0.09277167 | 0.007885 | 0.04746111 | 0.02365 |
| 47 | I | 6.02 | -0.275 | 0.09988444 | 0.008215 | 0.80608333 | 0.0481 |
| 48 | L | 5.98 | -1.176 | 0.08150278 | 0.009965 | 0.00168889 | 0.00205 |
| 49 | A | 6 | -0.655 | 0.08562333 | 0.010955 | 0.38947778 | 0.1222 |
| 50 | C | 5.07 | -1.413 | 0.09059 | 0.016775 | 0.10626667 | 0.0737 |
| 51 | K | 9.74 | 0.813 | 0.15558778 | 0.05041 | 1.57151111 | 0.24325 |
| 52 | T | 5.6 | -0.629 | 0.12434778 | 0.025065 | 0.23065 | 0.13245 |
| 53 | P | 6.3 | -0.598 | 0.16120056 | 0.029565 | 1.14746667 | 0.34085 |
| 54 | K | 9.74 | -0.662 | 0.17783944 | 0.023755 | 1.00003889 | 0.5337 |
| 55 | K | 9.74 | 0.238 | 0.20127278 | 0.09655 | 1.51768889 | 1.0954 |
| 56 | T | 5.6 | -0.27 | 0.17378389 | 0.042975 | 0.64583889 | 0.31175 |
| 57 | V | 5.96 | 1.569 | 0.20350111 | 0.05816 | 1.4403 | 0.42245 |
| 58 | S | 5.68 | 0.258 | 0.17756389 | 0.03856 | 0.88147778 | 0.26905 |
| 59 | S | 5.68 | -0.4 | 0.13204667 | 0.027645 | 0.17398889 | 0.31165 |
| 60 | R | 10.76 | -1.424 | 0.15349944 | 0.048335 | 1.14051667 | 0.4914 |
| 61 | L | 5.98 | -0.83 | 0.08654111 | 0.02218 | 0.08974444 | 0.06925 |
| 62 | E | 3.22 | -1.416 | 0.09983278 | 0.01733 | 0.26049444 | 0.2823 |
| 63 | W | 5.89 | -1.352 | 0.05493667 | 0.00563 | 0.00178333 | 0.00325 |
| 64 | K | 9.74 | -1.446 | 0.09740444 | 0.012775 | 0.31465556 | 0.07645 |
| 65 | K | 9.74 | -1.187 | 0.11698111 | 0.014775 | 0.47267778 | 0.0577 |
| 66 | L | 5.98 | 0.348 | 0.13394833 | 0.02637 | 0.54818333 | 0.26105 |
| 67 | G | 5.97 | 0.221 | 0.17171056 | 0.024705 | 0.51415556 | 0.06525 |
| 68 | R | 10.76 | 0.745 | 0.28448167 | 0.05027 | 2.70568889 | 0.1571 |
| 69 | S | 5.68 | 0.526 | 0.18946778 | 0.02565 | 1.00352222 | 0.1511 |
| 70 | V | 5.96 | -0.468 | 0.15309056 | 0.02225 | 0.81328889 | 0.08695 |
| 71 | S | 5.68 | -0.32 | 0.11717278 | 0.017085 | 0.56762778 | 0.08605 |
| 72 | F | 5.48 | -0.128 | 0.12360833 | 0.01355 | 0.82002222 | 0.0525 |
| 73 | V | 5.96 | -1.034 | 0.07540111 | 0.00924 | 0.00411667 | 0.00415 |
| 74 | Y | 5.66 | -0.267 | 0.08039111 | 0.01072 | 0.27429444 | 0.03575 |
| 75 | Y | 5.66 | 0.284 | 0.12624944 | 0.015715 | 0.83165 | 0.0677 |
| 76 | Q | 5.65 | 0.894 | 0.15821389 | 0.025675 | 1.25361667 | 0.1121 |
| 77 | Q | 5.65 | 0.783 | 0.14339722 | 0.031785 | 1.09502222 | 0.37575 |

| | | | | | | | |
|---|---|---|---|---|---|---|---|
| 78 | T | 5.6 | 1.673 | 0.11929444 | 0.014 | 0.71631667 | 0.0558 |
| 79 | L | 5.98 | 0.056 | 0.09798278 | 0.01693 | 0.26068333 | 0.0539 |
| 80 | Q | 5.65 | -0.3 | 0.11408722 | 0.01401 | 0.52205556 | 0.0528 |
| 81 | G | 5.97 | 0.42 | 0.12970056 | 0.027805 | 0.54498889 | 0.091 |
| 82 | D | 2.77 | 0.68 | 0.14855222 | 0.028915 | 1.06626667 | 0.27625 |
| 83 | F | 5.48 | -0.585 | 0.09765444 | 0.016315 | 0.11057222 | 0.04735 |
| 84 | K | 9.74 | 0.627 | 0.15401 | 0.03139 | 1.28431667 | 0.1063 |
| 85 | N | 5.41 | 0.934 | 0.15920389 | 0.023105 | 1.46011111 | 0.1568 |
| 86 | R | 10.76 | -1.426 | 0.10622222 | 0.01503 | 0.54646667 | 0.11255 |
| 87 | A | 6 | -1.071 | 0.08886667 | 0.014175 | 0.09215556 | 0.0469 |
| 88 | E | 3.22 | 0.927 | 0.126815 | 0.0137 | 0.77467778 | 0.23915 |
| 89 | M | 5.74 | 0.067 | 0.09034944 | 0.012485 | 0.50679444 | 0.0711 |
| 90 | I | 6.02 | 1.117 | 0.11802722 | 0.00867 | 0.83033889 | 0.14945 |
| 91 | D | 2.77 | 0.056 | 0.12585111 | 0.02104 | 1.10825556 | 0.26455 |
| 92 | F | 5.48 | -0.469 | 0.105855 | 0.02502 | 0.27259444 | 0.24145 |
| 93 | N | 5.41 | -0.611 | 0.08181056 | 0.007535 | 0.37573889 | 0.058 |
| 94 | I | 6.02 | -0.778 | 0.06916222 | 0.014765 | 0.00086111 | 0.00105 |
| 95 | R | 10.76 | 0.058 | 0.11043278 | 0.01442 | 0.69507778 | 0.0762 |
| 96 | I | 6.02 | -0.792 | 0.08780611 | 0.006405 | 0.0027 | 0.0033 |
| 97 | K | 9.74 | 1.634 | 0.12331667 | 0.00916 | 0.78436667 | 0.11855 |
| 98 | N | 5.41 | -0.641 | 0.13873278 | 0.01541 | 0.86207222 | 0.0805 |
| 99 | V | 5.96 | -0.851 | 0.10735556 | 0.0161 | 0.0006 | 0.00125 |
| 100 | T | 5.6 | -1.213 | 0.12150222 | 0.02191 | 0.41095 | 0.06865 |
| 101 | R | 10.76 | -1.157 | 0.13249111 | 0.030455 | 1.00988333 | 0.0554 |
| 102 | S | 5.68 | 0.506 | 0.12271833 | 0.023255 | 0.97226111 | 0.05 |
| 103 | D | 2.77 | -1.453 | 0.09934944 | 0.01383 | 0.08911667 | 0.03895 |
| 104 | A | 6 | -0.481 | 0.11115167 | 0.018445 | 0.34591667 | 0.0556 |
| 105 | G | 5.97 | -1.124 | 0.10511333 | 0.01949 | 0.23425556 | 0.01875 |
| 106 | K | 9.74 | 2.044 | 0.14478889 | 0.01969 | 1.19377778 | 0.0406 |
| 107 | Y | 5.66 | -1.366 | 0.07955333 | 0.008085 | 0.00434444 | 0.0042 |
| 108 | R | 10.76 | -0.775 | 0.11170889 | 0.01459 | 0.67871111 | 0.17775 |
| 109 | C | 5.07 | -1.417 | 0.06948056 | 0.01206 | 0.01062778 | 0.01855 |
| 110 | E | 3.22 | -1.278 | 0.100155 | 0.024125 | 0.26832222 | 0.15755 |
| 111 | V | 5.96 | -1.267 | 0.09442833 | 0.014075 | 0.03341667 | 0.0426 |
| 112 | S | 5.68 | -0.68 | 0.11768889 | 0.01896 | 0.18758333 | 0.1302 |
| 113 | A | 6 | -0.754 | 0.14620778 | 0.0167 | 0.12128333 | 0.1231 |
| 114 | P | 6.3 | 1.589 | 0.19180333 | 0.03279 | 0.82773889 | 0.3013 |
| 115 | S | 5.68 | 1.984 | 0.22713056 | 0.03169 | 0.53645556 | 0.1002 |
| 116 | E | 3.22 | -0.756 | 0.33919944 | 0.050985 | 1.81926111 | 0.05365 |
| 117 | Q | 5.65 | 0.969 | 0.32557167 | 0.03695 | 1.78007778 | 0.12555 |
| 118 | G | 5.97 | -0.685 | 0.23853778 | 0.02961 | 0.46742778 | 0.0994 |
| 119 | Q | 5.65 | -0.073 | 0.21769611 | 0.04487 | 1.27560556 | 0.3247 |
| 120 | N | 5.41 | 1.478 | 0.17790056 | 0.018275 | 0.77359444 | 0.22185 |
| 121 | L | 5.98 | 0.03 | 0.14401722 | 0.028125 | 0.99128333 | 0.19675 |
| 122 | E | 3.22 | 0.166 | 0.13839333 | 0.019505 | 0.67122222 | 0.2077 |

| | | | | | | | |
|---|---|---|---|---|---|---|---|
| 123 | E | 3.22 | -0.97 | 0.13303444 | 0.017545 | 1.00663333 | 0.0924 |
| 124 | D | 2.77 | -0.167 | 0.10545833 | 0.01856 | 0.3599 | 0.2481 |
| 125 | T | 5.6 | 0.745 | 0.10932778 | 0.019245 | 0.46276667 | 0.0697 |
| 126 | V | 5.96 | -0.768 | 0.09535389 | 0.011255 | 0.05631667 | 0.0292 |
| 127 | T | 5.6 | 0.526 | 0.11567056 | 0.01965 | 0.26749444 | 0.07925 |
| 128 | L | 5.98 | -1.13 | 0.10097444 | 0.01337 | 0.00952778 | 0.01095 |
| 129 | E | 3.22 | 0.338 | 0.14978944 | 0.030555 | 0.81147778 | 0.07155 |
| 130 | V | 5.96 | -1.233 | 0.12072111 | 0.022455 | 0.07612778 | 0.0186 |
| 131 | L | 5.98 | -0.635 | 0.14038222 | 0.030365 | 0.37228333 | 0.0715 |
| 132 | V | 5.96 | -1.459 | 0.12987278 | 0.02636 | 0.05508333 | 0.04085 |
| 133 | A | 6 | -0.819 | 0.12214056 | 0.022675 | 0.28141667 | 0.10395 |
| 134 | P | 6.3 | -1.219 | 0.10947667 | 0.014485 | 0.02854444 | 0.016 |
| 135 | A | 6 | -0.456 | 0.12034333 | 0.020265 | 0.37298889 | 0.1179 |
| 136 | V | 5.96 | 0.575 | 0.13140167 | 0.029385 | 1.00653333 | 0.18555 |
| 137 | P | 6.3 | -1.435 | 0.09889833 | 0.01358 | 0.06985 | 0.0379 |
| 138 | S | 5.68 | 0.32 | 0.11011722 | 0.010055 | 0.69123333 | 0.09405 |
| 139 | C | 5.07 | -0.949 | 0.09195333 | 0.01551 | 0.12612222 | 0.036 |
| 140 | E | 3.22 | 0.462 | 0.14103444 | 0.023475 | 1.10547778 | 0.24485 |
| 141 | V | 5.96 | -0.796 | 0.094245 | 0.01685 | 0.19271111 | 0.0539 |
| 142 | P | 6.3 | -1.435 | 0.10909278 | 0.015425 | 0.47546111 | 0.0294 |
| 143 | S | 5.68 | 0.056 | 0.14446722 | 0.02733 | 0.98515556 | 0.0993 |
| 144 | S | 5.68 | -0.927 | 0.14653 | 0.02488 | 0.73352778 | 0.17015 |
| 145 | A | 6 | -0.972 | 0.13234444 | 0.022955 | 0.10682778 | 0.09235 |
| 146 | L | 5.98 | -0.911 | 0.17405833 | 0.030945 | 1.24892778 | 0.212 |
| 147 | S | 5.68 | -0.904 | 0.15221056 | 0.03328 | 0.54635 | 0.2456 |
| 148 | G | 5.97 | -1.31 | 0.15246889 | 0.02962 | 0.51638333 | 0.177 |
| 149 | T | 5.6 | -0.193 | 0.14722278 | 0.0273 | 0.57840556 | 0.125 |
| 150 | V | 5.96 | 0.866 | 0.135165 | 0.028095 | 0.91817778 | 0.1533 |
| 151 | V | 5.96 | -0.804 | 0.10728722 | 0.024415 | 0.10459444 | 0.0595 |
| 152 | E | 3.22 | 0.119 | 0.130165 | 0.023725 | 0.75907222 | 0.28255 |
| 153 | L | 5.98 | -1.133 | 0.08613333 | 0.00893 | 0.00258333 | 0.0055 |
| 154 | R | 10.76 | 0.362 | 0.138725 | 0.04683 | 1.14356667 | 0.21415 |
| 155 | C | 5.07 | -1.417 | 0.08339278 | 0.01151 | 0.02526667 | 0.03335 |
| 156 | Q | 5.65 | 1.074 | 0.14098889 | 0.02611 | 0.91062222 | 0.25055 |
| 157 | D | 2.77 | -1.248 | 0.11191778 | 0.01581 | 0.17175 | 0.1319 |
| 158 | K | 9.74 | 1.083 | 0.18074944 | 0.027405 | 1.92222778 | 0.07755 |
| 159 | E | 3.22 | -0.273 | 0.13594333 | 0.02107 | 0.70636111 | 0.1834 |
| 160 | G | 5.97 | -1.004 | 0.11863222 | 0.01827 | 0.13908889 | 0.01485 |
| 161 | N | 5.41 | -0.425 | 0.14844889 | 0.02789 | 0.72872222 | 0.1331 |
| 162 | P | 6.3 | -1.435 | 0.13300111 | 0.027155 | 0.65921111 | 0.05045 |
| 163 | A | 6 | 0.207 | 0.12247 | 0.01686 | 0.876 | 0.06935 |
| 164 | P | 6.3 | -0.614 | 0.10649667 | 0.0127 | 0.08914444 | 0.0214 |
| 165 | E | 3.22 | 0.117 | 0.131565 | 0.010615 | 0.78040556 | 0.1496 |
| 166 | Y | 5.66 | -1.164 | 0.08366833 | 0.01179 | 0.13462222 | 0.0508 |
| 167 | T | 5.6 | 0.67 | 0.07486333 | 0.014995 | 0.25620556 | 0.1415 |

| | | | | | | | |
|---|---|---|---|---|---|---|---|
| 168 | W | 5.89 | -1.355 | 0.05989056 | 0.00862 | 0.00441667 | 0.00745 |
| 169 | F | 5.48 | -0.035 | 0.08018944 | 0.009045 | 0.23192778 | 0.05455 |
| 170 | K | 9.74 | -0.686 | 0.08374056 | 0.01332 | 0.10285 | 0.16475 |
| 171 | D | 2.77 | -0.804 | 0.10831778 | 0.01033 | 0.73293333 | 0.06805 |
| 172 | G | 5.97 | 0.122 | 0.11276667 | 0.007305 | 0.62495 | 0.0619 |
| 173 | I | 6.02 | 0.934 | 0.11928056 | 0.014185 | 0.97643889 | 0.3919 |
| 174 | R | 10.76 | 1.223 | 0.12653278 | 0.042215 | 1.30344444 | 0.38025 |
| 175 | L | 5.98 | -0.481 | 0.08636556 | 0.00983 | 0.03052778 | 0.02735 |
| 176 | L | 5.98 | 0.512 | 0.12612944 | 0.017335 | 0.68288889 | 0.2625 |
| 177 | E | 3.22 | 2.917 | 0.14745 | 0.02691 | 0.87772222 | 0.3882 |
| 178 | N | 5.41 | 0.083 | 0.16262111 | 0.02974 | 0.8097 | 0.1719 |
| 179 | P | 6.3 | -0.369 | 0.16580167 | 0.04493 | 0.50322222 | 0.42565 |
| 180 | R | 10.76 | 0.303 | 0.25600778 | 0.069395 | 2.16596667 | 0.19565 |
| 181 | L | 5.98 | 0.951 | 0.20805833 | 0.097485 | 1.40893333 | 0.65755 |
| 182 | G | 5.97 | 1.074 | 0.16915389 | 0.03321 | 0.38317778 | 0.2051 |
| 183 | S | 5.68 | 1.403 | 0.13952111 | 0.025775 | 0.30680556 | 0.2211 |
| 184 | Q | 5.65 | 0.979 | 0.19857056 | 0.037635 | 1.74611667 | 0.3028 |
| 185 | S | 5.68 | 0.523 | 0.12413944 | 0.03469 | 0.29263889 | 0.20375 |
| 186 | T | 5.6 | 1.227 | 0.10526056 | 0.013725 | 0.10062222 | 0.08835 |
| 187 | N | 5.41 | -1.301 | 0.14408111 | 0.03555 | 1.11103889 | 0.43235 |
| 188 | S | 5.68 | -0.418 | 0.10684722 | 0.022245 | 0.13471111 | 0.13555 |
| 189 | S | 5.68 | -0.791 | 0.11811944 | 0.015165 | 0.68101111 | 0.32205 |
| 190 | Y | 5.66 | -1.095 | 0.09256389 | 0.008935 | 0.14757778 | 0.1336 |
| 191 | T | 5.6 | 0.253 | 0.11886667 | 0.01848 | 0.81841667 | 0.0722 |
| 192 | M | 5.74 | 1.346 | 0.10413944 | 0.02096 | 0.21735 | 0.08125 |
| 193 | N | 5.41 | -0.528 | 0.11399889 | 0.012995 | 0.71282778 | 0.03035 |
| 194 | T | 5.6 | 2.626 | 0.10359 | 0.0091 | 0.28487222 | 0.1365 |
| 195 | K | 9.74 | 2.36 | 0.14527444 | 0.01583 | 1.50882778 | 0.17995 |
| 196 | T | 5.6 | -0.424 | 0.11796389 | 0.01927 | 0.82234444 | 0.09405 |
| 197 | G | 5.97 | -1.37 | 0.08418167 | 0.01213 | 0.00091111 | 0.00165 |
| 198 | T | 5.6 | 0.102 | 0.09973889 | 0.022155 | 0.29437778 | 0.27585 |
| 199 | L | 5.98 | -1.315 | 0.07520944 | 0.013865 | 0.00187778 | 0.00485 |
| 200 | Q | 5.65 | 0.675 | 0.13099389 | 0.018755 | 0.60105 | 0.07235 |
| 201 | F | 5.48 | -0.986 | 0.09117278 | 0.016705 | 0.00661667 | 0.0112 |
| 202 | N | 5.41 | 0.482 | 0.14466222 | 0.02847 | 0.78017222 | 0.3558 |
| 203 | T | 5.6 | 0.412 | 0.13656778 | 0.03105 | 0.74069444 | 0.2119 |
| 204 | V | 5.96 | -0.774 | 0.11643222 | 0.02463 | 0.02856667 | 0.0323 |
| 205 | S | 5.68 | 0.19 | 0.12279667 | 0.026015 | 0.33919444 | 0.1938 |
| 206 | K | 9.74 | -0.693 | 0.16625889 | 0.028415 | 1.35542222 | 0.0827 |
| 207 | L | 5.98 | 1.606 | 0.15578444 | 0.016855 | 1.34012778 | 0.30535 |
| 208 | D | 2.77 | -1.388 | 0.10251778 | 0.02392 | 0.01845 | 0.0756 |
| 209 | T | 5.6 | 0.128 | 0.11622944 | 0.012175 | 0.38489444 | 0.2392 |
| 210 | G | 5.97 | -1.064 | 0.09744833 | 0.01302 | 0.05416111 | 0.0854 |
| 211 | E | 3.22 | 0.335 | 0.12137889 | 0.009205 | 0.72193333 | 0.19835 |
| 212 | Y | 5.66 | -1.366 | 0.06840222 | 0.00954 | 0.00560556 | 0.01955 |

| | | | | | | | |
|---|---|---|---|---|---|---|---|
| 213 | S | 5.68 | 0.839 | 0.06889278 | 0.006335 | 0.13946111 | 0.04275 |
| 214 | C | 5.07 | -1.417 | 0.06186278 | 0.003795 | 0.00078889 | 0.00365 |
| 215 | E | 3.22 | -0.354 | 0.08527778 | 0.00953 | 0.09368889 | 0.0666 |
| 216 | A | 6 | -1.338 | 0.08181778 | 0.01168 | 0.00073333 | 0.00245 |
| 217 | R | 10.76 | 1.201 | 0.10808833 | 0.03289 | 1.10503889 | 0.11485 |
| 218 | N | 5.41 | -1.46 | 0.111175 | 0.02734 | 0.17926111 | 0.043 |
| 219 | S | 5.68 | 1.249 | 0.13968889 | 0.030675 | 1.09525 | 0.06315 |
| 220 | V | 5.96 | -0.647 | 0.13606833 | 0.03496 | 0.43792778 | 0.0428 |
| 221 | G | 5.97 | -1.427 | 0.11281778 | 0.02632 | 0.15546111 | 0.0239 |
| 222 | Y | 5.66 | 1.703 | 0.12841056 | 0.01683 | 1.28763889 | 0.245 |
| 223 | R | 10.76 | -0.113 | 0.13577389 | 0.027245 | 1.23858333 | 0.15695 |
| 224 | R | 10.76 | -0.306 | 0.10540444 | 0.01549 | 1.25701111 | 0.0678 |
| 225 | C | 5.07 | -1.039 | 0.09457556 | 0.01155 | 0.09658333 | 0.0505 |
| 226 | P | 6.3 | 2.552 | 0.11399944 | 0.014695 | 1.11941667 | 0.05175 |
| 227 | G | 5.97 | 0.522 | 0.10385667 | 0.01336 | 0.36389444 | 0.0501 |
| 228 | K | 9.74 | -0.114 | 0.129085 | 0.01451 | 0.97538889 | 0.12925 |
| 229 | R | 10.76 | 0.873 | 0.14559889 | 0.031005 | 1.48998333 | 0.35375 |
| 230 | M | 5.74 | -1.095 | 0.10897944 | 0.00878 | 0.00606111 | 0.0234 |
| 231 | Q | 5.65 | -0.167 | 0.14961278 | 0.0287 | 0.88933889 | 0.28795 |
| 232 | V | 5.96 | -0.987 | 0.13256667 | 0.02246 | 0.08195556 | 0.1404 |
| 233 | D | 2.77 | 0.271 | 0.18699556 | 0.04013 | 1.14017778 | 0.2092 |

[1]pI of residue amino acid at 25 °C.
[2]Consurf evolutionary conservation score of the residue.
[3]The average root-mean-square fluctuation (RMSF) across all frames and steps, and the ΔRMSF (maximum minus minimum across pH).
[4]The average solvent accessibility surface area (SASA) across all frames and steps, and the ΔSASA (maximum minus minimum across pH).

**Table S14. pH-dependent dynamics and evolutionary conservation of JAM-C residues.**

| Residue | AA | Residue pI[1] | Residue Conserv.[2] | Avg. RMSF[3] | ΔRMSF[3] | Avg. SASA[4] | ΔSASA[4] |
|---|---|---|---|---|---|---|---|
| 32 | V | 5.96 | -0.73 | 0.15238167 | 0.040935 | 0.5883 | 0.59125 |
| 33 | N | 5.41 | 0.019 | 0.14853222 | 0.0353 | 0.78888889 | 0.28905 |
| 34 | L | 5.98 | -0.664 | 0.10916833 | 0.027865 | 0.16058889 | 0.148 |
| 35 | K | 9.74 | 0.776 | 0.149245 | 0.031505 | 1.27495 | 0.12915 |
| 36 | S | 5.68 | -0.431 | 0.09753444 | 0.02756 | 0.21448333 | 0.1143 |
| 37 | S | 5.68 | 1.258 | 0.12149 | 0.04007 | 1.02638333 | 0.07985 |
| 38 | N | 5.41 | 0.106 | 0.11104667 | 0.02679 | 0.78813889 | 0.04355 |
| 39 | R | 10.76 | 2.316 | 0.13762222 | 0.033775 | 1.35476111 | 0.14785 |
| 40 | T | 5.6 | 1.479 | 0.11651111 | 0.020345 | 0.81445556 | 0.07305 |
| 41 | P | 6.3 | -0.79 | 0.09604611 | 0.019195 | 0.10311111 | 0.02725 |
| 42 | V | 5.96 | 0.712 | 0.12239944 | 0.0226 | 0.90961667 | 0.06085 |
| 43 | V | 5.96 | -0.725 | 0.11121667 | 0.02747 | 0.21483333 | 0.08445 |
| 44 | Q | 5.65 | 0.778 | 0.14549722 | 0.043235 | 0.79427778 | 0.2641 |
| 45 | E | 3.22 | -1.289 | 0.12993 | 0.04833 | 0.46055 | 0.2489 |
| 46 | F | 5.48 | -0.512 | 0.15773 | 0.053475 | 1.48767222 | 0.06265 |
| 47 | E | 3.22 | 1.276 | 0.14298 | 0.03259 | 0.87592778 | 0.13055 |
| 48 | S | 5.68 | 0.126 | 0.10325667 | 0.03267 | 0.50998889 | 0.12385 |
| 49 | V | 5.96 | -0.112 | 0.08759667 | 0.02084 | 0.13557778 | 0.02305 |
| 50 | E | 3.22 | -0.317 | 0.11123056 | 0.016725 | 0.85774444 | 0.1033 |
| 51 | L | 5.98 | -1.245 | 0.07430778 | 0.01716 | 0.00107778 | 0.00365 |
| 52 | S | 5.68 | -0.489 | 0.08012222 | 0.01807 | 0.14756667 | 0.1237 |
| 53 | C | 5.07 | -1.351 | 0.07702667 | 0.013385 | 0.00588889 | 0.0118 |
| 54 | I | 6.02 | 0.347 | 0.11072667 | 0.01526 | 0.80421111 | 0.12685 |
| 55 | I | 6.02 | -0.227 | 0.10934722 | 0.01877 | 0.35576111 | 0.25535 |
| 56 | T | 5.6 | 0.136 | 0.13636111 | 0.03076 | 0.71865 | 0.30635 |
| 57 | D | 2.77 | -0.406 | 0.16417556 | 0.02647 | 0.88234444 | 0.47545 |
| 58 | S | 5.68 | -0.524 | 0.15761722 | 0.058045 | 0.51121111 | 0.5636 |
| 59 | Q | 5.65 | -0.22 | 0.22276944 | 0.044575 | 1.73143889 | 0.3197 |
| 60 | T | 5.6 | -0.473 | 0.171085 | 0.028925 | 0.60655 | 0.42885 |
| 61 | S | 5.68 | 1.605 | 0.18193278 | 0.02358 | 1.05961111 | 0.15015 |
| 62 | D | 2.77 | 1.056 | 0.16444167 | 0.02774 | 0.88807778 | 0.2483 |
| 63 | P | 6.3 | -0.336 | 0.12535611 | 0.01826 | 0.16622778 | 0.12555 |
| 64 | R | 10.76 | -1.384 | 0.12480833 | 0.040145 | 0.59103889 | 0.5474 |
| 65 | I | 6.02 | -0.776 | 0.09349944 | 0.01576 | 0.13516667 | 0.092 |
| 66 | E | 3.22 | -1.342 | 0.09199278 | 0.009645 | 0.09039444 | 0.03365 |
| 67 | W | 5.89 | -1.295 | 0.05546278 | 0.00591 | 0.00155 | 0.00645 |
| 68 | K | 9.74 | -1.238 | 0.09211667 | 0.01964 | 0.32156667 | 0.27615 |
| 69 | K | 9.74 | -0.935 | 0.09186889 | 0.01893 | 0.09343889 | 0.10385 |
| 70 | I | 6.02 | 0.391 | 0.12096444 | 0.03132 | 0.43329444 | 0.20955 |
| 71 | Q | 5.65 | 0.981 | 0.16054667 | 0.03126 | 0.57053333 | 0.2559 |
| 72 | D | 2.77 | 0.984 | 0.18667 | 0.04845 | 1.22812778 | 0.3109 |
| 73 | E | 3.22 | 1.018 | 0.21444889 | 0.024785 | 1.75701667 | 0.0903 |
| 74 | Q | 5.65 | 0.577 | 0.18339167 | 0.02823 | 1.30648889 | 0.06765 |

| | | | | | | | |
|---|---|---|---|---|---|---|---|
| 75 | T | 5.6 | -0.433 | 0.12834056 | 0.02488 | 0.75446111 | 0.08245 |
| 76 | T | 5.6 | -0.206 | 0.11065722 | 0.02094 | 0.61425 | 0.0932 |
| 77 | Y | 5.66 | -0.366 | 0.11925944 | 0.017295 | 0.93657778 | 0.30815 |
| 78 | V | 5.96 | -1.144 | 0.07338167 | 0.01207 | 0.00595556 | 0.0084 |
| 79 | F | 5.48 | -0.118 | 0.07835389 | 0.011685 | 0.20944444 | 0.09805 |
| 80 | F | 5.48 | -0.023 | 0.12360167 | 0.015355 | 0.75427222 | 0.21335 |
| 81 | D | 2.77 | 0.72 | 0.13539167 | 0.03981 | 0.76692222 | 0.48865 |
| 82 | N | 5.41 | 0.178 | 0.14175611 | 0.017085 | 1.14822222 | 0.3528 |
| 83 | K | 9.74 | 1.219 | 0.16170444 | 0.02221 | 1.47176111 | 0.19905 |
| 84 | I | 6.02 | 0.12 | 0.095665 | 0.019235 | 0.33609444 | 0.156 |
| 85 | Q | 5.65 | -0.018 | 0.11862333 | 0.019595 | 0.59906667 | 0.23495 |
| 86 | G | 5.97 | 0.222 | 0.10816333 | 0.02468 | 0.58376667 | 0.0576 |
| 87 | D | 2.77 | 0.645 | 0.12299278 | 0.031 | 0.87505556 | 0.18605 |
| 88 | L | 5.98 | -0.705 | 0.09008889 | 0.015345 | 0.00767778 | 0.01165 |
| 89 | A | 6 | 0.833 | 0.10602556 | 0.01938 | 0.55467222 | 0.05435 |
| 90 | G | 5.97 | 1.297 | 0.11276333 | 0.02387 | 0.87434444 | 0.0421 |
| 91 | R | 10.76 | -1.31 | 0.09501722 | 0.024265 | 0.60745 | 0.0871 |
| 92 | A | 6 | -0.828 | 0.08073833 | 0.018645 | 0.08078889 | 0.0779 |
| 93 | E | 3.22 | 1.043 | 0.12690889 | 0.019305 | 1.04348889 | 0.10785 |
| 94 | I | 6.02 | 0.397 | 0.08764278 | 0.009755 | 0.38195 | 0.1476 |
| 95 | L | 5.98 | 0.896 | 0.11866167 | 0.02462 | 0.81112222 | 0.31715 |
| 96 | G | 5.97 | -0.199 | 0.11173889 | 0.01482 | 0.71916111 | 0.13325 |
| 97 | K | 9.74 | 1.173 | 0.15422667 | 0.057155 | 1.46917222 | 0.84975 |
| 98 | T | 5.6 | -0.517 | 0.08360778 | 0.020435 | 0.12276667 | 0.5383 |
| 99 | S | 5.68 | -0.498 | 0.06882889 | 0.011535 | 0.112 | 0.0497 |
| 100 | L | 5.98 | -0.741 | 0.06794778 | 0.00967 | 0.00017222 | 0.00065 |
| 101 | K | 9.74 | -0.128 | 0.09309611 | 0.013625 | 0.5474 | 0.1315 |
| 102 | I | 6.02 | -0.849 | 0.07854722 | 0.02159 | 0.00038889 | 0.00145 |
| 103 | W | 5.89 | 1.21 | 0.10345778 | 0.032255 | 1.03617778 | 0.10735 |
| 104 | N | 5.41 | -0.709 | 0.12638611 | 0.033695 | 0.81806111 | 0.04595 |
| 105 | V | 5.96 | -0.652 | 0.09944667 | 0.02678 | 0.00118333 | 0.0015 |
| 106 | T | 5.6 | -1.076 | 0.10873056 | 0.026755 | 0.43283889 | 0.1196 |
| 107 | R | 10.76 | -1.055 | 0.12541056 | 0.039615 | 0.80029444 | 0.56195 |
| 108 | R | 10.76 | 1.331 | 0.14035278 | 0.051455 | 1.62666667 | 0.2754 |
| 109 | D | 2.77 | -1.354 | 0.08995778 | 0.01686 | 0.02257778 | 0.0198 |
| 110 | S | 5.68 | -0.312 | 0.103155 | 0.02046 | 0.31601667 | 0.2279 |
| 111 | A | 6 | -0.974 | 0.09259611 | 0.01177 | 0.14018333 | 0.0423 |
| 112 | L | 5.98 | 1.84 | 0.11170222 | 0.021245 | 0.62011667 | 0.12365 |
| 113 | Y | 5.66 | -1.228 | 0.06945889 | 0.00672 | 0.0004 | 0.0015 |
| 114 | R | 10.76 | -0.602 | 0.10088611 | 0.026395 | 0.59685556 | 0.3361 |
| 115 | C | 5.07 | -1.353 | 0.06711722 | 0.007385 | 0.00073333 | 0.00125 |
| 116 | E | 3.22 | -1.023 | 0.10003389 | 0.02263 | 0.18226667 | 0.282 |
| 117 | V | 5.96 | -1.224 | 0.09805556 | 0.015235 | 0.00485556 | 0.01395 |
| 118 | V | 5.96 | -0.645 | 0.11819556 | 0.014515 | 0.10316111 | 0.21215 |
| 119 | A | 6 | -0.505 | 0.14153389 | 0.012635 | 0.19988333 | 0.09355 |

| | | | | | | | |
|---|---|---|---|---|---|---|---|
| 120 | R | 10.76 | 1.075 | 0.16923222 | 0.061185 | 1.24653333 | 0.32495 |
| 121 | N | 5.41 | 1.385 | 0.220055 | 0.021365 | 1.56833333 | 0.0909 |
| 122 | D | 2.77 | -0.701 | 0.19055333 | 0.038525 | 0.7232 | 0.33705 |
| 123 | R | 10.76 | 1.093 | 0.22043944 | 0.123785 | 1.93956667 | 1.0821 |
| 124 | K | 9.74 | 0.204 | 0.19021111 | 0.02654 | 1.52600556 | 0.1426 |
| 125 | E | 3.22 | 1.573 | 0.15146611 | 0.03468 | 0.72071111 | 0.74565 |
| 126 | I | 6.02 | 0.594 | 0.132905 | 0.02619 | 0.72871111 | 0.44665 |
| 127 | D | 2.77 | 0.353 | 0.11650167 | 0.027155 | 0.5538 | 0.376 |
| 128 | E | 3.22 | -1.067 | 0.12420722 | 0.01727 | 0.77031667 | 0.22975 |
| 129 | I | 6.02 | 0.173 | 0.09551389 | 0.01142 | 0.11756111 | 0.1082 |
| 130 | V | 5.96 | 1.477 | 0.10085778 | 0.01461 | 0.50656667 | 0.15955 |
| 131 | I | 6.02 | -0.637 | 0.08296 | 0.00998 | 0.00069444 | 0.00235 |
| 132 | E | 3.22 | 0.986 | 0.12737222 | 0.017875 | 0.85612778 | 0.07725 |
| 133 | L | 5.98 | -1.309 | 0.09057167 | 0.016635 | 0.03122222 | 0.01365 |
| 134 | T | 5.6 | -0.356 | 0.11260389 | 0.024945 | 0.36887778 | 0.12535 |
| 135 | V | 5.96 | -1.29 | 0.11007778 | 0.024495 | 0.01103333 | 0.0184 |
| 136 | Q | 5.65 | -0.695 | 0.14205444 | 0.02729 | 0.2598 | 0.0895 |
| 137 | V | 5.96 | -1.392 | 0.12489889 | 0.02727 | 0.07708889 | 0.0589 |
| 138 | K | 9.74 | -0.92 | 0.14633278 | 0.02963 | 1.14783889 | 0.34995 |
| 139 | P | 6.3 | -1.369 | 0.10702278 | 0.027115 | 0.03886111 | 0.0225 |
| 140 | V | 5.96 | -0.737 | 0.12900833 | 0.035965 | 0.79703333 | 0.2532 |
| 141 | T | 5.6 | 0.451 | 0.126575 | 0.027275 | 1.11596111 | 0.29985 |
| 142 | P | 6.3 | -1.369 | 0.09401722 | 0.013645 | 0.05931111 | 0.0321 |
| 143 | V | 5.96 | 0.522 | 0.11662389 | 0.013715 | 0.70198333 | 0.1418 |
| 144 | C | 5.07 | -1.048 | 0.08879778 | 0.012425 | 0.08910556 | 0.05855 |
| 145 | R | 10.76 | 0.393 | 0.13850889 | 0.024005 | 1.54761667 | 0.25745 |
| 146 | V | 5.96 | -0.757 | 0.08694611 | 0.01722 | 0.17622222 | 0.05925 |
| 147 | P | 6.3 | -1.261 | 0.10152944 | 0.027455 | 0.44331667 | 0.0739 |
| 148 | K | 9.74 | 0.274 | 0.16439944 | 0.05981 | 1.71556111 | 0.27335 |
| 149 | A | 6 | -0.974 | 0.12909 | 0.047715 | 0.46372222 | 0.13345 |
| 150 | V | 5.96 | -1.011 | 0.12234556 | 0.030265 | 0.10721111 | 0.0342 |
| 151 | P | 6.3 | -0.793 | 0.14171167 | 0.045485 | 0.65870556 | 0.1415 |
| 152 | V | 5.96 | -0.816 | 0.14766444 | 0.038255 | 0.74506111 | 0.0617 |
| 153 | G | 5.97 | -1.24 | 0.14002222 | 0.03801 | 0.41479444 | 0.0605 |
| 154 | K | 9.74 | -0.081 | 0.17050944 | 0.03725 | 1.37987778 | 0.07015 |
| 155 | M | 5.74 | 1.216 | 0.17010222 | 0.03698 | 1.27945 | 0.0545 |
| 156 | A | 6 | -0.877 | 0.10344778 | 0.02094 | 0.05093889 | 0.02425 |
| 157 | T | 5.6 | 0.189 | 0.11170944 | 0.01897 | 0.60241667 | 0.1294 |
| 158 | L | 5.98 | -1.246 | 0.07874 | 0.013065 | 0.00025 | 0.0005 |
| 159 | H | 7.95 | 0.389 | 0.10872778 | 0.013785 | 0.73559444 | 0.16205 |
| 160 | C | 5.07 | -1.353 | 0.08507722 | 0.009145 | 0.0302 | 0.03295 |
| 161 | Q | 5.65 | 1.075 | 0.14504722 | 0.01404 | 0.95874444 | 0.14475 |
| 162 | E | 3.22 | -1.038 | 0.11049 | 0.02188 | 0.18523333 | 0.1129 |
| 163 | S | 5.68 | 1.331 | 0.14088389 | 0.043695 | 0.94656667 | 0.1509 |
| 164 | E | 3.22 | -0.208 | 0.14554778 | 0.045455 | 0.65898333 | 0.32095 |

| | | | | | | | |
|---|---|---|---|---|---|---|---|
| 165 | G | 5.97 | -1.078 | 0.11875111 | 0.02879 | 0.05277778 | 0.11665 |
| 166 | H | 7.95 | -0.372 | 0.14000611 | 0.05127 | 0.70720556 | 0.59245 |
| 167 | P | 6.3 | -1.369 | 0.12856944 | 0.03133 | 0.55533889 | 0.0893 |
| 168 | R | 10.76 | 0.334 | 0.17168833 | 0.057315 | 1.6913 | 0.30215 |
| 169 | P | 6.3 | -0.317 | 0.10968722 | 0.021285 | 0.07062222 | 0.04375 |
| 170 | H | 7.95 | 0.056 | 0.13569111 | 0.030425 | 1.08017778 | 0.21355 |
| 171 | Y | 5.66 | -1.093 | 0.08900056 | 0.0126 | 0.16378333 | 0.073 |
| 172 | S | 5.68 | 0.603 | 0.07510389 | 0.014125 | 0.43262778 | 0.0966 |
| 173 | W | 5.89 | -1.179 | 0.061225 | 0.00767 | 0.01520556 | 0.0206 |
| 174 | Y | 5.66 | 0.005 | 0.07808056 | 0.01607 | 0.53171667 | 0.0549 |
| 175 | R | 10.76 | -0.512 | 0.08855333 | 0.0178 | 0.21978333 | 0.27775 |
| 176 | N | 5.41 | -0.672 | 0.11489444 | 0.01946 | 0.80246111 | 0.1605 |
| 177 | D | 2.77 | 1.21 | 0.11979 | 0.029235 | 1.03341111 | 0.0561 |
| 178 | V | 5.96 | 0.753 | 0.103065 | 0.011525 | 0.57786111 | 0.3406 |
| 179 | P | 6.3 | 1.198 | 0.096595 | 0.013485 | 0.77492222 | 0.09945 |
| 180 | L | 5.98 | -0.635 | 0.09407778 | 0.014235 | 0.09995556 | 0.0518 |
| 181 | P | 6.3 | -0.733 | 0.11080556 | 0.018925 | 0.3477 | 0.18805 |
| 182 | T | 5.6 | 3.162 | 0.14578444 | 0.0418 | 0.89092222 | 0.1961 |
| 183 | D | 2.77 | 0.053 | 0.17316556 | 0.053415 | 0.73231111 | 0.48495 |
| 184 | S | 5.68 | -0.518 | 0.15837278 | 0.069825 | 0.26077778 | 0.738 |
| 185 | R | 10.76 | 0.38 | 0.21966778 | 0.06348 | 1.75253889 | 0.5848 |
| 186 | A | 6 | 0.787 | 0.19596889 | 0.02933 | 0.79601667 | 0.23145 |
| 187 | N | 5.41 | 0.139 | 0.173795 | 0.04373 | 0.47746667 | 0.3306 |
| 188 | P | 6.3 | 2.614 | 0.19571778 | 0.028305 | 1.07455556 | 0.16595 |
| 189 | R | 10.76 | 0.335 | 0.20613167 | 0.067955 | 1.78156667 | 0.5236 |
| 190 | F | 5.48 | 0.283 | 0.13651611 | 0.022815 | 0.22906111 | 0.31095 |
| 191 | R | 10.76 | 2.047 | 0.21763833 | 0.053195 | 1.58932222 | 0.71245 |
| 192 | N | 5.41 | -1.195 | 0.18313556 | 0.044745 | 1.27802778 | 0.3522 |
| 193 | S | 5.68 | -0.222 | 0.12811222 | 0.026675 | 0.13511111 | 0.12405 |
| 194 | S | 5.68 | -0.907 | 0.12951056 | 0.03262 | 0.45800556 | 0.2194 |
| 195 | F | 5.48 | -1.09 | 0.10500444 | 0.020885 | 0.11843889 | 0.14255 |
| 196 | H | 7.95 | 0.334 | 0.14960889 | 0.026245 | 1.10740556 | 0.24885 |
| 197 | L | 5.98 | 0.96 | 0.11452611 | 0.026695 | 0.18997778 | 0.1583 |
| 198 | N | 5.41 | -0.463 | 0.13266833 | 0.027645 | 0.66713333 | 0.11675 |
| 199 | S | 5.68 | 3.072 | 0.13083389 | 0.04451 | 0.4274 | 0.40665 |
| 200 | E | 3.22 | 1.643 | 0.18038722 | 0.047895 | 1.32795556 | 0.52975 |
| 201 | T | 5.6 | -0.221 | 0.13282056 | 0.03326 | 0.7964 | 0.06895 |
| 202 | G | 5.97 | -1.301 | 0.09515056 | 0.02245 | 0.00098889 | 0.0026 |
| 203 | T | 5.6 | 0.058 | 0.103965 | 0.01477 | 0.29866667 | 0.1482 |
| 204 | L | 5.98 | -1.246 | 0.07965111 | 0.013085 | 0.00135 | 0.00385 |
| 205 | V | 5.96 | 1.056 | 0.10532778 | 0.021745 | 0.39061667 | 0.1197 |
| 206 | F | 5.48 | -1.031 | 0.09573444 | 0.020715 | 0.0114 | 0.0199 |
| 207 | T | 5.6 | 0.275 | 0.142555 | 0.03751 | 0.75131667 | 0.1457 |
| 208 | A | 6 | 1.039 | 0.12772889 | 0.029355 | 0.37081111 | 0.10405 |
| 209 | V | 5.96 | -0.62 | 0.11443722 | 0.026765 | 0.01077778 | 0.02335 |

| | | | | | | | |
|---|---|---|---|---|---|---|---|
| 210 | H | 7.95 | 0.426 | 0.14682222 | 0.048035 | 0.79716667 | 0.40175 |
| 211 | K | 9.74 | -0.439 | 0.16359333 | 0.026745 | 1.35587778 | 0.3112 |
| 212 | D | 2.77 | 1.378 | 0.15070722 | 0.02121 | 1.11762222 | 0.2921 |
| 213 | D | 2.77 | -1.311 | 0.09886833 | 0.012555 | 0.04207222 | 0.0973 |
| 214 | S | 5.68 | -0.046 | 0.10819667 | 0.01623 | 0.40345 | 0.0468 |
| 215 | G | 5.97 | -0.988 | 0.09403833 | 0.012765 | 0.12661111 | 0.03465 |
| 216 | Q | 5.65 | 0.412 | 0.12982944 | 0.020335 | 0.71333889 | 0.069 |
| 217 | Y | 5.66 | -1.157 | 0.06613944 | 0.008725 | 0.00209444 | 0.0064 |
| 218 | Y | 5.66 | 1.448 | 0.07623611 | 0.019735 | 0.17310556 | 0.09205 |
| 219 | C | 5.07 | -1.353 | 0.06292889 | 0.01281 | 0.00042222 | 0.001 |
| 220 | I | 6.02 | -0.134 | 0.07973833 | 0.02102 | 0.33608333 | 0.12775 |
| 221 | A | 6 | -1.258 | 0.08188611 | 0.02135 | 0.00030556 | 0.001 |
| 222 | S | 5.68 | 0.976 | 0.09423833 | 0.02764 | 0.39621111 | 0.1259 |
| 223 | N | 5.41 | -1.367 | 0.10709333 | 0.030415 | 0.2104 | 0.05 |
| 224 | D | 2.77 | 1.352 | 0.15001167 | 0.035175 | 1.27873333 | 0.07045 |
| 225 | A | 6 | -0.63 | 0.12448833 | 0.03598 | 0.21572778 | 0.1301 |
| 226 | G | 5.97 | -1.303 | 0.10608167 | 0.036175 | 0.21102778 | 0.0326 |
| 227 | S | 5.68 | 1.7 | 0.09985111 | 0.02693 | 0.77652222 | 0.05265 |
| 228 | A | 6 | -0.452 | 0.087975 | 0.02636 | 0.18739444 | 0.06805 |
| 229 | R | 10.76 | 0.68 | 0.12434556 | 0.021725 | 1.4304 | 0.2851 |
| 230 | C | 5.07 | -1.175 | 0.08447333 | 0.015855 | 0.05321667 | 0.03895 |
| 231 | E | 3.22 | 2.214 | 0.14370111 | 0.02265 | 1.41147222 | 0.11735 |
| 232 | E | 3.22 | 0.929 | 0.12637278 | 0.02291 | 0.90272778 | 0.0783 |
| 233 | Q | 5.65 | 0.024 | 0.09881167 | 0.01789 | 0.54691111 | 0.125 |
| 234 | E | 3.22 | 1.743 | 0.14102944 | 0.025395 | 0.96234444 | 0.21975 |
| 235 | M | 5.74 | -1.097 | 0.10392778 | 0.014765 | 0.00811111 | 0.0114 |
| 236 | E | 3.22 | -0.686 | 0.14819944 | 0.03239 | 0.86967222 | 0.19415 |
| 237 | V | 5.96 | -0.974 | 0.124605 | 0.030855 | 0.09073889 | 0.05595 |
| 238 | Y | 5.66 | -0.058 | 0.18207111 | 0.047295 | 1.67244444 | 0.17095 |

[1]pI of residue amino acid at 25 °C.
[2]Consurf evolutionary conservation score of the residue.
[3]The average root-mean-square fluctuation (RMSF) across all frames and steps, and the ΔRMSF (maximum minus minimum across pH).
[4]The average solvent accessibility surface area (SASA) across all frames and steps, and the ΔSASA (maximum minus minimum across pH).

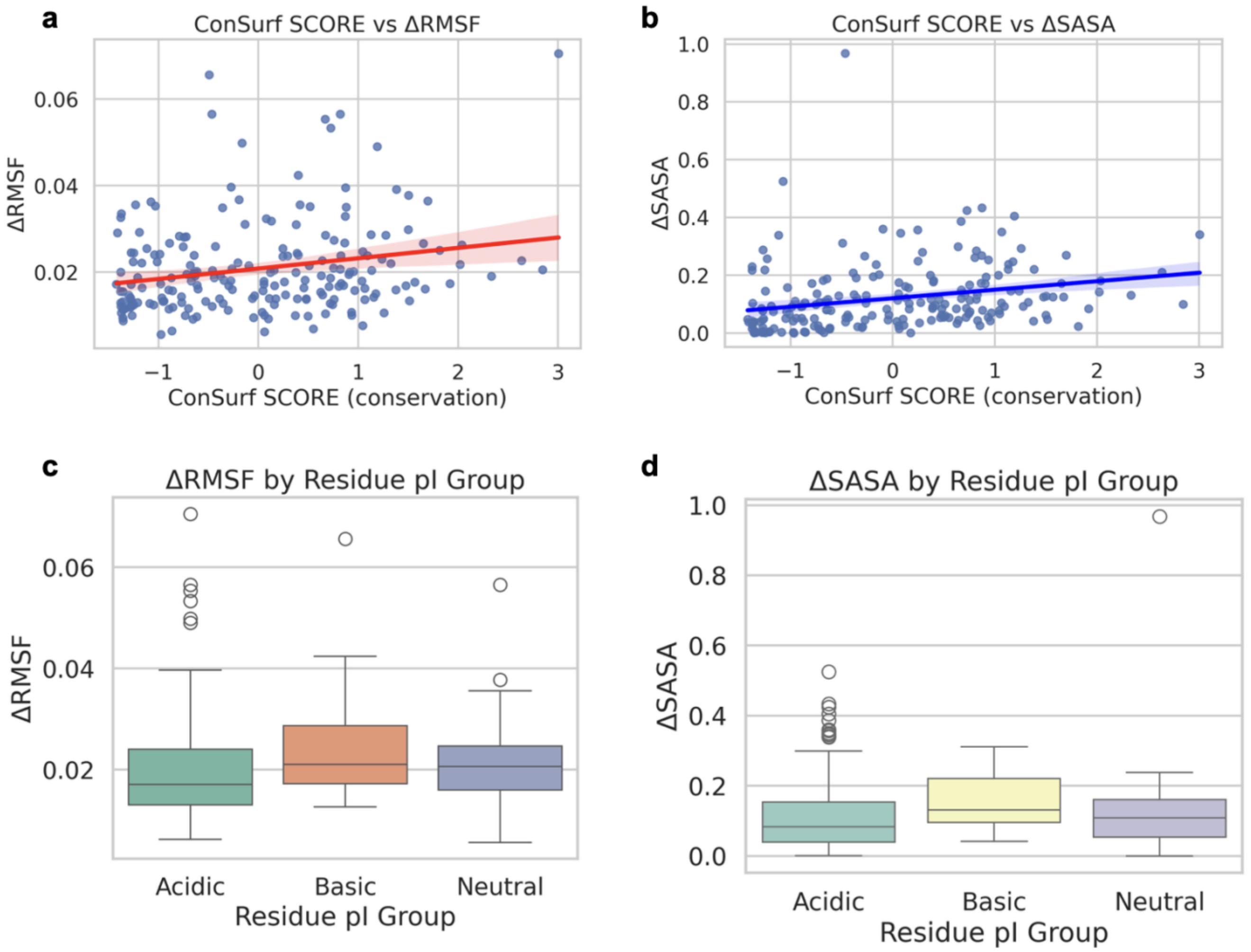

**Figure S7. Interplay between protein dynamics with residue conservation and pI of JAM-A.**
Per-residue root-mean-square fluctuations (RMSF) and solvent-accessible surface area (SASA) vs. evolutionary conservation **(a, b)** and residue pI **(c, d)**.

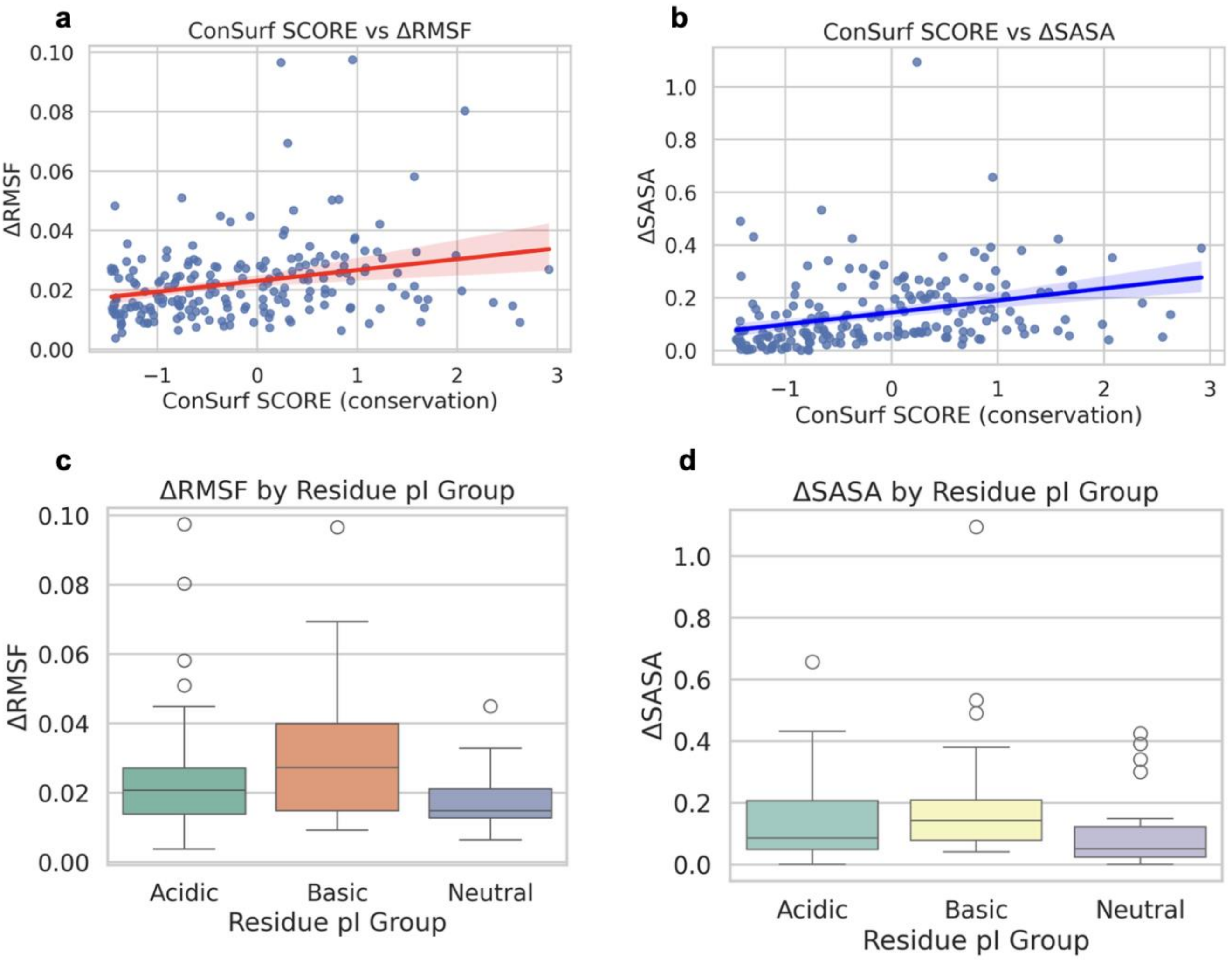

**Figure S8. Interplay between protein dynamics with residue conservation and pI of JAM-B.**
Per-residue root-mean-square fluctuations (RMSF) and solvent-accessible surface area (SASA) vs. evolutionary conservation **(a, b)** and residue pI **(c, d)**.

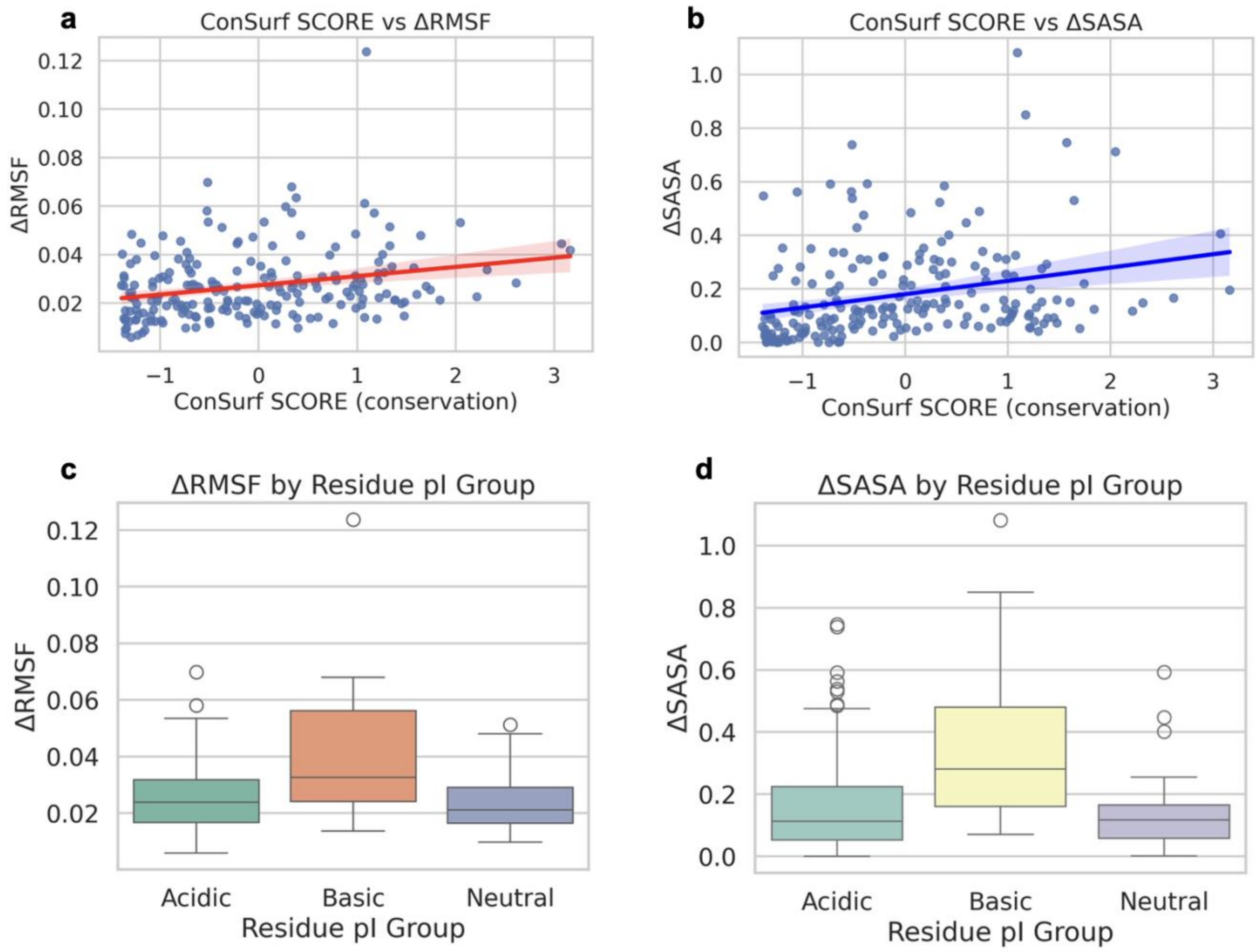

**Figure S9. Interplay between protein dynamics with residue conservation and pI of JAM-C.**
Per-residue root-mean-square fluctuations (RMSF) and solvent-accessible surface area (SASA) vs. evolutionary conservation **(a, b)** and residue pI **(c, d)**.
